# Nuclear Localization Signals Enable the Cellular Delivery of an Anti-CRISPR Protein to Control Genome Editing

**DOI:** 10.1101/2025.10.28.685205

**Authors:** Axel O. Vera, Franklin J. Avilés-Vázquez, Taekjip Ha, Amit Choudhary, Ronald T. Raines

## Abstract

Precise regulation of Cas9 activity is essential to minimize off-target effects, mosaicism, chromosomal alterations, immunogenicity, and genotoxicity in genome editing. Although type II anti-CRISPR proteins (Acrs) can inhibit and regulate Cas9, their size and anionic charge generally prevent them from crossing the cell membrane. Existing Acr delivery methods employing vectors or electroporation are either slow and persistent or require external equipment, limiting their therapeutic utility. To address these challenges, we developed a cell-permeable Acr (6×NLS-Acr), which uses nuclear localization signals (NLSs) to cross the cell membrane. We conjugated 6×NLS-Acr to a fluorescent dye to elucidate its cellular entry mechanism and directly visualized its binding to a fluorescent Cas9·gRNA complex to study its inhibitory mechanism. 6×NLS-Acr (IC_50_ = 0.47 µM) directly transduces human cells, including immortalized cell lines, embryonic stem cells, and 3D cell cultures, within 5 min, inhibiting up to 99% of Cas9 activity and increasing genome-editing specificity by nearly 100%. We further compared 6×NLS-Acr with our anthrax-derived Acr delivery platform. Our results demonstrate that 6×NLS-Acr is the most efficacious cell-permeable CRISPR-Cas inhibitor, significantly enhancing the precision and therapeutic potential of CRISPR-based genome editing.

## INTRODUCTION

Clustered regularly interspaced short palindromic repeats (CRISPR)/CRISPR-associated (Cas) systems use RNA-guided endonucleases (*e.g.*, Cas9) to cleave foreign DNA and defend bacteria against phage infections.^1,2^ Due to its ease of programmability, Cas9 has been adapted as a powerful genome-editing tool in human cells.^3,4^ Cas9-based technologies are advancing the treatment of genetic disorders,^5-7^ but their broader application is hindered by safety concerns, including unintended DNA modifications (*i*.*e*., off-target effects),^8-11^ genetic mosaicism,^12^ chromosomal abnormalities,^13^ immunogenicity,^14^ and genotoxicity.^15,16^

Type II anti-CRISPR proteins (Acrs) can address these limitations by inhibiting Cas9 in cells. Evolved by phages to disable bacterial CRISPR immunity, Acrs bind Cas9 with high specificity and affinity.^17,18^ They inhibit Cas9 at one or more of its three interference stages: gRNA loading, DNA binding, or DNA cleavage.^19^ For instance, AcrIIA4, an anionic Acr, predominantly binds to the PAM-interacting domain of the Cas9·gRNA ribonucleoprotein (RNP) complex, thereby preventing DNA binding.^20-22^ Acrs have been used to regulate Cas9 activity,^23,24^ mitigate its toxicity,^25^ and increase its efficiency^26^ and specificity.^22,27-31^

Due to their inability to cross cell membranes, Acrs require delivery vehicles to enter cells. Viral vectors (*e.g.*, adenovirus,^25^ AAVs,^27,28,32^ and lentiviruses,^33,34^) and lipid nanoparticles^35^ have been used to deliver Acr-encoding DNA and mRNA, *ex vivo* and in vivo, validating the therapeutic potential of Acrs. Viral vectors, however, require expression, are immunogenic, and can integrate into the host genome.^36^ In contrast, mRNA delivery is transient and avoids genomic integration but still relies on cellular translation, which takes time and can be influenced by cell type and stress. Consequently, DNA- and mRNA-based Acr delivery methods are slow, delaying Cas9 inhibition by several hours.^37^

Delivering Acrs directly as proteins via nucleofection could overcome this delay, enabling immediate Cas9 inhibition.^22^ As a non-viral method, nucleofection avoids host genome integration, prolonged Acr expression, and vector packaging constraints. Yet, nucleofection requires specialized equipment and electrical pulses, limiting its use to *ex vivo* applications. Additionally, delivering Cas9 and Acrs sequentially via nucleofection can increase cellular stress and toxicity.^38^

As an alternative to nucleofection, we recently established LF_N_-Acr/PA—a protein-based delivery system for introducing Acrs into human cells.^31^ LF_N_-Acr/PA consists of two proteins derived from anthrax toxin. The first component, LF_N_-Acr, is a fusion between the nontoxic N-terminal domain of lethal factor from *Bacillus anthracis* and AcrIIA4 from *Listeria monocytogenes* prophages. The second component, protective antigen (PA) from Bacillus anthracis, binds the human receptors ANTRX1 and ANTRX2 and, after protease-mediated activation, forms a pH-triggered endosomal pore that allows LF_N_-Acr to enter the cell. LF_N_-Acr/PA is the most potent known cell-permeable CRISPR-Cas inhibition system, inhibiting Cas9 at concentrations as low as 2.5 pM and increasing its specificity by up to 40%. The LF_N_-Acr/PA system, however, is limited by its reliance on two separate protein components—LF_N_-Acr and PA—for Acr delivery. Additionally, PA pore formation requires binding to either the ANTRX1 or ANTRX2 receptor on the cell surface, followed by proteolytic cleavage by a cell-surface furin protease.^39-41^ As a result, PA-mediated Acr delivery requires the coordinated action of at least four distinct proteins, adding further complexity to the already intricate process of Cas9 and gRNA delivery.

Here, we describe the first peptide-based delivery system for introducing Acrs into human cells. We show that 6×NLS-Acr, an Acr fused to six nuclear localization signals (NLSs), readily enters cells via a novel direct protein transduction mechanism, inhibits up to 99% of Cas9 activity, and increases genome-editing specificity by 90%. We compare the delivery of 6×NLS-Acr to the PA-mediated delivery of LF_N_-Acr. We show that 6×NLS-Acr enters cells less effectively than LF_N_-Acr/PA at low concentrations, but more effectively above 1 µM. Unlike LF_N_-Acr/PA, 6×NLS-Acr effectively enters cells after Cas9 lipofection. Thus, 6×NLS-Acr is a more efficacious inhibitor, owing to its simpler, one-component, receptor-independent delivery mechanism.

## RESULTS

### Design of the Cellular Delivery System

NLSs are short peptide sequences that transport proteins from the cytosol to the nucleus with the assistance of nuclear import proteins.^42^ NLSs (*e*.*g*., PKKKRKV, derived from the SV40 antigen) can also function as cell-penetrating peptides (CPPs).^43,44^ CPPs can internalize proteins into cells via a combination of mechanisms that include endosomal escape and direct penetration.^45^ Tandem arrays (>2) of SV40 NLSs have been used to deliver zinc finger nucleases^46^ and Cas proteins (*e*.*g*., Cas9 inserted between four N-terminal NLSs and two C-terminal NLSs) into cells.^47-51^

Thus, we hypothesized that NLSs could mediate the cellular delivery of Acrs (Figure 1A). A cell-permeable NLS-Acr fusion would have the advantage of being a non-viral, single-component entity. It would enter cells rapidly (within 1 h) if it behaves like other NLS fusion proteins.^44^ Cell-permeable NLS fusion proteins have been shown to penetrate tissues in vivo;^47,50,51^ hence, NLS-Acrs could also be applicable in vivo.

**Figure 1.**
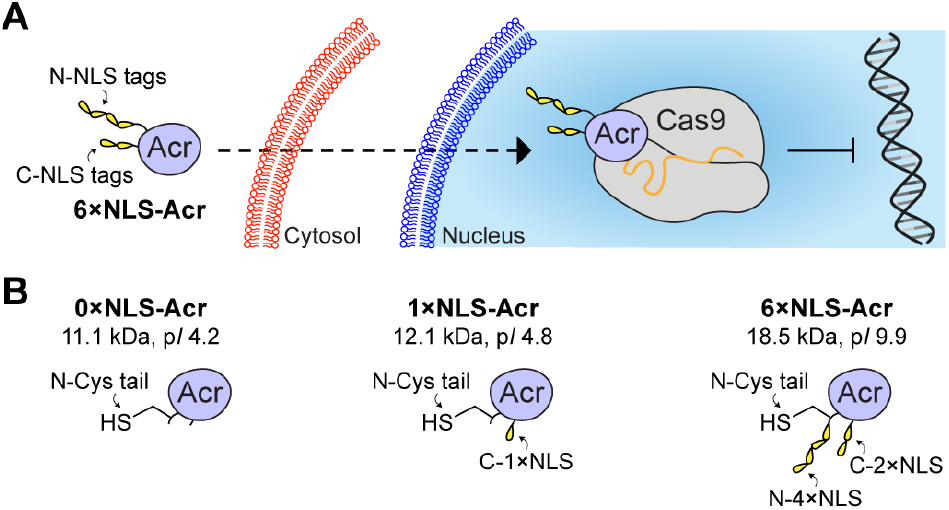
Nuclear localization signals (NLSs) to deliver Acrs into cells. (A) Cationic NLSs deliver Acrs into the cytosol and nucleus of a cell. (B) Cartoon representations of 0×NLS-Acr, 1×NLS-Acr, and 6×NLS-Acr. Acrs were recombinantly produced and purified in *Escherichia coli*. The Acr sequence corresponds to AcrIIA4 from *Listeria monocytogenes* prophages, engineered with an N-terminal G_4_CG_4_S sequence (*i.e.*, an N-Cys tail) to facilitate site-specific bioconjugation. Acrs are labeled based on the number (0, 1, or 6×) of SV40 NLSs (sequence: PKKKRKV) inserted at either terminus. The molar mass and isoelectric point (p*I*) of each Acr variant were determined using the Expasy ProtParam tool.

We chose AcrIIA4 as a model Acr because it potently inhibits *Streptococcus pyogenes* Cas9 (SpCas9) and tolerates modifications at its termini, including the insertion of dual cysteine residues.^20,52^ We first introduced a cysteine residue in the N-terminus of AcrIIA4 to allow site-specific bioconjugation. We then produced and purified Cys-AcrIIA4 without NLSs (0×NLS-Acr), Cys-AcrIIA4 with a C-terminal NLS (1×NLS-Acr), and Cys-AcrIIA4 inserted between four N-terminal NLSs and two C-terminal NLSs (6×NLS-Acr) using an MBP-TEV system in *Escherichia coli* and assessed purity by SDS–PAGE and LC–MS (Figure 1B and S1–S3). 0×NLS-Acr and 1×NLS-Acr are anionic proteins (p*I* 4.2–4.8), whereas 6×NLS-Acr is a cationic protein (p*I* 9.9). Their masses range from 11.1 to 18.5 kDa.

### 6×NLS-Acr inhibits Cas9 in a Dose- and Time-Dependent Manner with High Specificity

We tested Acr delivery by monitoring Cas9 nuclease activity using the *GFP*-disruption assay. We delivered Cas9 RNP targeting a *GFP* gene via nucleofection into a cell line stably expressing a GFP–PEST fusion (U2OS-EGFP.PEST).^53^ We then treated the seeded cells with Acrs at time points ranging from 0 to 24 h and imaged at 48 h. Cas9-mediated cleavage of *GFP* reduces GFP fluorescence. Conversely, Acr-mediated inhibition of Cas9 conserves GFP fluorescence (Figure 2A). 6×NLS-Acr (0.13–4 µM) inhibited Cas9 activity (always normalized to 100% without Acrs) from 100 to 4.9% (Figures 2B, 2C, and S4–S9). 0×NLS-Acr (4 µM) and 1×NLS-Acr (4 µM) did not inhibit Cas9, which shows that multiple NLS tags are needed to mediate Acr delivery. 6×NLS-Acr (2–4 µM) inhibited Cas9 but not Cas12a (Figures S10 and S11), validating the target specificity of 6×NLS-Acr.

**Figure 2.**
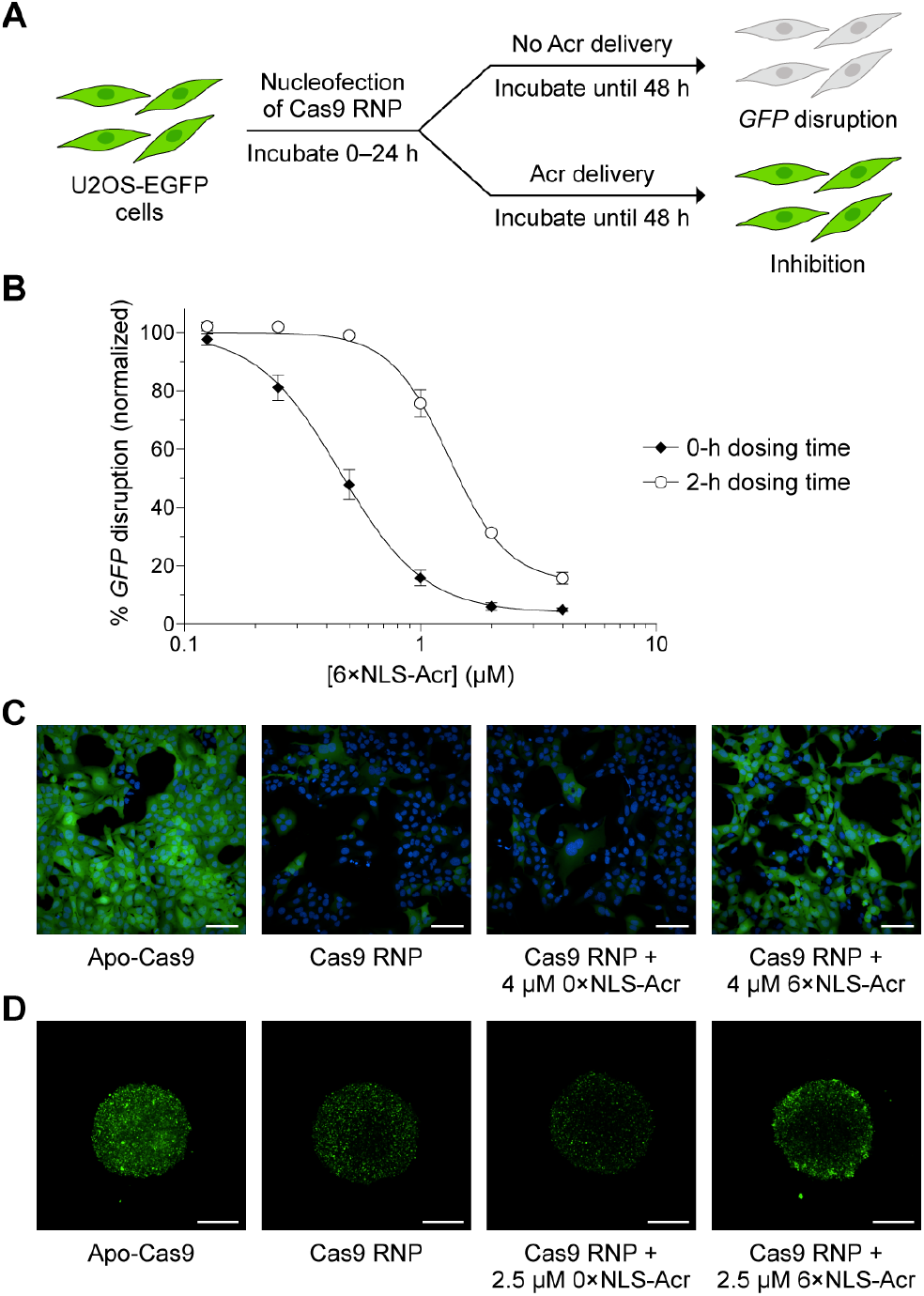
Effect of Acr delivery on *GFP* disruption. (A) A cartoon illustrating the experimental setup for the *GFP*-disruption assay. Acr delivery prevents *GFP* disruption. (B) Dose-dependent inhibition of Cas9 by 6×NLS-Acr in the *GFP*-disruption assay. SpCas9-NLS RNP (20 pmol) targeting *GFP* was delivered via nucleofection into U2OS-EGFP.PEST cells. The cells were seeded, incubated at 37 °C, and treated with 6×NLS-Acr (0.13, 0.25, 0.5, 1, 2, and 4 µM) at 0 h or 2 h. After 48 h, live cells were stained with Hoechst 33342 and imaged using high-throughput confocal microscopy. The values were normalized to Cas9 RNP (100%, not shown) and are the mean ± standard deviation (SD) of three independent replicates. Some error bars are too small to visualize. (C) Representative fluorescence images (20× magnification) from the *GFP*-disruption assay described in panel B (2-h dosing time). Controls include nucleofection of Apo-Cas9, nucleofection of Cas9 RNP, and nucleofection of Cas9 RNP followed by treatment with 0×NLS-Acr (4 µM) at 2 h. The cell nucleus was stained with Hoechst 33342 (blue). Scale bars, 100 µm. (D) Representative fluorescence images (5× magnification) from a *GFP*-disruption assay adapted for 3D cell cultures. SpCas9-NLS RNP (20 pmol), targeting *GFP*, was delivered via nucleofection into U2OS-EGFP.PEST cells. The cells were seeded in a spheroid plate, incubated at 37 °C, and treated with 6×NLS-Acr (2.5 µM) at 7 h. After 48 h, live cells were imaged via high-throughput confocal microscopy. Controls include nucleofection of Apo-Cas9, nucleofection of Cas9 RNP, and nucleofection of Cas9 RNP followed by treatment with 0×NLS-Acr (2.5 µM) at 7 h. Scale bars, 500 µm.

### 6×NLS-Acr inhibits Cas9 in 3D Cell Cultures

We evaluated 6×NLS-Acr delivery using a *GFP*-disruption assay adapted for three-dimensional (3D) cell cultures.^31^ We delivered Cas9 RNP into U2OS-EGFP.PEST cells via nucleofection and seeded them in a spheroid plate. The cells were then treated with Acrs at 7 and 13 h. Fluorescence imaging showed loosely aggregated spheroids at 7 and 13 h with an average diameter of 2.53 and 2.10 mm, respectively (Figures S12 and S13), and well-defined spheroids at 48 h with an average diameter of 1.25 mm (Figure S14). 6×NLS-Acr (2.5 µM) reduced Cas9 activity from 100 to 32% in the outer spheroid region (0.20 mm in depth) and from 100 to 61% when assessing the mean fluorescence of the entire spheroid region (Figures 2D, S14, and S15). 6×NLS-Acr did not reach the inner spheroid region. 0×NLS-Acr (2.5 µM) did not inhibit Cas9 in 3D cell cultures.

### 6×NLS-Acr Inhibits Cas9 in Human Embryonic Stem Cells

We further validated 6×NLS-Acr delivery by monitoring Cas9 nuclease activity using the HiBiT-knock-in assay. We delivered Cas9 RNP targeting *GAPDH* and a HiBiT single-stranded oligodeoxynucleotide (ssODN) donor via nucleofection into human embryonic stem cells (HUES 8). We then treated the seeded cells with Acrs at time points ranging from 0 to 16 h, and quantified luminescence (Figure 3A). Cas9-mediated cleavage of *GAPDH* and insertion of the HiBiT ssODN via homology-directed repair (HDR) leads to the formation of NanoLuc in the presence of LgBiT and furimazine. Conversely, inhibition of Cas9 with 6×NLS-Acr prevents NanoLuc formation. 6×NLS-Acr (0.13–4 µM) inhibited Cas9 activity from 100 to 1.4% (Figures 3B and 3C). 0×NLS-Acr (4 µM) did not inhibit Cas9.

**Figure 3.**
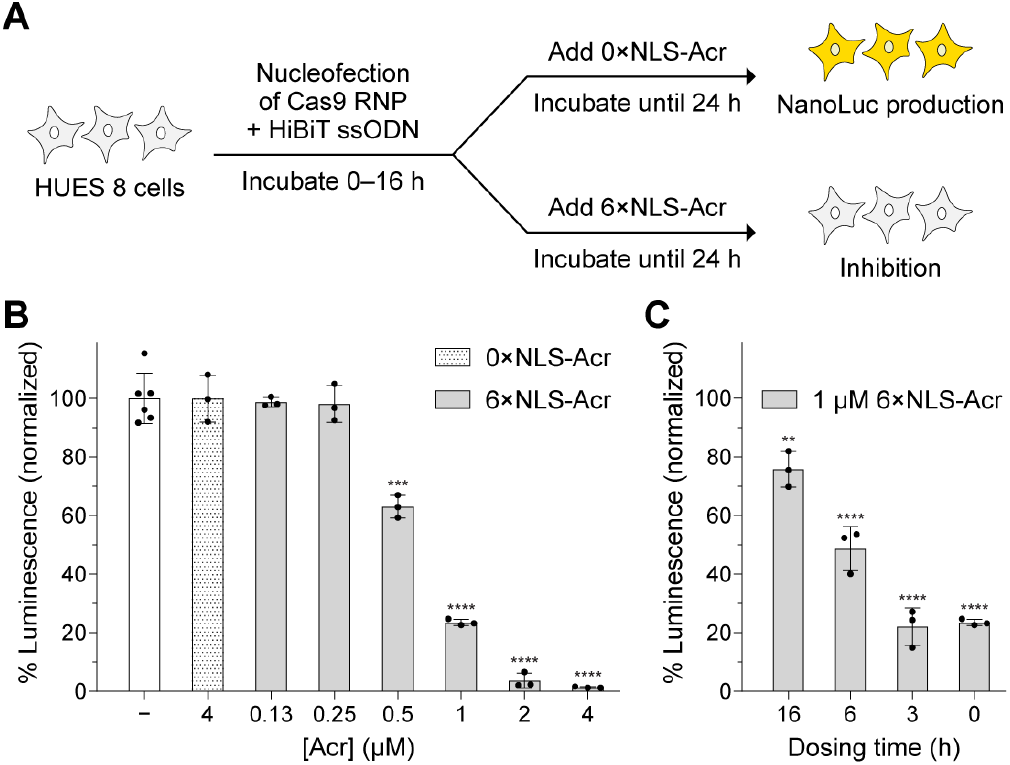
Effect of Acr delivery on HiBiT knock-in. (A) A cartoon illustrating the experimental setup for the HiBiT-knock-in assay. Acr delivery prevents Cas9-mediated knock-in and NanoLuc formation. (B, C) Dose- and time-dependent inhibition of Cas9 by 6×NLS-Acr in the HiBiT*-*knock-in assay. SpCas9-NLS RNP (20 pmol) targeting *GAPDH* and the HiBiT ssODN (80 pmol) were co-delivered via nucleofection into HUES 8 cells. The cells were seeded, incubated at 37 °C, and treated with 6×NLS-Acr (0.13–4 µM) at 0 h (B) or treated with 6×NLS-Acr (1 µM), varying the dosing time from 0–16 h (C). After 24 h, the cells were lysed and combined with LgBiT and furimazine, and their luminescence was quantified. Controls include co-nucleofection of Cas9 RNP and ssODN (white bar) and co-nucleofection of Cas9 RNP and ssODN followed by treatment with 4 µM 0×NLS-Acr at 0 h (panel B). The values were normalized to Cas9 RNP with ssODN and are the mean ± SD of three independent replicates. The significance of 6×NLS-Acr delivery was determined using an unpaired, two-tailed *t*-test versus Cas9 RNP with ssODN, where *, **, ***, and **** refer to *P* ≤ 0.05, *P* ≤ 0.01, *P* ≤ 0.001, and *P* ≤ 0.0001, respectively.

We also used the HiBiT knock-in assay to confirm the delivery of 6×NLS-Acr into HEK293T cells (Figure S16). In addition, we assessed cell viability during the HiBiT knock-in assay to confirm that 6×NLS-Acr is nontoxic to HUES8 and HEK293T cells (Figures S17 and S18).

### Comparing the delivery of 6×NLS-Acr and LF_N_-Acr after Cas9 nucleofection

Acr delivery was assessed in Cas9-nucleofected cells using the *GFP*-disruption and HiBiT knock-in assays, which provide functional measurements of delivery effectiveness. Specifically, we define delivery effectiveness as the extent to which the delivered Acr inhibits Cas9 activity. In contrast, delivery efficiency is defined as the ratio of intracellular Acr concentration to the total Acr concentration administered to the well. Delivery effectiveness reflects the functional impact of intracellular Acr and generally increases with delivery efficiency. However, this relationship is not strictly proportional, as Cas9 inhibition can occur when a small fraction of the administered Acr enters the cell, depending on factors such as Acr potency and the intracellular abundance of Cas9. Potency reflects how effectively the Acr inhibits Cas9 at a given concentration, in this case, the concentration required to reduce Cas9 activity by 50% (IC_50_). Efficacy is the maximum level of Cas9 inhibition achieved by the Acr and depends on delivery efficiency and effectiveness.

The *GFP*-disruption and HiBiT-knock-in assays confirmed that 6×NLS-Acr effectively enters Cas9-nucleofected cells (Figures 2 and 3). We previously used the same assay conditions to assess PA-mediated delivery of LF_N_-Acr.^31^ Hence, these assays provide a consistent framework for comparing the delivery effectiveness of different Acrs.

6×NLS-Acr (IC_50_ = 0.47 µM, Table S1) is 160-fold less potent than the LF_N_-Acr/PA system (IC_50_ = 2.9 nM, Table S1) in the *GFP*-disruption assay (Figure S19). Similarly, 6×NLS-Acr (IC_50_ = 0.63 µM, Table S1) is 200-fold less potent than the LF_N_-Acr/PA system (IC_50_ = 3.1 nM, Table S1) in the HiBiT-knock-in assay (Figure S20). This effect is expected because PA, a pore-forming protein, mediates the delivery of LF_N_-Acr. PA generally delivers cargo more efficiently than CPPs at lower concentrations.^54,55^

6×NLS-Acr inhibited Cas9 more effectively than LF_N_-Acr/PA at concentrations above 1 µM in both the GFP-disruption (efficacy: 96 versus 86%) and HiBiT knock-in assays (efficacy: 99 versus 88%) (Table S1). 6×NLS-Acr delivery does not depend on receptors; therefore, its efficiency and effectiveness increase consistently with dose, resulting in higher efficacy. In contrast, LF_N_-Acr/PA relies on receptor-mediated uptake that saturates above 1 µM, probably due to a limited number of PA-targeted receptors on the cell surface.

6×NLS-Acr is the most potent known cell-permeable protein for Cas9 inhibition, whereas LF_N_-Acr/PA is the most potent protein-based system. 6×NLS-Acr is the most efficacious Cas9 inhibitor. Depending on the application, either 6×NLS-Acr or LF_N_-Acr/PA can be more effective. Thus, these delivery strategies are complementary.

### 6×NLS-Acr Increases Cas9 Specificity

Delivery of Acrs via nucleofection enhances Cas9 specificity in cells.^22^ To determine whether 6×NLS-Acr delivery has a similar effect, we delivered Cas9 and gRNA plasmids via lipid transfection (lipofection) into HEK293T cells, then treated the cells with Acrs. After 72 h, we used next-generation sequencing (NGS) to assess Cas9 cleavage at the *EMX1* on-target and off-target sites (Figure 4A). 6×NLS-Acr (4 µM) reduced Cas9 on-target activity from 100 to 57% and off-target activity from 80 to 44, 57 to 29, 17 to 7.2, and 11 to 4.5% at off-target (OT) sites 1, 2, 3, and 4, respectively (Figure 4B). 6×NLS-Acr increased Cas9 specificity by 1.10-to 1.41-fold (*i*.*e*., 10–41%) within each target pair (Figure 4C and eq S1–S3). Across all target pairs, delivery of 6×NLS-Acr led to a 90% improvement in Cas9 specificity (total specificity increase, eq S4).

**Figure 4.**
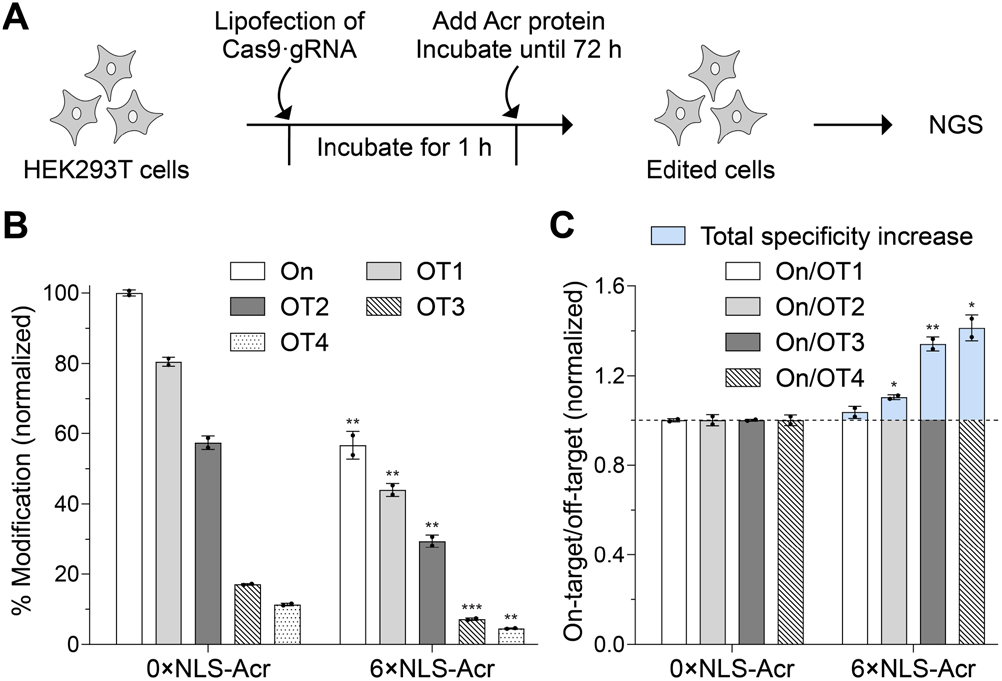
Effect of Acr delivery on genome-editing activity and specificity. (A) A cartoon illustrating the experimental setup for Acr delivery, in which next-generation sequencing (NGS) is used to analyze genome editing. (B) 6×NLS-Acr delivery regulates Cas9 activity. Plasmids encoding 2×NLS-SpCas9 (750 ng) and *EMX1*-targeting gRNA (250 ng) were delivered via lipofection into HEK293T cells. The cells were incubated at 37 °C and treated with 0×NLS-Acr (4 µM, control) or 6×NLS-Acr (4 µM) at 1 h. After 72 h, genomic DNA was extracted, sequenced with NGS, and analyzed using CRISPResso2 to determine the % modification at the on-target (On) and off-target (OT1, OT2, OT3, and OT4) sites. The values were normalized to Cas9·gRNA with 0×NLS-Acr (On) and are the mean ± SD of two independent replicates. The significance of 6×NLS-Acr delivery was determined using an unpaired, two-tailed *t*-test versus Cas9·gRNA with 0×NLS-Acr (separately for On, OT1, OT2, OT3, and OT4), where *, **, ***, and **** refer to *P* ≤ 0.05, *P* ≤ 0.01, *P* ≤ 0.001, and *P* ≤ 0.0001, respectively. (C) 6×NLS-Acr delivery increases Cas9 specificity. Specificity was calculated as the ratio of on-target to off-target % modification using data from panel B, normalized to the on-target to off-target % modification ratio of Cas9·gRNA with 0×NLS-Acr (separately for On/OT1, On/OT2, On/OT3, and On/OT4). The significance of the specificity was determined using an unpaired, two-tailed *t*-test compared to the specificity of Cas9·gRNA with 0×NLS-Acr (separately for On/OT1, On/OT2, On/OT3, and On/OT4), where *, **, ***, and **** refer to *P* ≤ 0.05, *P* ≤ 0.01, *P* ≤ 0.001, and *P* ≤ 0.0001, respectively. The total specificity increase is shaded in blue.

Treating cells with LF_N_-Acr (250 nM) and PA (20 nM) after Cas9 lipofection did not affect genome-editing activity or specificity (Figure S21). In contrast, treating cells with LF_N_-Acr (250 nM) and PA (20 nM) after Cas9 nucleofection inhibited Cas9 and increased its specificity.^31^ These contradicting results suggest that the process of lipofection hinders PA-mediated LF_N_-Acr delivery.

We also used NGS to assess the activity of a less active Cas9 variant at the *EMX1* site (Figures S22 and S23). These experiments support that 6×NLS-Acr but not LF_N_-Acr/PA enters cells to inhibit Cas9 after lipofection.

### 6×NLS-Acr Reduces the Local Concentration of DNA-Bound Cas9

We previously developed a platform to study the spatiotemporal dynamics and outcomes of Cas9-mediated genome editing.^56^ In this platform, a U2OS cell line stably expressing a Cas9–GFP fusion protein (U2OS-Cas9–GFP cells) is transfected with Ch3Rep gRNA, a truncated gRNA that targets a highly repetitive region on chromosome 3 (Ch3Rep) without inducing its cleavage (Figure 5A). Cas9–GFP in its apo form localizes in the nucleolus. But after pairing with Ch3Rep gRNA, Cas9–GFP accumulates at ∼600 target sites on Ch3Rep, forming distinct nuclear foci observable by fluorescence microscopy (Figure S24). 53BP1–mCherry, also stably expressed in this cell line, serves as a nuclear marker. We adapted this platform to directly visualize Acr-mediated inhibition of Cas9·gRNA in human cells.

**Figure 5.**
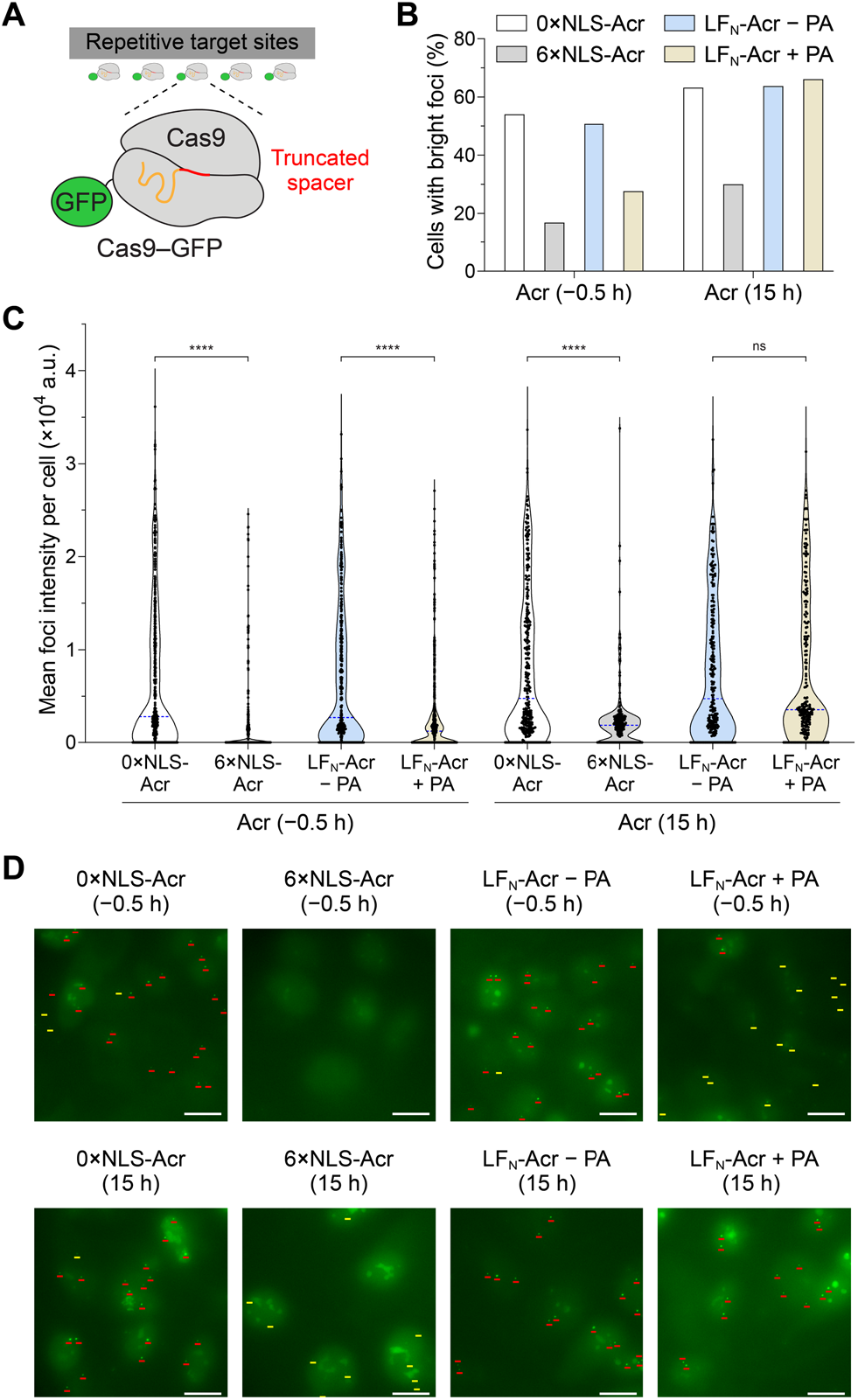
Directly visualizing the Acr-mediated inhibition of fluorescently labeled Cas9. (A) Cas9–GFP complexed with a truncated gRNA targeting a highly repetitive region on chromosome 3 (Ch3Rep) produces nuclear foci in cells. Because the Ch3Rep gRNA spacer is an 11-mer truncation, Cas9–GFP can bind the Ch3Rep target but cannot cleave it. (B, C) Quantifying the inhibition of Cas9–GFP by 6×NLS-Acr and LF_N_-Acr. Ch3Rep gRNA was delivered via lipofection into a U2OS cell line stably expressing SpCas9–GFP. Cells were treated with 6×NLS-Acr (2.5 µM) or LF_N_-Acr (2.5 µM) plus PA (20 nM) either 0.5 h before (−0.5 h) or 15 h after (15 h) gRNA transfection. Controls include cells treated with 0×NLS-Acr (2.5 µM) or LF_N_-Acr (2.5 µM) at −0.5 h or 15 h. Live cells were imaged using widefield fluorescence microscopy (60× objective) at 15 and 20 h for −0.5- and 15-h Acr treatments, respectively. Nuclear foci intensity was quantified. Cells with “bright foci” have a mean foci intensity above 2,500 a.u. and are shown as a percentage of total cells in Panel B. The mean foci intensity per cell is shown as a violin plot in Panel C, with the median of these values indicated by a dashed blue line. The total number of quantified cells per condition was 505, 297, 390, 373, 318, 263, 254, and 233 from left to right. The significance of 6×NLS-Acr and LF_N_-Acr/PA delivery was determined using a Mann–Whitney test versus 0×NLS-Acr and LF_N_-Acr delivery, respectively. ns, *, **, ***, and **** refer to not significant, *P* ≤ 0.05, *P* ≤ 0.01, *P* ≤ 0.001, and *P* ≤ 0.0001, respectively. (D) Representative maximum intensity projection images from the experiment described in Panels B and C. Visually bright foci are marked with red lines, and visually faint foci with yellow lines. This classification is qualitative and not based on quantitative measurements; therefore, the terms “visually bright” and “visually faint” are used instead. Scale bars, 20 µm.

We delivered Ch3Rep gRNA via lipofection into U2OS-Cas9–GFP cells. Cells were treated with 6×NLS-Acr (2.5 µM) or LF_N_-Acr (2.5 µM) plus PA (20 nM) either 0.5 h before (−0.5 h) or 15 h after (15 h) Ch3Rep gRNA delivery. Pretreating cells with 6×NLS-Acr or LF_N_-Acr/PA at −0.5 h prevented the binding of the Cas9–GFP·gRNA complex to Ch3Rep with decreasing effectiveness. 6×NLS-Acr reduced the mean foci intensity, which in turn decreased the percentage of cells with bright foci—those with a mean foci intensity above 2,500 a.u.— from 54 to 17% (Figures 5B and 5C). LF_N_-Acr/PA lowered the mean foci intensity, reducing the percentage of cells with bright foci from 51 to 28%. Most foci were visually faint or undetectable in cells pretreated with LFN-Acr/PA or 6×NLS-Acr, respectively (Figures 5D and S24). Treating cells with 6×NLS-Acr at 15 h reduced the binding of the Cas9– GFP·gRNA complex at Ch3Rep, lowering the mean foci intensity and percentage of cells with bright foci from 63 to 30% (Figures 5B and 5C). Foci were visually faint after treating cells with 6×NLS-Acr at 15 h (Figures 5D and S24). Treatment with LF_N_-Acr/PA at 15 h failed to inhibit the Cas9–GFP·gRNA complex, confirming that lipofection reagents interfere with PA-mediated Acr delivery.

### Comparing the delivery of 6×NLS-Acr and LF_N_-Acr after Cas9 and gRNA lipofection

6×NLS-Acr effectively entered cells after Cas9 and gRNA lipofection (Figures 4, 5, S22, and S24). In contrast, PA-mediated LF_N_-Acr delivery was undetectable after Cas9 and gRNA lipofection (Figures 5, S21, S23, and S24). A possible explanation for these results is that lipofection reagents can shield the cellular receptors required for PA pore formation and subsequent LF_N_-Acr uptake. Therefore, the one-component nature and receptor-independent delivery mechanism of 6×NLS-Acr are advantageous when Acr delivery follows Cas9 or gRNA lipofection.

### 6×NLS-Acr Enters Cells via a Non-Endocytic Direct Delivery Mechanism

To directly visualize Acr delivery and assess its efficiency, we conjugated 0×NLS-Acr and 6×NLS-Acr to a Cy5 dye using thiol–maleimide chemistry (Figures 6A and S25). We then incubated U2OS cells and INS-1E rat β-cells with Cy5-0×NLS-Acr (1–4 µM) or Cy5-6×NLS-Acr (1–4 µM) and visualized Acr uptake using confocal fluorescence microscopy. Cy5-6×NLS-Acr fluorescence was predominantly diffuse and localized to the nucleus in both cell types (Figures 6B and S26–S28). Discrete cytosolic puncta were observed in a small subset of cells, suggesting that while endocytic uptake occurs, it is largely outcompeted by direct cell entry. Cy5-6×NLS-Acr fluorescence increased in a dose- and time-dependent manner (Figures S27 and S28). Cellular uptake occurred within the first 5 min. Similar to arginine-rich CPP– protein conjugates that directly transduce cells,^57^ Cy5-6×NLS-Acr is often localized to RNA-rich nucleoli, visible as high-intensity, round structures within the nucleus. In contrast, Cy5-0×NLS-Acr uptake was undetectable relative to that of Cy5-6×NLS-Acr, confirming that the 6×NLS tag mediates Acr delivery. Cy5-6×NLS-Acr exhibited exclusively diffuse fluorescence when incubated with cells at 4 °C, a condition that inhibits endocytosis (Figures 6B and S29). Furthermore, pretreating cells with the macropinocytosis inhibitor amiloride and the endosomal acidification inhibitors bafilomycin A1 and chloroquine did not affect Cy5-6×NLS-Acr uptake, compared to that in a water-treated control (Figure S30). These findings indicate that Cy5-6×NLS-Acr can enter cells via a direct delivery mechanism and that inhibiting endocytic pathways does not reduced the efficiency of direct uptake.

**Figure 6.**
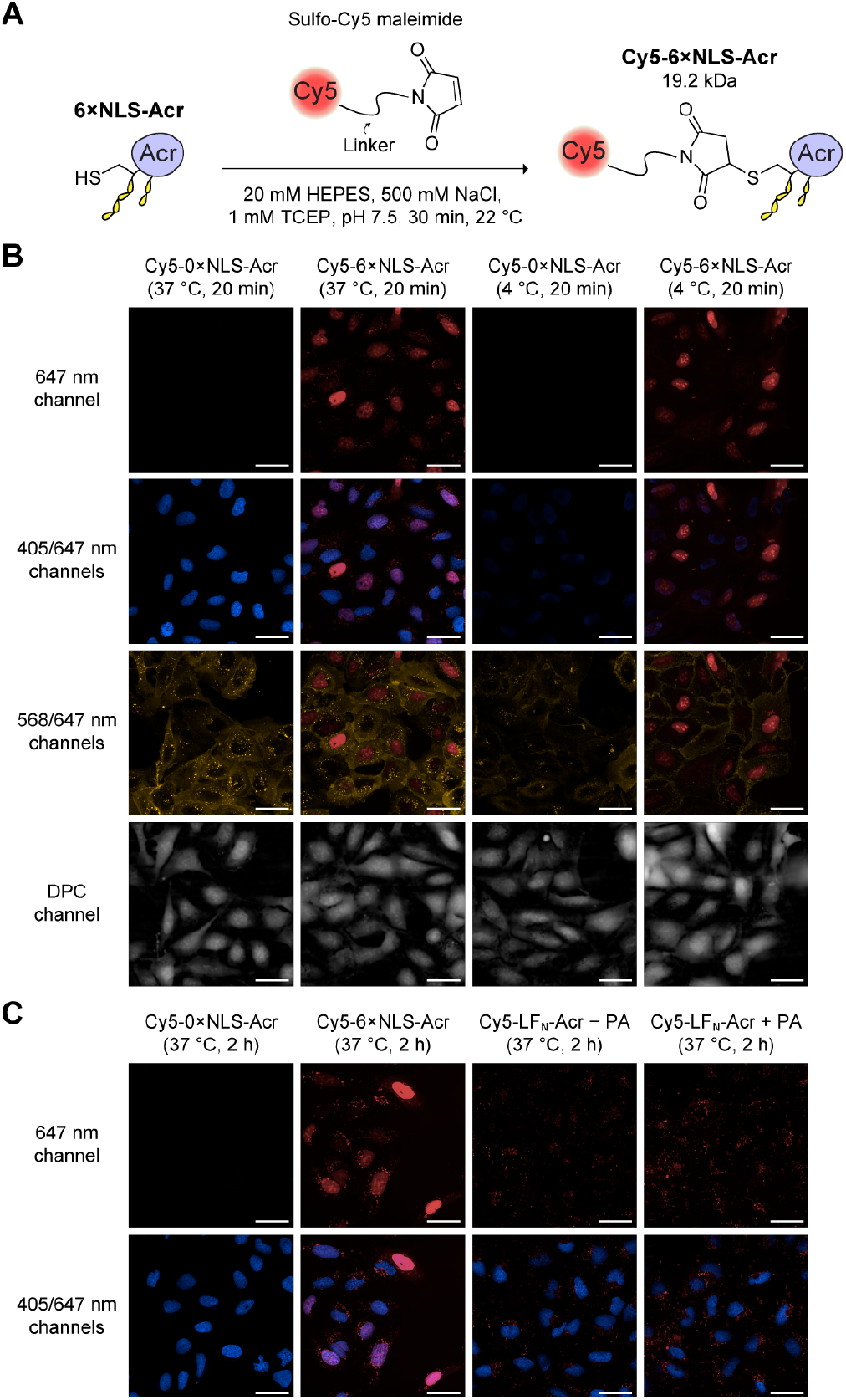
Visualizing the delivery of fluorescently labeled Acrs. Synthesis of Cy5-6×NLS-Acr. 6×NLS-Acr was reacted with sulfo-Cyanine5 (sulfo-Cy5) maleimide to generate Cy5-6×NLS-Acr. (B, C) U2OS cells were seeded and incubated overnight at 37 °C. Cells were then treated with Cy5-0×NLS-Acr (1.5 µM), Cy5-6×NLS-Acr (1.5 µM), and Cy5-LF_N_-Acr (2.5 µM) ± PA (20 nM), and incubated either for 20 min at 37 or 4 °C (panel B) or for 2 h at 37 °C (panel C). Cells were washed and stained with Hoechst 33342 and WGA555 in FluoroBrite DMEM, then imaged using confocal microscopy (60× objective) to detect fluorescence from Hoechst 33342 (blue nuclear marker; 405 nm excitation), WGA555 (yellow cytosolic marker; 568 nm excitation), and Cy5 (red; 647 nm excitation). Digital phase contrast (DPC) microscopy was also performed. Scale bars, 40 µm.

For comparison, we synthesized Cy5-LF_N_-Acr and visualized its PA-mediated delivery using confocal fluorescence microscopy (Figures 6C, S31, and S32). Cy5-LF_N_-Acr fluorescence appeared as discrete cytosolic puncta, consistent with endosomal entrapment. Fluorescence in cells treated with Cy5-LF_N_-Acr (2.5 µM) was nearly indistinguishable from that in cells co-treated with Cy5-LF_N_-Acr (2.5 µM) and PA (20 nM), suggesting that the amount of Cy5-LF_N_-Acr translocated to the cytosol through the PA pore is below the detection limit or that the Cy5 dye prevents LF_N_-Acr from translocating through the PA pore. To distinguish between these two possibilities, we co-treated U2OS-EGFP.PEST cells with Cy5-LF_N_-Acr and PA following Cas9 nucleofection. We then assessed *GFP* disruption (Figures S33 and S34) and directly visualized LF_N_-Acr delivery using the Cy5 tag (Figure S35). Cy5-LF_N_-Acr inhibited Cas9 activity in the presence of PA but not when administered alone. Consistently, Cy5-LF_N_-Acr fluorescence in cells remained indistinguishable regardless of PA addition. Thus, Cy5-LF_N_-Acr translocates through the PA pore, but its cellular concentration is below the detection limit of confocal microscopy or overshadowed by endosomally entrapped Cy5-LF_N_-Acr. Together, these findings indicate that 6×NLS-Acr delivery is more efficient than PA-mediated delivery of LF_N_-Acr above 1 µM, due to its ability to achieve higher intracellular Acr concentrations.

## DISCUSSION

We introduce 6×NLS-Acr—the first cell-permeable anti-CRISPR protein—and its application to regulating CRISPR-Cas9 activity and specificity. 6×NLS-Acr enters cells within 5 min, reaching higher intracellular concentrations and inhibiting Cas9 with higher efficacy than LF_N_-Acr/PA, the most potent protein-based anti-CRISPR delivery system. Therefore, 6×NLS-Acr is the most efficacious cell-permeable Cas9 inhibitor.

6×NLS-Acr inhibited Cas9 in spheroid-like 3D cell aggregates. We assessed 6×NLS-Acr delivery in a loosely aggregated spheroid model because spheroids have diffusional limits similar to in vivo tissues.^58^ The observed decrease in GFP intensity with spheroid depth (Figures 2D, S14, and S15) indicates limited penetration of 6×NLS-Acr into the inner region. This result was also observed with LF_N_-Acr/PA because diffusion restricts the penetration of molecules to the outer layers of spheroids (50–200 µm in depth).^31,58^ 6×NLS-Acr inhibited Cas9 within the outer 200 µm region of the spheroid with effectiveness comparable to that of LF_N_-Acr/PA.^31^

6×NLS-Acr inhibited Cas9 in a dose- and time-dependent manner in embryonic stem cells (Figures 3B and 3C). 6×NLS-Acr inhibited Cas9 more effectively than LF_N_-Acr at concentrations above 1 µM, highlighting the potential of 6×NLS-Acr for ex vivo, Acr-based stem cell therapy applications.^25,33,59^

6×NLS-Acr delivery increased Cas9 specificity by up to 41% per off-target site and by 90% when considering multiple off-target sites (Figure 4C). Due to the limitations of target-amplicon sequencing with NGS, we assessed genome editing at only four known *EMX1* off-target sites. Thus, the 90% figure underrepresents the total increase in Cas9 specificity achieved by delivering 6×NLS-Acr. We anticipate that genome-wide off-target analyses (*e*.*g*., GUIDE-Seq^60^) will reveal the full potential of 6×NLS-Acr in the future. Furthermore, milder 6×NLS-Acr variants could be engineered to selectively inhibit Cas9 off-target activity, as previously demonstrated with anti-CRISPR vector systems.^29,61,62^

PA-mediated delivery of LF_N_-Acr after Cas9 nucleofection increased genome-editing specificity by 40%.^31^ Yet, LF_N_-Acr delivery after Cas9 lipofection did not affect genome-editing activity or specificity (Figures S21 and S23), likely because lipofection reagents disrupt PA’s receptor-dependent protein delivery mechanism. PA effectively mediated the delivery of LF_N_-Acr before Cas9 lipofection (Figures 5 and S24), supporting this hypothesis. Further research is needed to understand how lipofection reagents affect PA-mediated protein delivery and whether other lipid-based Cas9 delivery methods (*e*.*g*., lipid nanoparticles) interfere with PA’s mechanism.

As a one-component system that bypasses receptor-mediated uptake, 6×NLS-Acr is more practical than LF_N_-Acr/PA when receptors are inaccessible, for example, when Cas9-lipofection reagents coat the cell surface. Unlike receptor-dependent systems, 6×NLS-Acr delivery is less prone to saturation. Furthermore, like other CPP-based protein delivery methods,^63,64^ 6×NLS-Acr delivery might be effective in bacterial and plant cells that lack the anthrax receptors targeted by LF_N_-Acr/PA. Besides delivery applications, the cationic 6×NLS tag could serve as an affinity tag and enable the adsorption of Acrs to a solid support.^65^

6×NLS-Acr and LF_N_-Acr/PA enabled live visualization of the inhibition of a fluorescent Cas9·gRNA complex that binds but does not cleave DNA in cells (Figures 5 and S24). 6×NLS-Acr inhibited Cas9·gRNA more effectively than LF_N_-Acr/PA.

6×NLS-Acr inhibited Cas9·gRNA less effectively when delivered at 15 h than at −0.5 h. After 15 h, Cas9·gRNA has bound its DNA target. Because AcrIIA4 cannot bind to dCas9·gRNA·DNA,^20^ 6×NLS-Acr likely only inhibits Cas9– GFP·gRNA transiently displaced from DNA (*e*.*g*., by cellular factors). LF_N_-Acr/PA did not inhibit Cas9–GFP·gRNA when delivered at 15 h, likely because lipofection reagents blocked its cellular entry. In contrast, our previous CRISPRa experiments showed that PA-mediated delivery of LF_N_-Acr at 12 h can inhibit DNA binding by nucleofected dCas9.^31^

Plasmids expressing Acrs were previously used to visualize the inhibition of a fluorescent dCas9 system in cells.^17,66,67^ Similarly, 6×NLS-Acr produced a comparable inhibitory effect on the DNA-binding activity of Cas9–GFP·gRNA, although plasmid-based Acr expression was not necessary. In both experiments, inhibition of the fluorescent Cas9 system resulted in diminished foci intensity and elevated background fluorescence.

Our work also established a site-specific Acr conjugation platform. Conjugation of 6×NLS-Acr to a fluorescent dye facilitated elucidation of its delivery mechanism (Figures 6 and S25–S30). 6×NLS-Acr entered cells within 5 min (Figure S28), primarily via direct cytosolic uptake, with minimal endocytic entrapment. Inhibition of endocytic pathways by lowering the incubation temperature to 4 °C or using small molecules did not reduce the efficiency of direct cellular uptake of 6×NLS-Acr (Figures 6B, S29, and S30). 6×NLS-Acr achieved intracellular concentrations much higher than LF_N_-Acr, highlighting the advantage of a receptor-independent delivery mechanism (Figure 6C). In the future, our site-specific conjugation platform could enable the addition of cleavable moieties to regulate Acr delivery and activity reversibly.^68,69^

Narrowing the cell entry time of 6×NLS-Acr to ≤5 min has important implications for future studies of Cas9 kinetics and DNA repair. For example, our time-dependent studies using the HiBiT knock-in assay confirmed that Cas9 RNP remains active for at least 16 h ± 5 min in human embryonic stem cells. By tracking Cas9 and DNA repair dynamics and understanding the timescale of 6×NLS-Acr delivery, we are poised to explore a broader range of fundamental questions in genome biology.

In vivo Acr delivery has been validated in mice without observable toxicity.^27,35^ These studies, however, did not determine the safety and immunity profiles of delivered Acrs. 6×NLS-Acr delivery could be effective in vivo, though immunogenic responses against the 6×NLS tag or Acr would limit its use in clinical settings. When delivered to mice via intracranial injection, 4×NLS-Cas9-2×NLS RNP caused an immune response, which was lowered by removing endotoxin contamination.^50^ In addition, the efficacy of 6×NLS-Acr delivery in vivo can be improved by employing immunosuppressants and screening for pre-existing immunity.^14^

Without in vivo studies, the tissue specificity of 6×NLS-Acr is unclear. Still, 6×NLS-Acr could be targetable to neurons via intracranial injection, as demonstrated previously with cell-permeable NLS-Cas9 variants.^47,50,51^ Additionally, we can conjugate cell-targeting ligands to 6×NLS-Acr to improve its tissue specificity.^70^ Ongoing work focuses on determining the efficiency, immunogenicity, and tissue specificity of 6×NLS-Acr in vivo to validate its use as a co-therapy for therapeutic genome editing.

## CONCLUSION

We established the first peptide-based delivery method for introducing Acrs into human cells. 6×NLS-Acr is the most potent cell-permeable anti-CRISPR protein identified to date and enters cells via direct protein transduction. It effectively inhibits Cas9 in stem cell and 3D cultures, enhancing genome-editing specificity by nearly 100%. 6×NLS-Acr holds promise for studying the dynamics of genome editing and reducing off-target genome editing, both ex vivo and in vivo, potentially improving the safety and efficacy of therapeutic genome editing.

## ASSOCIATED CONTENT

### Supporting Information

The Supporting Information is available free of charge on the ACS Publications website at DOI: 10.1021/xxxx.xxxxxxx. Experimental methods, Figures S1–S35, Tables S1–S11, and plasmid sequences (PDF)

## AUTHOR INFORMATION

### Authors

**Axel O. Vera** – *Department of Chemistry, Massachusetts Institute of Technology, Cambridge, Massachusetts 02139, United States; Chemical Biology and Therapeutics Science Program, Broad Institute of MIT and Harvard, Cambridge, Massachusetts 02142, United States;*

**Franklin J. Avilés-Vázquez** – *Howard Hughes Medical Institute and Program in Cellular and Molecular Medicine, Boston Children’s Hospital, Boston, Massachusetts 02115, United States; Department of Biophysics and Biophysical Chemistry, Johns Hopkins University, Baltimore, Maryland 21218, United States;*

**Taekjip Ha** – *Howard Hughes Medical Institute and Program in Cellular and Molecular Medicine, Boston Children’s Hospital, Boston, Massachusetts 02115, United States; Department of Pediatrics, Harvard Medical School, Boston, Massachusetts 02115, United States;*

## Author Contributions

A.O.V., A.C., and R.T.R. developed the project idea. A.O.V. and F.J.A.V. designed and performed experiments. A.O.V. wrote the original manuscript draft. A.O.V., F.J.A.V., T.H., A.C., and R.T.R. analyzed data. A.C. and R.T.R. supervised the research. All authors read and approved the final manuscript.

## Funding

A.O.V. was supported by an MIT Dean of Science Fellowship, an NIH predoctoral fellowship (F31 GM148042), and an HHMI Gilliam fellowship (GT15945). Cell imaging was performed with an Opera Phenix High-Throughput imaging system that was supported by Grant S10 OD026839 (NIH). This work was supported by Grants R01 GM137606, R35 GM148220, R35 GM122569, and U01 DK127432 (NIH), and by the Merkin Institute of Transformative Technologies in Healthcare.

## Notes

The authors declare the following competing financial interest: The Broad Institute of MIT and Harvard has filed patent applications claiming the invention of genome editing methods described in this manuscript. Inventors: A.O.V., A.C., and R.T.R.

## ACKNOWLEDGMENTS

The authors thank Dr. Santosh Chaudhary (Broad Institute of MIT and Harvard) for providing a plasmid for the bacterial production and purification of wild-type AcrIIA4 and sharing his purification method, Michael C. Stark (Broad Institute of MIT and Harvard) for acquiring HUES 8 cells for our laboratory and providing INS-1E rat β-cells for Figure S27, and Prof. J. Keith Joung (Harvard Medical School) for providing the U2OS-EGFP.PEST cell line.

## For Table of Contents use only

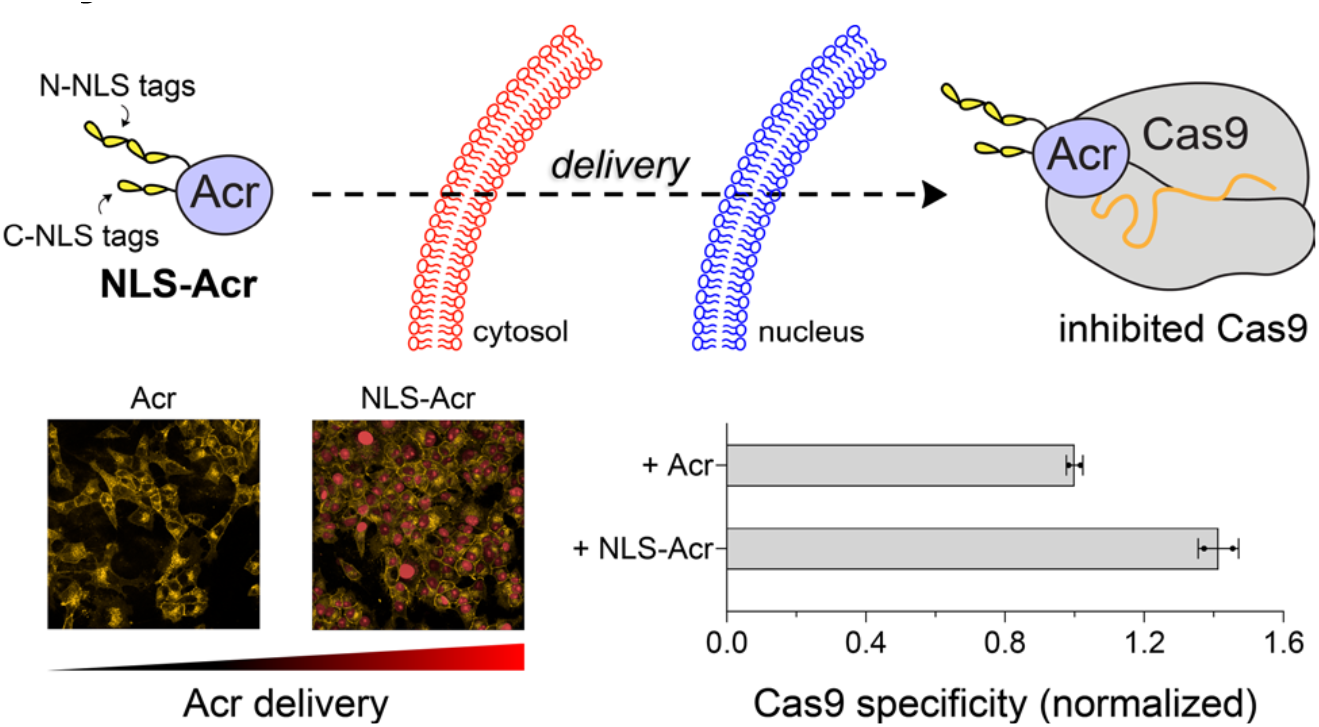

## REFERENCES

(1) Barrangou, R.; Fremaux, C.; Deveau, H.; Richards, M.; Boyaval, P.; Moineau, S.; Romero, D. A.; Horvath, P. CRISPR provides acquired resistance against viruses in prokaryotes. Science 2007, 315, 1709–1712.

(2) Jinek, M.; Chylinski, K.; Fonfara, I.; Hauer, M.; Doudna, J. A.; Charpentier, E. A programmable dual-RNA–guided DNA endonuclease in adaptive bacterial immunity. Science 2012, 337, 816–821.

(3) Cong, L.; Ran, F. A.; Cox, D.; Lin, S.; Barretto, R.; Habib, N.; Hsu, P. D.; Wu, X.; Jiang, W.; Marraffini, L. A. Multiplex genome engineering using CRISPR/Cas systems. Science 2013, 339, 819–823.

(4) Mali, P.; Yang, L.; Esvelt, K. M.; Aach, J.; Guell, M.; DiCarlo, J. E.; Norville, J. E.; Church, G. M. RNA-guided human genome engineering via Cas9. Science 2013, 339, 823–826.

(5) Frangoul, H.; Altshuler, D.; Cappellini, M. D.; Chen, Y.-S.; Domm, J.; Eustace, B. K.; Foell, J.; de la Fuente, J.; Grupp, S.; Handgretinger, R.; Ho, T. W.; Kattamis, A.; Kernytsky, A.; Lekstrom-Himes, J.; Li, A. M.; Locatelli, F.; Mapara, M. Y.; de Montalembert, M.; Rondelli, D.; Sharma, A.; Sheth, S.; Soni, S.; Steinberg, M. H.; Wall, D.; Yen, A.; Corbacioglu, S. CRISPR-Cas9 gene editing for sickle cell disease and β-thalassemia. N. Engl. J. Med. 2020, 384, 252–260.

(6) Gillmore, J. D.; Gane, E.; Taubel, J.; Kao, J.; Fontana, M.; Maitland, M. L.; Seitzer, J.; O’Connell, D.; Walsh, K. R.; Wood, K.; Phillips, J.; Xu, Y.; Amaral, A.; Boyd, A. P.; Cehelsky, J. E.; McKee, M. D.; Schiermeier, A.; Harari, O.; Murphy, A.; Kyratsous, C. A.; Zambrowicz, B.; Soltys, R.; Gutstein, D. E.; Leonard, J.; Sepp-Lorenzino, L.; Lebwohl, D. CRISPR-Cas9 in vivo gene editing for transthyretin amyloidosis. N. Engl. J. Med. 2021, 385, 493–502.

(7) Chiesa, R.; Georgiadis, C.; Syed, F.; Zhan, H.; Etuk, A.; Gkazi, S. A.; Preece, R.; Ottaviano, G.; Braybrook, T.; Chu, J.; Kubat, A.; Adams, S.; Thomas, R.; Gilmour, K.; O’Connor, D.; Vora, A.; Qasim, W. Base-edited CAR7 T cells for relapsed T-cell acute lymphoblastic leukemia. N. Engl. J. Med. 2023, 389, 899–10.

(8) Fu, Y.; Foden, J. A.; Khayter, C.; Maeder, M. L.; Reyon, D.; Joung, J. K.; Sander, J. D. High-frequency off-target mutagenesis induced by CRISPR-Cas nucleases in human cells. Nat. Biotechnol.2013, 31, 822–826.

(9) Zuo, E.; Sun, Y.; Wei, W.; Yuan, T.; Ying, W.; Sun, H.; Yuan, L.; Steinmetz, L. M.; Li, Y.; Yang, H. Cytosine base editor generates substantial off-target single-nucleotide variants in mouse embryos. Science 2019, 364, 289–292.

(10) Liang, S.-Q.; Liu, P.; Ponnienselvan, K.; Suresh, S.; Chen, Z.; Kramme, C.; Chatterjee, P.; Zhu, L. J.; Sontheimer, E. J.; Xue, W.; Wolfe, S. A. Genome-wide profiling of prime editor off-target sites in vitro and in vivo using PE-tag. Nat. Methods 2023, 20, 898–907.

(11) Guo, C.; Ma, X.; Gao, F.; Guo, Y. Off-target effects in CRISPR/Cas9 gene editing. Front. Bioeng. Biotechnol. 2023, 11, 1143157.

(12) Mehravar, M.; Shirazi, A.; Nazari, M.; Banan, M. Mosaicism in CRISPR/Cas9-mediated genome editing. Dev. Biol. 2019, 445, 156–162.

(13) Zuccaro, M. V.; Xu, J.; Mitchell, C.; Marin, D.; Zimmerman, R.; Rana, B.; Weinstein, E.; King, R. T.; Palmerola, K. L.; Smith, M. E.; Tsang, S. H.; Goland, R.; Jasin, M.; Lobo, R.; Treff, N.; Egli, D. Allele-specific chromosome removal after Cas9 cleavage in human embryos. Cell 2020, 183, 1650–1664.

(14) Ewaisha, R.; Anderson, K. S. Immunogenicity of CRISPR therapeutics—critical considerations for clinical translation. Front. Bioeng. Biotechnol. 2023, 11, 1138596.

(15) Haapaniemi, E.; Botla, S.; Persson, J.; Schmierer, B.; Taipale, J. CRISPR–Cas9 genome editing induces a p53-mediated DNA damage response. Nat. Med. 2018, 24, 927–930.

(16) Fiumara, M.; Ferrari, S.; Omer-Javed, A.; Beretta, S.; Albano, L.; Canarutto, D.; Varesi, A.; Gaddoni, C.; Brombin, C.; Cugnata, F.; Zonari, E.; Naldini, M. M.; Barcella, M.; Gentner, B.; Merelli, I.; Naldini, L. Genotoxic effects of base and prime editing in human hematopoietic stem cells. Nat. Biotechnol. 2023, 877–891.

(17) Pawluk, A.; Amrani, N.; Zhang, Y.; Garcia, B.; Hidalgo-Reyes, Y.; Lee, J.; Edraki, A.; Shah, M.; Sontheimer, E. J.; Maxwell, K. L.; Davidson, R. Naturally occurring off-switches for CRISPR-Cas9. Cell 2016, 167, 1829–1838.

(18) Rauch, B. J.; Silvis, M. R.; Hultquist, J. F.; Waters, C. S.; McGregor, M. J.; Krogan, N. J.; Bondy-Denomy, J. Inhibition of CRISPR-Cas9 with bacteriophage proteins. Cell 2017, 168, 150–158.

(19) Hwang, S.; Maxwell, K. L. Diverse mechanisms of CRISPR-Cas9 inhibition by type II anti-CRISPR proteins. J. Mol. Biol. 2023, 435, 168041.

(20) Yang, H.; Patel, D. J. Inhibition mechanism of an anti-CRISPR suppressor AcrIIA4 targeting SpyCas9. Mol. Cell 2017, 67, 117–127.

(21) Dong Guo, M.; Wang, S.; Zhu, Y.; Wang, S.; Xiong, Z.; Yang, J.; Xu, Z.; Huang, Z. Structural basis of CRISPR-SpyCas9 inhibition by an anti-CRISPR protein. Nature 2017, 546, 436–439.

(22) Shin, J.; Jiang, F.; Liu, J.-J.; Bray, N. L.; Rauch, B. J.; Baik, S. H.; Nogales, E.; Bondy-Denomy, J.; Corn, J. E.; Doudna, J. A. Disabling Cas9 by an anti-CRISPR DNA mimic. Sci. Adv. 2017, 3, e1701620.

(23) Kraus, C.; Sontheimer, E. J. Applications of anti-CRISPR proteins in genome editing and biotechnology. J. Mol. Biol. 2023, 435, 168120.

(24) Gebhardt, C. M.; Niopek, D., Anti-CRISPR proteins and their application to control CRISPR effectors in mammalian systems. In Mammalian Synthetic Systems, Ceroni, F.; Polizzi, K., Eds. Springer US: New York, NY, 2024; pp 205–231.

(25) Li, C.; Psatha, N.; Gil, S.; Wang, H.; Papayannopoulou, T.; Lieber, HDAd5/35++ adenovirus vector expressing anti-CRISPR peptides decreases CRISPR/Cas9 toxicity in human hematopoietic stem cells. Mol. Ther. Methods Clin. Dev. 2018, 9, 390–401.

(26) Matsumoto, D.; Tamamura, H.; Nomura, W. A cell cycle-dependent CRISPR-Cas9 activation system based on an anti-CRISPR protein shows improved genome editing accuracy. Commun. Biol. 2020, 3, 601.

(27) Lee, J.; Mou, H.; Ibraheim, R.; Liang, S.-Q.; Liu, P.; Xue, W.; Sontheimer, E. J. Tissue-restricted genome editing in vivo specified by microRNA-repressible anti-CRISPR proteins. RNA 2019, 25, 1421–1431.

(28) Hoffmann, M. D.; Aschenbrenner, S.; Grosse, S.; Rapti, K.; Domenger, C.; Fakhiri, J.; Mastel, M.; Börner, K.; Eils, R.; Grimm, D.; Niopek, D. Cell-specific CRISPR–Cas9 activation by microRNA-dependent expression of anti-CRISPR proteins. Nucleic Acids Res. 2019, 47, e75.

(29) Aschenbrenner, S.; Kallenberger, S. M.; Hoffmann, M. D.; Huck, A.; Eils, R.; Niopek, D. Coupling Cas9 to artificial inhibitory domains enhances CRISPR-Cas9 target specificity. Sci. Adv. 2020, 6, eaay0187.

(30) Jain, S.; Xun, G.; Abesteh, S.; Ho, S.; Lingamaneni, M.; Martin, T. A.; Tasan, I.; Yang, C.; Zhao, H. Precise regulation of Cas9-mediated genome engineering by anti-CRISPR-based inducible CRISPR controllers. ACS Synth. Biol. 2021, 10, 1320–1327.

(31) Vera, A. O.; Truex, N. L.; Sreekanth, V.; Pentelute, B. L.; Choudhary, A.; Raines, R. T. Protective antigen–mediated delivery of an anti-CRISPR protein for precision genome editing. Proc. Natl. Acad. Sci. U. S. A. 2025, 122, e2426960122.

(32) Bubeck, F.; Hoffmann, M. D.; Harteveld, Z.; Aschenbrenner, S.; Bietz, A.; Waldhauer, M. C.; Börner, K.; Fakhiri, J.; Schmelas, C.; Dietz, L.; Grimm, D.; Correia, B. E.; Eils, R.; Niopek, D. Engineered anti-CRISPR proteins for optogenetic control of CRISPR–Cas9. Nat. Methods 2018, 15, 924–927.

(33) Liu, X. S.; Wu, H.; Krzisch, M.; Wu, X.; Graef, J.; Muffat, J.; Hnisz, D.; Li, C. H.; Yuan, B.; Xu, C.; Li, Y.; Vershkov, D.; Cacace, A.; Young, R. A.; Jaenisch, R. Rescue of Fragile X syndrome neurons by DNA methylation editing of the FMR1 gene. Cell 2018, 172, 979–992.

(34) Nakamura, M.; Srinivasan, P.; Chavez, M.; Carter, M. A.; Dominguez, A. A.; La Russa, M.; Lau, M. B.; Abbott, T. R.; Xu, X.; Zhao, D.; Gao, Y.; Kipniss, N. H.; Smolke, C. D.; Bondy-Denomy, J.; Qi, L. S. Anti-CRISPR-mediated control of gene editing and synthetic circuits in eukaryotic cells. Nat. Commun. 2019, 10, 194.

(35) Beyersdorf, J. P.; Bawage, S.; Iglesias, N.; Peck, H. E.; Hobbs, R. A.; Wroe, J. A.; Zurla, C.; Gersbach, C. A.; Santangelo, P. J. Robust, durable gene activation in vivo via mRNA-encoded activators. ACS Nano 2022, 16, 5660–5671.

(36) Wang, D.; Zhang, F.; Gao, G. CRISPR-based therapeutic genome editing: Strategies and in vivo delivery by AAV vectors. Cell 2020, 181, 136–150.

(37) Barkau, C. L.; O’Reilly, D.; Eddington, S. B.; Damha, M. J.; Gagnon, K. T. Small nucleic acids and the path to the clinic for anti-CRISPR. Biochem. Pharmacol. 2021, 189, 114492.

(38) Gangopadhyay, S. A.; Cox, K. J.; Manna, D.; Lim, D.; Maji, B.; Zhou, Q.; Choudhary, A. Precision control of CRISPR-Cas9 using small molecules and light. Biochemistry 2019, 58, 234–244.

(39) Klimpel, K. R.; Molloy, S. S.; Thomas, G.; Leppla, S. H. Anthrax toxin protective antigen is activated by a cell surface protease with the sequence specificity and catalytic properties of furin. Proc. Natl. Acad. Sci. U. S. A. 1992, 89, 10277–10281.

(40) Bradley, K. A.; Mogridge, J.; Mourez, M.; Collier, R. J.; Young, J. T. Identification of the cellular receptor for anthrax toxin. Nature 2001, 414, 225–229.

(41) Scobie, H. M.; Rainey, G. J. A.; Bradley, K. A.; Young, J. A. T. Human capillary morphogenesis protein 2 functions as an anthrax toxin receptor. Proc. Natl. Acad. Sci. U. S. A. 2003, 100, 5170–5174.

(42) Lange, A.; Mills, R. E.; Lange, C. J.; Stewart, M.; Devine, S. E.; Corbett, A. H. Classical nuclear localization signals: Definition, function, and interaction with importin α. J. Biol. Chem. 2007, 282, 5101–5105.

(43) Kalderon, D.; Roberts, B. L.; Richardson, W. D.; Smith, A. E. A short amino acid sequence able to specify nuclear location. Cell 1984, 39, 499–509.

(44) Morris, M. C.; Depollier, J.; Mery, J.; Heitz, F.; Divita, G. A peptide carrier for the delivery of biologically active proteins into mammalian cells. Nat. Biotechnol. 2001, 19, 1173–1176.

(45) Böhmová, E.; Machová, D.; Pechar, M.; Pola, R.; Venclíková, K.; Janoušková, O.; Etrych, T. Cell-penetrating peptides: A useful tool for the delivery of various cargoes into cells. Physiol. Res. 2018, 67 (Suppl. 2), S267–S279.

(46) Liu, J.; Gaj, T.; Wallen, M. C.; Barbas, C. F. Improved cell-penetrating zinc-finger nuclease proteins for precision genome engineering. Mol. Ther. Nucleic Acids 2015, 4, e232.

(47) Staahl, B. T.; Benekareddy, M.; Coulon-Bainier, C.; Banfal, A. A.; Floor, S. N.; Sabo, J. K.; Urnes, C.; Munares, G. A.; Ghosh, A.; Doudna, J. Efficient genome editing in the mouse brain by local delivery of engineered Cas9 ribonucleoprotein complexes. Nat. Biotechnol. 2017, 35, 431–434.

(48) Lobba, M. J.; Fellmann, C.; Marmelstein, A. M.; Maza, J. C.; Kissman, E. N.; Robinson, S. A.; Staahl, B. T.; Urnes, C.; Lew, R. J.; Mogilevsky, C. S.; Doudna, J. A.; Francis, M. B. Site-specific bioconjugation through enzyme-catalyzed tyrosine-cysteine bond formation. ACS Cent. Sci. 2020, 6, 1564–1571.

(49) Zhang, Z.; Baxter, A. E.; Ren, D.; Qin, K.; Chen, Z.; Collins, S. M.; Huang, H.; Komar, C. A.; Bailer, P. F.; Parker, J. B.; Blobel, G. A.; Kohli, R. M.; Wherry, E. J.; Berger, S. L.; Shi, J. Efficient engineering of human and mouse primary cells using peptide-assisted genome editing. Nat. Biotechnol. 2023, 42, 305–315.

(50) Stahl, E. C.; Sabo, J. K.; Kang, M. H.; Allen, R.; Applegate, E.; Kim, S. E.; Kwon, Y.; Seth, A.; Lemus, N.; Salinas-Rios, V.; Soczek, K. M.; Trinidad, M.; Vo, L. T.; Jeans, C.; Wozniak, A.; Morris, T.; Kimberlin, A.; Foti, T.; Savage, D. F.; Doudna, J. A. Genome editing in the mouse brain with minimally immunogenic Cas9 RNPs. Mol. Ther. 2023, 31, 2422–2438.

(51) Chen, K.; Stahl, E. C.; Kang, M. H.; Xu, B.; Allen, R.; Trinidad, M.; Doudna, J. A. Engineering self-deliverable ribonucleoproteins for genome editing in the brain. Nat. Commun. 2024, 15, 1727.

(52) Johnston, R. K.; Seamon, K. J.; Saada, E. A.; Podlevsky, J. D.; Branda, S. S.; Timlin, J. A.; Harper, J. C. Use of anti-CRISPR protein AcrIIA4 as a capture ligand for CRISPR/Cas9 detection. Biosens. Bioelectron. 2019, 141, 111361.

(53) Reyon, D.; Tsai, S. Q.; Khayter, C.; Foden, J. A.; Sander, J. D.; Joung, J. K. FLASH assembly of TALENs for high-throughput genome editing. Nat. Biotechnol. 2012, 30, 460–465.

(54) Liao, X.; Rabideau, A. E.; Pentelute, B. L. Delivery of antibody mimics into mammalian cells via anthrax toxin protective antigen. ChemBioChem 2014, 15, 2458–2466.

(55) Verdurmen, W. P. R.; Mazlami, M.; Pluckthun, A. A quantitative comparison of cytosolic delivery via different protein uptake systems. Sci. Rep. 2017, 7, 13194.

(56) Liu, Y.; Zou, R. S.; He, S.; Nihongaki, Y.; Li, X.; Razavi, S.; Wu, B.; Ha, T. Very fast CRISPR on demand. Science 2020, 368, 1265–1269.

(57) Herce, H. D.; Schumacher, D.; Schneider, A. F. L.; Ludwig, A. K.; Mann, F. A.; Fillies, M.; Kasper, M.-A.; Reinke, S.; Krause, E.; Leonhardt, H.; Cardoso, M. C.; Hackenberger, C. P. R. Cell-permeable nanobodies for targeted immunolabelling and antigen manipulation in living cells. Nat. Chem. 2017, 9, 762–771.

(58) Mehta, G.; Hsiao, A. Y.; Ingram, M.; Luker, G. D.; Takayama, S. Opportunities and challenges for use of tumor spheroids as models to test drug delivery and efficacy. J. Control. Release 2012, 164, 192–204.

(59) Hendriks, D.; Clevers, H.; Artegiani, B. CRISPR-Cas tools and their application in genetic engineering of human stem cells and organoids. Cell Stem Cell 2020, 27, 705–731.

(60) Tsai, S. Q.; Zheng, Z.; Nguyen, N. T.; Liebers, M.; Topkar, V. V.; Thapar, V.; Wyvekens, N.; Khayter, C.; Iafrate, A. J.; Le, L. P.; Aryee, M. J.; Joung, J. K. GUIDE-seq enables genome-wide profiling of off-target cleavage by CRISPR-Cas nucleases. Nat Biotechnol 2015, 33, 187–197.

(61) Marsiglia, J.; Vaalavirta, K.; Knight, E.; Nakamura, M.; Cong, L.; Hughes, N. W. Protein language model-guided engineering of an anti-CRISPR protein for precise genome editing in human cells. Cell Rep. Methods 2024, 4, 100882.

(62) Sharrar, A.; Arake de Tacca, L.; Meacham, Z.; Staples-Ager, J.; Collingwood, T.; Rabuka, D.; Schelle, M. Discovery and engineering of AiEvo2, a novel Cas12a nuclease for human gene editing applications. J. Biol. Chem. 2024, 300, 105685.

(63) Lee, H.-M.; Ren, J.; Tran, K. M.; Jeon, B.-M.; Park, W.-U.; Kim, H.; Lee, K. E.; Oh, Y.; Choi, M.; Kim, D.-S.; Na, D. Identification of efficient prokaryotic cell-penetrating peptides with applications in bacterial biotechnology. Commun. Biol. 2021, 4, 205.

(64) Li, J.; Li, S.; Du, M.; Song, Z.; Han, H. Nuclear delivery of exogenous gene in mature plants using nuclear location signal and cell-penetrating peptide nanocomplex. ACS Appl. Nano Mater. 2023, 6, 160–170.

(65) Fuchs, S. M.; Raines, R. T. Polyarginine as a multifunctional fusion tag. Protein Sci. 2005, 14, 1538–1544.

(66) Lee, J.; Mir, A.; Edraki, A.; Garcia, B.; Amrani, N.; Lou, H. E.; Gainetdinov, I.; Pawluk, A.; Ibraheim, R.; Gao, X. D.; Liu, P.; Davidson, R.; Maxwell, K. L.; Sontheimer, E. J. Potent Cas9 inhibition in bacterial and human cells by AcrIIC4 and AcrIIC5 anti-CRISPR proteins. mBio 2018, 9, e02321–18.

(67) Song, G.; Zhang, F.; Tian, C.; Gao, X.; Zhu, X.; Fan, D.; Tian, Y. Discovery of potent and versatile CRISPR–Cas9 inhibitors engineered for chemically controllable genome editing. Nucleic Acids Res. 2022, 50, 2836–2853.

(68) Schneider, A. F. L.; Wallabregue, A. L. D.; Franz, L.; Hackenberger, C. P. R. Targeted subcellular protein delivery using cleavable cyclic cell-penetrating peptides. Bioconjugate Chem. 2019, 30, 400–404.

(69) Chen, F.-J.; Gao, J. Fast cysteine bioconjugation chemistry. Chem. Eur. J. 2022, 28, e202201843.

(70) Yang, D.; Liu, B.; Sha, H. Advances and prospects of cell-penetrating peptides in tumor immunotherapy. Sci. Rep. 2025, 15, 3392.

