## Supporting Information for "Nuclear Localization Signals Enable the Cellular Delivery of an Anti-CRISPR Protein to Control Genome Editing"

#### Table of Contents

**Materials and Methods**
Conditions ........ S4
Recombinant DNA ........ S4
Protein Purification ........ S4
Cell Culture ........ S4
Acr Addition/Delivery to Cells ........ S5
*GFP*-Disruption Assay ........ S5
*GFP*-Disruption Assay (3D Cell Cultures) ........ S6
HiBiT Knock-In Assay (HUES 8 Cells) ........ S6
HiBiT Knock-In Assay (HEK293T Cells) ........ S6
Next-Generation Sequencing ........ S7
Direct Visualization of Anti-CRISPR–Mediated Cas9–GFP Inhibition ........ S7
Fluorescently Labeling Anti-CRISPR Proteins ........ S7
Direct Visualization of Anti-CRISPR Delivery ........ S8

**Supplementary Tables**
Table S1 ........ S9
Table S2 ........ S9
Table S3 ........ S10
Table S4 ........ S10
Table S5 ........ S11
Table S6 ........ S11

Table S7 .......... S11
Table S8 .......... S11
Table S9 .......... S12
Table S10 .......... S13
Table S11 .......... S14

#### Supplementary Figures

Figure S1 .......... S14
Figure S2 .......... S15
Figure S3 .......... S16
Figure S4 .......... S17
Figure S5 .......... S18
Figure S6 .......... S19
Figure S7 .......... S20
Figure S8 .......... S21
Figure S9 .......... S22
Figure S10 .......... S23
Figure S11 .......... S24
Figure S12 .......... S25
Figure S13 .......... S26
Figure S14 .......... S27
Figure S15 .......... S28
Figure S16 .......... S29
Figure S17 .......... S30
Figure S18 .......... S31
Figure S19 .......... S32
Figure S20 .......... S33
Figure S21 .......... S34
Figure S22 .......... S35
Figure S23 .......... S36
Figure S24 .......... S37
Figure S25 .......... S43
Figure S26 .......... S44
Figure S27 .......... S45
Figure S28 .......... S46
Figure S29 .......... S47
Figure S30 .......... S48
Figure S31 .......... S50
Figure S32 .......... S51
Figure S33 .......... S52
Figure S34 .......... S53
Figure S35 .......... S54

**Plasmid Sequence**
pAV13-MBP-TEV-4×NLS-Cys-AcrIIA4-2×NLS ..... S55

**Equations** ..... S58

**References** ..... S59

### Materials and Methods

#### Conditions

All procedures were performed in air at ambient temperature (~22 °C) and pressure (1.0 atm) unless indicated otherwise.

#### Recombinant DNA

A DNA fragment encoding the sequence of Cys-AcrIIA4 was inserted downstream of a 6×His tag, maltose binding protein (MBP), a TEV protease cleavage site, and a 4×SV40 NLS tag and upstream of a C-terminal 2×SV40 NLS tag in a custom vector (Addgene plasmid #88917)[1] by Gibson assembly to generate a plasmid for bacterial production and purification of 6×NLS-Acr (pAV13-MBP-TEV-4×NLS-Cys-AcrIIA4-2×NLS).

#### Protein Purification

pAV14, pAV17, and pAV18 plasmids were used to produce and purify $LF_N$-Acr, 0×NLS-Acr and 1×NLS-Acr, as previously described.[2] 6×NLS-Acr was purified using the following method: *Escherichia coli* Rosetta2 (DE3) cells transformed with pAV13 were cultured in Lysogeny Broth supplemented with 100 µg/mL ampicillin and 2% w/v glucose at 37 °C for 16 h. The starter culture was subcultured in Terrific Broth supplemented with 100 µg/mL ampicillin and 2% w/v glucose at 37 °C until the $OD_{600}$ was 0.6–0.8. Protein production was induced with 500 µM isopropyl-β-D-thiogalactoside (IPTG), and the cultures were grown at 16 °C for 16 h. The cells were harvested, resuspended in lysis buffer (20 mM Tris–HCl buffer, pH 7.5, containing 500 mM NaCl, 20 mM imidazole, 1 mM tris(2-carboxyethyl)phosphine (TCEP), and 5% v/v glycerol) supplemented with 0.5 mM phenylmethanesulfonyl fluoride and Roche cOmplete Protease Inhibitor Cocktail, and lysed by sonication. MBP–6×NLS-Acr was purified using Ni-NTA Superflow resin (Qiagen, 30410) in lysis buffer supplemented with either 20 mM imidazole (wash) or 300 mM imidazole (elution). Eluted MBP–6×NLS-Acr was cleaved with TEV protease (purified in-house using Addgene plasmid #8827)[3] overnight at 4 °C and dialyzed into 20 mM HEPES buffer, pH 7.5, containing 300 mM NaCl, 1 mM TCEP, and 5% v/v glycerol. Cleaved 6×NLS-Acr was loaded onto an MBPTrap HP (Cytiva, 28918779) upstream of a HiTrap SP HP (Cytiva, 17115201). 6×NLS-Acr was eluted over a linear gradient of NaCl (0.3–1.00 M) in 20 mM HEPES–HCl buffer, pH 7.5, containing 1 mM TCEP and 5% v/v glycerol. 6×NLS-Acr was then loaded onto a HiLoad 16/600 Superdex 200 pg (Cytiva, 28989335) and eluted with 20 mM HEPES–HCl buffer, pH 7.5, containing 500 mM NaCl, 1 mM TCEP, and 5% v/v glycerol. Acr concentration was measured using $A_{280\ nm}$ and a BCA assay kit (Thermo Fisher Scientific). Acr purity was assessed by SDS–PAGE (Coomassie blue staining) and Q-TOF LC–MS.

#### Cell Culture

U2OS cells, U2OS cells with an integrated *GFP–PEST* fusion gene (U2OS-EFFP.PEST), U2OS cells stably expressing Cas9–GFP and 53BP1–mCherry (U2OS-Cas9-GFP), and HEK293T cells were cultured in Dulbecco's modified Eagle's medium with GlutaMax (DMEM, Gibco, 10564-011) supplemented with 10% v/v fetal bovine serum (FBS, Corning, 35-016-CV), 100 U/mL penicillin−streptomycin (Gibco, 15140-122), and 1 mM sodium pyruvate (Gibco, 11360-070) at

37 °C in a 5% v/v $CO_2$(g) atmosphere. INS-1E rat β-cells were cultured in Roswell Park Memorial Institute (RPMI) medium (Gibco, 11875-093) supplemented with 10% FBS, 1 mM pyruvate, 100 U/mL penicillin-streptomycin, and 50 µM of 2-mercaptoethanol. FBS was never removed from the medium for Acr delivery experiments in U2OS cell lines, INS-1E rat β-cells, and HEK293T cells. HUES 8 cells (Harvard Stem Cell Institute iPS Core Facility) were cultured in mTeSR1 medium (STEMCELL Technologies, 85850) over a Matrigel-coated surface (Corning, 356234) at 37 °C in a 5% v/v $CO_2$(g) atmosphere. mTeSR1 medium was supplemented with 10 µM ROCK inhibitor Y-27632 (Enzo Life Sciences, ALX-270-333) for cell splitting and genome editing experiments.

#### Acr Addition/Delivery to Cells

At a 0-h dosing time or time point, Cas9-transfected cells were treated with Acrs immediately before starting the incubation at 37 °C. At a >0-h dosing time or time point, Cas9-transfected cells were treated with Acrs >0 h after starting the incubation at 37 °C. Cas9 transfection refers to delivering Cas9 into cells either via nucleofection or lipofection. For dose-dependency studies at 2 h in the *GFP* disruption assay, Acrs (0.125–4 µM) were added to the cells 2 h after Cas9 transfection by replacing the cell culture medium in the well with fresh medium containing the Acrs diluted to the desired concentration. The final buffer concentration in the medium was <1% v/v at all times. For dose-dependency studies at 0 h or for time-dependency studies at a constant concentration, Acrs (0.125–4 µM) were added to the cells 0–24 h (excluding 2 h) after Cas9 transfection by directly pipetting an Acr stock solution in buffer to the cell culture medium in the well. The Acr stock solution was prepared so that, upon dilution, the final buffer concentration in the medium was 4.8% v/v (e.g., adding 5 µL of Acr stock solution to 100 µL of cell culture medium).

#### *GFP*-Disruption Assay

SpCas9-NLS RNP (20 pmol) targeting the *GFP* gene was delivered via nucleofection into U2OS-EGFP.PEST cells (300,000 cells) using an SE Cell Line 4D-Nucleofector X Kit (Lonza, V4SC-1096, DN-100 protocol). The nucleofected cells were resuspended in cell culture medium (100,000 cells/mL), transferred to a Revvity (6055302) 96-well PhenoPlate (100 µL), and incubated at 37 °C. 0–24 h after starting the incubation (dosing time), 0×NLS-Acr, 1×NLS-Acr, or 6×NLS-Acr were added to the cells via the medium-exchange (2-h dosing time) or direct addition method (all other dosing times). After 48 h, live cells were stained with 2 µg/mL Hoechst 33342 (37 °C, 20 min) in FluoroBrite DMEM (Gibco, A1896701) and imaged using high-throughput confocal microscopy on a Revvity Opera Phenix microscope equipped with a 20× objective, which enabled detection of GFP (green) and Hoechst 33342 (blue) fluorescence upon 488 nm and 405 nm laser excitation. Digital phase contrast (DPC) images were acquired using the same microscope. Images were analyzed using Revvity Harmony Software. Cells were selected based on the Hoechst 33343 nuclear stain, and the cytosol region was defined using digital phase contrast. A mean GFP threshold intensity was chosen by comparing ApoCas9- and Cas9 RNP-nucleofected cells. The distribution of cells above this threshold (GFP positive) was used to determine % *GFP* disruption. Cas9-mediated cleavage disrupts the *GFP* gene. Acr delivery prevents Cas9-mediated *GFP* disruption.

#### *GFP*-Disruption Assay (3D Cell Cultures)

SpCas9-NLS RNP (20 pmol) targeting the *GFP* gene was delivered via nucleofection into U2OS-EGFP.PEST cells (300,000 cells) using an SE Cell Line 4D-Nucleofector X Kit (Lonza, V4SC-1096, DN-100 protocol). The nucleofected cells were resuspended in cell culture medium (850,000 cells/mL), transferred to a Revvity (6055330) CellCarrier Spheroid ULA 96-well plate (100 µL), and incubated at 37 °C. 7–13 h after starting the incubation (dosing time), 6×NLS-Acr (2.5 µM) was added to the cells by the direct addition of a concentrated 6×NLS-Acr solution (5 µL). After 48 h, live cells were imaged using high-throughput confocal microscopy on a Revvity Opera Phenix microscope equipped with a 5× objective, which enabled detection of GFP (green) fluorescence upon 488 nm laser excitation. Brightfield images were acquired using the same microscope. Live cell imaging was also performed before 6×NLS-Acr addition at 7 and 13 h. The 48-h images were analyzed using Revvity Harmony Software. The spheroid region was defined based on residual GFP fluorescence and partitioned into an outer, entire, and inner spheroid region (Figure S15). The mean GFP intensity in each region was used to determine % *GFP* disruption. Cas9-mediated cleavage disrupts the *GFP* gene. Successful delivery of 6×NLS-Acr prevents Cas9-mediated *GFP* disruption.

#### HiBiT Knock-In Assay (HUES 8 Cells)

SpCas9-NLS RNP (20 pmol) targeting the *GAPDH* gene and the HiBiT ssODN (80 pmol) were delivered via nucleofection into HUES 8 cells (450,000 cells) using a P3 Primary Cell 4D-Nucleofector X Kit (Lonza, V4XP-3032, CA-137 protocol). The nucleofected cells were resuspended in mTeSR1 medium supplemented with 10 µM Y-27632 (420,000 cells/mL), transferred to a Matrigel-coated 96-well plate (100 µL), and incubated at 37 °C. Cells were treated with 0×NLS-Acr or 6×NLS-Acr (0.125–4 µM) 0–16 h after starting the incubation (dosing time). After 24 h, the cells were incubated with the PrestoBlue HS reagent (Thermo Fisher Scientific, P50200), and their viability was measured using fluorescence spectroscopy (560 nm excitation/590 nm emission, SpectraMax M5). The cells were lysed in the presence of LgBiT and furimazine (Nano-Glo HiBiT Lytic Detection System, Promega, N3030), and their luminescence was quantified (Perkin Elmer EnVison). Cas9-mediated cleavage and HiBiT tag insertion via homology-directed repair produce NanoLuc after adding LgBiT and furimazine. Acr delivery prevents Cas9-mediated cleavage, HiBiT tag insertion, and NanoLuc formation.

#### HiBiT Knock-In Assay (HEK293T Cells)

SpCas9-NLS RNP (20 pmol) targeting the *GAPDH* gene and the HiBiT ssODN (80 pmol) were delivered via nucleofection into HEK293T cells (300,000 cells) using an SF Cell Line 4D-Nucleofector X Kit (Lonza, V4SC-2096, CM-130 protocol). The nucleofected cells were resuspended in cell culture medium (150,000 cells/mL), transferred to a poly(D-lysine)-coated 96-well plate (100 µL), and incubated at 37 °C. Cells were treated with 0×NLS-Acr or 6×NLS-Acr (0.4–1.5 µM) at 0 h. After 72 h, the cells were incubated with the PrestoBlue HS reagent (Thermo Fisher Scientific, P50200), and their viability was measured using fluorescence spectroscopy (560 nm excitation/590 nm emission, SpectraMax M5). The cells were lysed in the presence of LgBiT and furimazine (Nano-Glo HiBiT Lytic Detection System, Promega, N3030), and their luminescence was quantified (Perkin Elmer EnVison). Cas9-mediated cleavage and HiBiT tag insertion via homology-directed repair produce NanoLuc after adding LgBiT and furimazine. Acr delivery prevents Cas9-mediated cleavage, HiBiT tag insertion, and NanoLuc formation.

#### Next-Generation Sequencing

HEK293T cells were seeded at 150,000 cells per well in 571.4 µL of cell culture medium on a poly(D-lysine)-coated 24-well plate and incubated at 37 °C. After 24 h, the cells were transfected via lipofection (Lipofectamine 3000, Invitrogen, L3000-015) with plasmids encoding 2×NLS-SpCas9 (750 ng, Addgene plasmid #42230)[4] and *EMX1*-targeting gRNA (250 ng).[5] Cells were treated with 0×NLS-Acr (4 µM), 6×NLS-Acr (4 µM), or $LF_N$-Acr (250 nM) ± PA (20 nM) at 1 h. After 72 h, genomic DNA was extracted (QIAamp DNA Mini Kit, Qiagen, 51304) and the target and off-target sites were PCR amplified, barcoded, and purified by gel extraction (MinElute Gel Extraction Kit, Qiagen, 28604). The amplicons were sequenced with NGS and analyzed using the CRISPResso2 software pipeline to determine % modification.

#### Direct Visualization of Anti-CRISPR–Mediated Cas9–GFP Inhibition

U2OS cells stably expressing Cas9–GFP and 53BP1–mCherry fusion proteins (U2OS-Cas9-GFP cells) were seeded at 100,000 cells per well in 100 µL of cell culture medium on a 10 mm microwell inside a 35 mm glass-bottom dish. The dish was then covered with 1.9 mL of cell culture medium and incubated at 37 °C. After 24 h, the cells were transfected via lipofection (RNAiMAX, Invitrogen, 13778-030) with a truncated gRNA that targets a highly repetitive region on chromosome 3 (Ch3 gRNA). Cells were treated with 0×NLS-Acr (2.5 µM), 6×NLS-Acr (2.5 µM), or $LF_N$-Acr (2.5 µM) ± PA (20 nM) either 0.5 h before (−0.5 h) or 15 h after (15 h) gRNA transfection. Before imaging, live cells were washed with Leibovitz's L-15 medium supplemented with 10% v/v FBS and 100 U/mL penicillin-streptomycin and maintained in this medium. Cells were imaged using widefield fluorescence microscopy at 15 and 20 h for −0.5- and 15-h Acr treatments, respectively. A Nikon Ti-E fluorescence microscope, equipped with a 60× objective and a stage-top incubator at 37 °C, was used to detect Cas9–GFP and 53BP1–mCherry fluorescence upon 485 nm and 560 nm laser excitation. Emission was collected using 525/50 nm and 630/75 nm filters, respectively. Z-stacks (25 slices, 0.5 µm step size) encompassing the entire nuclear volume were acquired using dual Hamamatsu C9100-13 EMCCD cameras. Image analysis was performed using a custom pipeline in a Python-based Jupyter notebook. In brief, nuclei were segmented from the 53BP1–mCherry channel using a voxel-aware watershed algorithm (skimage.segmentation.watershed) after generating seeds from the local maxima of a Gaussian-filtered image (sigma = 2 pixels). Cas9–GFP foci were identified in 3D by first applying a rolling-ball background subtraction (ball radius = 5 pixels) followed by white-tophat filtering (ball radius = 3 pixels) to enhance punctate features. Candidate foci were then localized using the DAOStarFinder class from Photutils with a detection threshold of 5 standard deviations above the local background and a full-width at half-maximum (FWHM) of 3 pixels. Finally, the integrated intensity of each focus was measured using an ellipsoidal aperture (semi-major axis = 3 pixels), with local background subtraction from a concentric ring (inner radius = 4 pixels, outer radius = 6 pixels). All data were collated and analyzed using the pandas library.

#### Fluorescently Labeling Anti-CRISPR Proteins

**Synthesis of Cy5-0×NLS-Acr.** An 8.3 mM stock solution of sulfo-Cyanine5 (sulfo-Cy5) maleimide (Lumiprobe, 23380) was prepared in water. 14.5 μL of the 8.3 mM sulfo-Cy5 maleimide stock solution (14.7 equiv) were added to 40 μL of 204 μM 0×NLS-Acr (1 equiv) in 20 mM Tris–HCl buffer, pH 7.5, containing 150 mM NaCl, 1 mM TCEP, and 5% v/v glycerol. The reaction mixture was gently shaken (80 RPM) for 30 min at 22 °C. Afterward, the reaction mixture was passed through a 0.5 mL Zeba spin desalting column (7 kDa MWCO), and the buffer was exchanged to 20 mM HEPES–HCl buffer, pH 7.5, containing 500 mM NaCl, 1 mM TCEP, and 5% v/v glycerol.

**Synthesis of Cy5-6×NLS-AcrIIA4.** An 8.3 mM stock solution of sulfo-Cy5 maleimide was prepared in water. 4 μL of the 8.3 mM sulfo-Cy5 maleimide stock solution (6.9 equiv) were added to 27.8 μL of 172 μM 6×NLS-Acr (1 equiv) in 20 mM HEPES–HCl buffer, pH 7.5, containing 500 mM NaCl, 1 mM TCEP, and 5% v/v glycerol. The reaction mixture was gently shaken (80 RPM) for 30 min at 22 °C. Afterward, the reaction mixture was passed through a 0.5 mL Zeba spin desalting column (7 kDa MWCO), and the buffer was exchanged to 20 mM HEPES–HCl buffer, pH 7.5, containing 500 mM NaCl, 1 mM TCEP, and 5% v/v glycerol.

**Synthesis of Cy5-LF$_N$-Acr.** A 6.2 mM stock solution of sulfo-Cy5 maleimide was prepared in water. 10 μL of the 6.2 mM sulfo-Cy5 maleimide stock solution (11.3 equiv) were added to 45 μL of 122 μM LF$_N$-Acr (1 equiv) in 20 mM Tris–HCl buffer, pH 7.5, containing 150 mM NaCl, 1 mM TCEP, and 5% v/v glycerol. The reaction mixture was gently shaken (80 RPM) for 30 min at 22 °C. Afterward, the reaction mixture was purified using a Pierce Microdyalisis Plate (10 kDa MWCO), and the buffer was exchanged to 20 mM Tris–HCl buffer, pH 7.5, containing 150 mM NaCl, 1 mM TCEP, and 5% v/v glycerol.

#### Direct Visualization of Anti-CRISPR Delivery

U2OS cells were seeded in a 96-well plate at a density of 10,000 cells per well in 100 μL of culture medium and incubated at 37 °C for approximately 18 h. INS-1E rat β-cells were seeded in an ECM-coated 96-well plate at a density of 75,000 cells per well in 100 μL of culture medium and incubated at 37 °C for approximately 24 h. Cells were then treated with Cy5-0×NLS-Acr (1.5–4 μM), Cy5-6×NLS-Acr (1.5–4 μM), and Cy5-LF$_N$-Acr (2.5 μM) ± PA (20 nM), and incubated for 5 min to 2 h at 37 °C. Cells were washed with FluoroBrite DMEM. Cells were stained with Hoechst 33342 (4 μg/mL, 10 min, 37 °C) and WGA555 (2.5 μg/mL, 5 min, 22 °C) in FluoroBrite DMEM, then imaged using confocal microscopy (60× objective) to detect fluorescence from Hoechst 33342 (blue; 405 nm excitation), WGA555 (yellow; 568 nm excitation), and Cy5 (red; 647 nm excitation). Digital phase contrast (DPC) microscopy was also performed.

### Supplementary Tables

##### Table S1. Compilation of $IC_{50}$ and Efficacy Values

| Inhibitor | Dosing time (h) | End point (h) | Assay | Cell type | Figure | $IC_{50}$ (nM)[a] | Efficacy (%)[a] |
|---|---|---|---|---|---|---|---|
| 6×NLS-Acr | 0 | 48 | *GFP* | U2OS-EGFP | 2B, S12 | 470 | 96 |
| 6×NLS-Acr | 2 | 48 | *GFP* | U2OS-EGFP | 2B | 1,460 | 86 |
| $LF_N$-Acr | 0 | 48 | *GFP* | U2OS-EGFP | S12 | 2.9 | 86 |
| 6×NLS-Acr | 0 | 24 | HiBiT | HUES 8 | S20 | 630 | 99 |
| $LF_N$-Acr | 0 | 24 | HiBiT | HUES 8 | S20 | 3.1 | 88 |

[a]The "Absolute $IC_{50}$, X is log(concentration)" function in GraphPad Prism software was used to fit the data from the *GFP*-disruption and *HiBiT*-knock-in assays to determine $IC_{50}$ and maximum Cas9 inhibition values. The maximum Cas9 inhibition value or efficacy was calculated as 100 minus the 'Bottom' value of the fitted curve, where 'Bottom' represents the baseline Cas9 activity, which approaches zero with increasing Acr concentration.

##### Table S2. Cell Lines Used in This Study

| Cell Line | Purpose | Reference |
|---|---|---|
| NEB® 5-alpha Competent *E. coli* (High Efficiency) | Cloning bacterial expression vectors | NEB |
| Rosetta 2(DE3) Competent Cells (Novagen) | Producing and purifying Acrs | MilliporeSigma |
| NEB® Stable Competent *E. coli* (High Efficiency) | Cloning mammalian expression vectors | NEB |
| U2OS-EGFP.PEST | *GFP*-disruption assay | Reyon et al. (2012)[6] |
| HUES 8 (RRID:CVCL_B207) | HiBiT-knock-in assay | Harvard Stem Cell Institute (HSCI) iPS Core Facility; Cowan et al. (2004)[7] |
| HEK293T | HiBiT-kock-in, T7E1, and NGS assays | ATCC |
| U2OS-Cas9-GFP | Direct visualization of the inhibition of Cas9–GFP | Liu et al. (2020)[8] |
| U2OS | Direct visualization of Acr delivery | ATCC |
| INS-1E rat β-cells | Direct visualization of Acr delivery | Millipore Sigma |

**Table S3. Proteins Used in This Study**

| Protein | Source | Reference |
|---|---|---|
| 6×NLS-Acr | Purified in-house | This work |
| 0×NLS-Acr | Purified in-house | Vera et al. (2025)[2] |
| 1×NLS-Acr | Purified in-house | Vera et al. (2025)[2] |
| $LF_N$-Acr | Purified in-house | Vera et al. (2025)[2] |
| Protective Antigen | Purified in-house | Pomerantsev et al. (2021)[9] |
| SpCas9-NLS | GenScript | Cat. No. Z03385 |
| 2×NLS-LbCas12a | NEB | Cat. No. M0653T |
| TEV protease | Purified in-house | Kapust et al. (2001)[3] |

**Table S4. Plasmids Used in This Study**

| Plasmid name | Purpose | Reference |
|---|---|---|
| 4xNLS-pMJ915v2 | Amplifying the backbone used to clone pAV13 | Staahl et al. (2017)[1] Addgene #88917 |
| pAV13-MBP-TEV-4×NLS-Cys-AcrIIA4-2×NLS | Bacterial production and purification of 6×NLS-Acr | This work |
| pAV17-MBP-TEV-Cys-AcrIIA4 | Bacterial production and purification of 0×NLS-Acr | Vera et al. (2025)[2] |
| pAV18-MBP-TEV-Cys-AcrIIA4-NLS | Bacterial production and purification of 1×NLS-Acr | Vera et al. (2025)[2] |
| pAV14-MBP-TEV-$LF_N$-Cys-AcrIIA4-NLS | Bacterial production and purification of $LF_N$-Acr | Vera et al. (2025)[2] |
| pYS5-PA BH500 | Bacterial production and purification of PA | Singh et al. (1989)[10] |
| pRK793 | Bacterial production and purification of TEV protease | Kapust et al. (2001)[3] Addgene #8827 |
| pX330-U6-Chimeric_BB-CBh-hSpCas9 | Mammalian production of 2×NLS-SpCas9 | Cong et al. (2013)[4] Addgene #42230 |
| JDS246 | Mammalian production of SpCas9-NLS | Addgene #43861 |
| pEMX1-gRNA | Mammalian production of EMX1 gRNA | Sreekanth et al. (2020)[5] |

**Table S5. Primer Sequences for gRNA Cloning and Synthesis for RNP Experiments**

| Primer | Sequence[a] |
|---|---|
| GFP Fwd | TAATACGACTCACTATA**GGTGGTGCAGATGAACTTCA**GTTTTAGAGCTAGAAAT |
| GAPDH Fwd | TAATACGACTCACTATA**GGTCCAGGGGTCTTACTCCT**GTTTTAGAGCTAGAAAT |
| Universal Rev | AAAAGCACCGACTCGGTGCCACTTTTTCAAGTTGATAACGGACTAGCCTTATTTT AACTTGCTATTTCTAGCTCTAAAAC |

[a]The spacer sequence is in bold typeface.

**Table S6. Sequence[a] of the LbCas12a crRNA Used in the *GFP*-Disruption Assay**

| | |
|---|---|
| LbCas12a crRNA | UAAUUUCUACUAAGUGUAGAU**CGUCGCCGUCCAGCUCGACCAGG** |

[a]The spacer sequence is in bold typeface. LbCas12a crRNA was ordered from IDT.

**Table S7. ssODN Sequence**

| | |
|---|---|
| GAPDH HiBiT | TCTTCTAGGTATGACAACGAATTTGGCTACAGCAACAGGGTGGTGGACCTCATGGCCCACA TGGCCTCCAAGGAGGTGAGCGGCTGGCGGCTGTTCAAGAAGATTAGCTAAGACCCCTGGAC CACCAGCCCCAGCAAGAGCACAAGAGGAAGAGAGAGACCCTCACTGCTGGGGAGTCCCTGC |

**Table S8. Spacer Sequence of Plasmid-Encoded EMX1 gRNA**

| | |
|---|---|
| EMX1 | GAGTCCGAGCAGAAGAAGAA |

**Table S9. Primers Used to Generate Amplicons for Next-Generation Sequencing**

| | Target | PAM | Forward Primer | Reverse Primer |
|---|---|---|---|---|
| EMX1 On | GAGTCCGAGCAGAAGAAGAA | GGG | ACACTCTTTCCCTACACGACGCTCTTCCGATCTNNNNCAGCTCAGCCTGAGTGTTGA | TGGAGTTCAGACGTGTGCTCTTCCGATCTCTCGTGGGTTTGTGGTTGC |
| EMX1 OT1 | GAGTTAGAGCAGAAGAAGAA | AGG | ACACTCTTTCCCTACACGACGCTCTTCCGATCTNNNNTTCTGAGGGCTGCTACCTGT | TGGAGTTCAGACGTGTGCTCTTCCGATCTGCCCAATCATTGATGCTTTT |
| EMX1 OT2 | GAGTCTAAGCAGAAGAAGAA | GAG | ACACTCTTTCCCTACACGACGCTCTTCCGATCTNNNNCACGGCCTTTGCAAATAGAG | TGGAGTTCAGACGTGTGCTCTTCCGATCTGGCTTTCACAAGGATGCAGT |
| EMX1 OT3 | AAGTCTGAGCACAAGAAGAA | TGG | ACACTCTTTCCCTACACGACGCTCTTCCGATCTNNNNGTTCTGACATTCCTCCTGAGGGA | TGGAGTTCAGACGTGTGCTCTTCCGATCTATGGCTTACATATTTATTAGATAAAATGTATTCC |
| EMX1 OT4 | GAGTCCTAGCAGGAGAAGAA | GAG | ACACTCTTTCCCTACACGACGCTCTTCCGATCTNNNNCCAGACTCAGTAAAGCCTGGA | TGGAGTTCAGACGTGTGCTCTTCCGATCTTGGCCCCAGTCTCTCTTCTA |

**Table S10. Amplicon Sequences**

| Target | Sequence |
|---|---|
| EMX1 On | CAGCTCAGCCTGAGTGTTGAGGCCCCAGTGGCTGCTCTGGGGGCCTCCTGAGTTTCTC<br>ATCTGTGCCCCTCCCTCCCTGGCCCAGGTGAAGGTGTGGTTCCAGAACCGGAGGACAA<br>AGTACAAACGGCAGAAGCTGGAGGAGGAAGGGCCTGAGTCCGAGCAGAAGAAGAAGGG<br>CTCCCATCACATCAACCGGTGGCGCATTGCCACGAAGCAGGCCAATGGGGAGGACATC<br>GATGTCACCTCCAATGACTAGGGTGGGCAACCACAAACCCACGAG |
| EMX1 OT1 | TTCTGAGGGCTGCTACCTGTACATCTGCACAAGATTGCCTTTACTCCATGCCTTTCTT<br>CTTCTGCTCTAACTCTGACAATCTGTCTTGCCATGCCATAAGCCCCTATTCTTTCTGT<br>AACCCCAAGATGGTATAAAAGCATCAATGATTGGGC |
| EMX1 OT2 | CACGGCCTTTGCAAATAGAGCCCTTTATTCATAGTAGACAAGAGTCTAAGCAGAAGAA<br>GAAGAGAGCCACTACCCAACCATCTACTCTTCTAATGGTGTTTTCCTACAAAGGCCAA<br>GTCATGAGACTGCATCCTTGTGAAAGCC |
| EMX1 OT3 | GTTCTGACATTCCTCCTGAGGGAAAATAAATAAATTAATTAAAAATATATATATATAT<br>GTATAATGATAAACATGCTAACAAAGTCTGAGCACAAGAAGAATGGTGAGAAGGAATA<br>CATTTTATCTAATAAATATGTAAGCCAT |
| EMX1 OT4 | CCAGACTCAGTAAAGCCTGGAGGCTGCCAGGTAGGGCTGGGGCCAGCATGACCTGAGT<br>CCTAGCAGGAGAAGAAGAGGCAGCCTAGAGTCTTCTGTGAAGTGCACATAGAAGAGAG<br>ACTGGGGCCA |

**Table S11. Sequence[a] of Truncated Ch3Rep gRNA**

| | |
|---|---|
| Ch3Rep gRNA | G**UGAUAUCACAG**GUUUUAGAGCUAGAAAUAGCAAGUUAAAAUAAGGCUAGUCCGUUAUCA<br>ACUUGAAAAAGUGGCACCGAGUCGGUGCUUUU |

[a]The truncated spacer sequence is in bold typeface.

#### Supplementary Figures

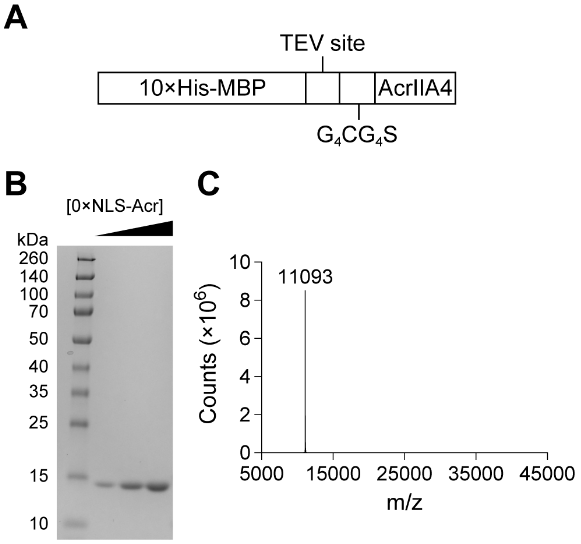


**Figure S1.** Purification of 0×NLS-Acr. (A) The construct that was used to produce and purify 0×NLS-Acr (pAV17-MBP-TEV-Cys-AcrIIA4). A protein fusion between a polyhistidine-tagged maltose-binding protein (10×His-MBP, for solubility and purification), a TEV protease cleavage site, a glycine–cysteine–serine linker ($G_4CG_4S$, for flexibility and optional bioconjugation), and AcrIIA4 (for Cas9 inhibition). (B) Assessing 0×NLS-Acr purity by SDS–PAGE. The concentration of 0×NLS-Acr increases from left to right. (C) Assessing 0×NLS-Acr purity by Q-TOF LC–MS.

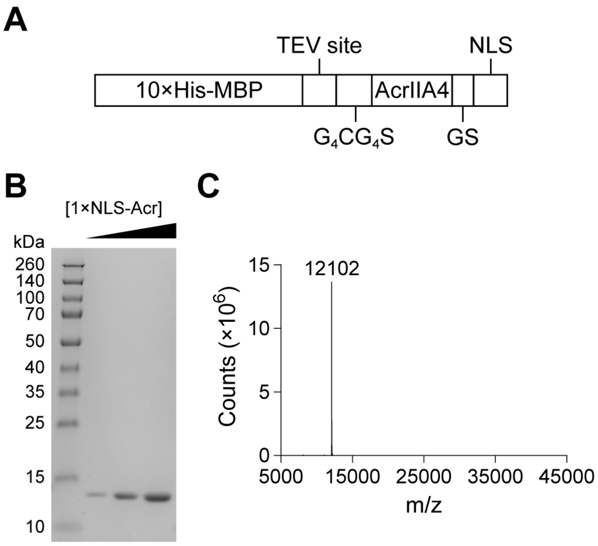


**Figure S2.** Purification of 1×NLS-Acr. (A) The construct that was used to produce and purify 1×NLS-Acr (pAV18-MBP-TEV-Cys-AcrIIA4-NLS). A protein fusion between a 10×His-MBP tag, a TEV protease cleavage site, a $G_4CG_4S$ linker, AcrIIA4, a GS linker (for flexibility), and an SV40 nuclear localization signal (NLS). (B) Assessing 1×NLS-Acr purity by SDS–PAGE. The concentration of 1×NLS-Acr increases from left to right. (C) Assessing 1×NLS-Acr purity by Q-TOF LC–MS.

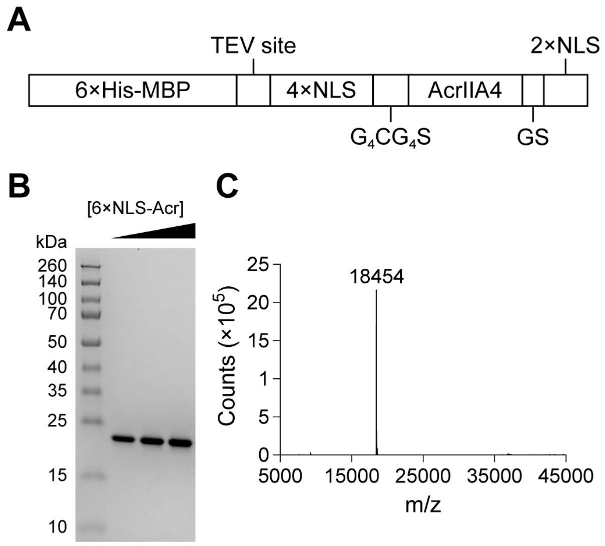


**Figure S3.** Purification of 6×NLS-Acr. (A) The construct that was used to produce and purify 6×NLS-Acr (pAV13-MBP-TEV-6×NLS-AcrIIA4). A protein fusion between a 6×His-MBP tag, a TEV protease cleavage site, a 4×NLS tag (for cellular delivery and nuclear localization), a $G_4CG_4S$ linker, AcrIIA4, a GS linker, and a 2×NLS tag (for cellular delivery and nuclear localization). (B) Assessing 6×NLS-Acr purity by SDS–PAGE. The concentration of 6×NLS-Acr increases from left to right. (C) Assessing 6×NLS-Acr purity by Q-TOF LC–MS.

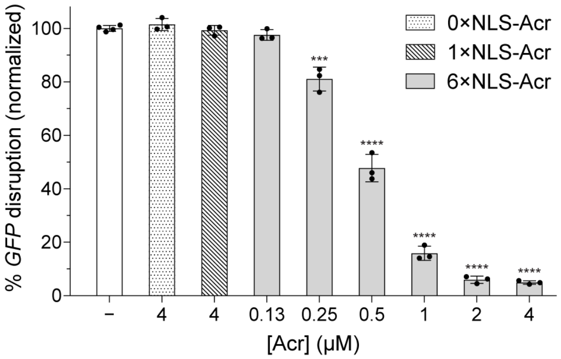


**Figure S4.** Dose-dependent inhibition of Cas9 by 6×NLS-Acr in the *GFP*-disruption assay (0-h dosing time). SpCas9-NLS RNP (20 pmol) targeting *GFP* was delivered via nucleofection into U2OS-EGFP.PEST cells. The cells were seeded, incubated at 37 °C, and treated with 6×NLS-Acr (0.13–4 µM) at 0 h. After 48 h, live cells were stained with Hoechst 33342 and imaged using high-throughput confocal microscopy. Controls include nucleofection of Cas9 RNP (white bar) and nucleofection of Cas9 RNP followed by treatment with 0×NLS-Acr (4 µM) or 1×NLS-Acr (4 µM) at 0 h. The values were normalized to Cas9 RNP and are the mean ± standard deviation (SD) of three independent replicates. The significance of 6×NLS-Acr delivery was determined with an unpaired, two-tailed *t*-test versus Cas9 RNP, where *, **, ***, and **** refer to $P \leq 0.05$, $P \leq 0.01$, $P \leq 0.001$, and $P \leq 0.0001$, respectively.

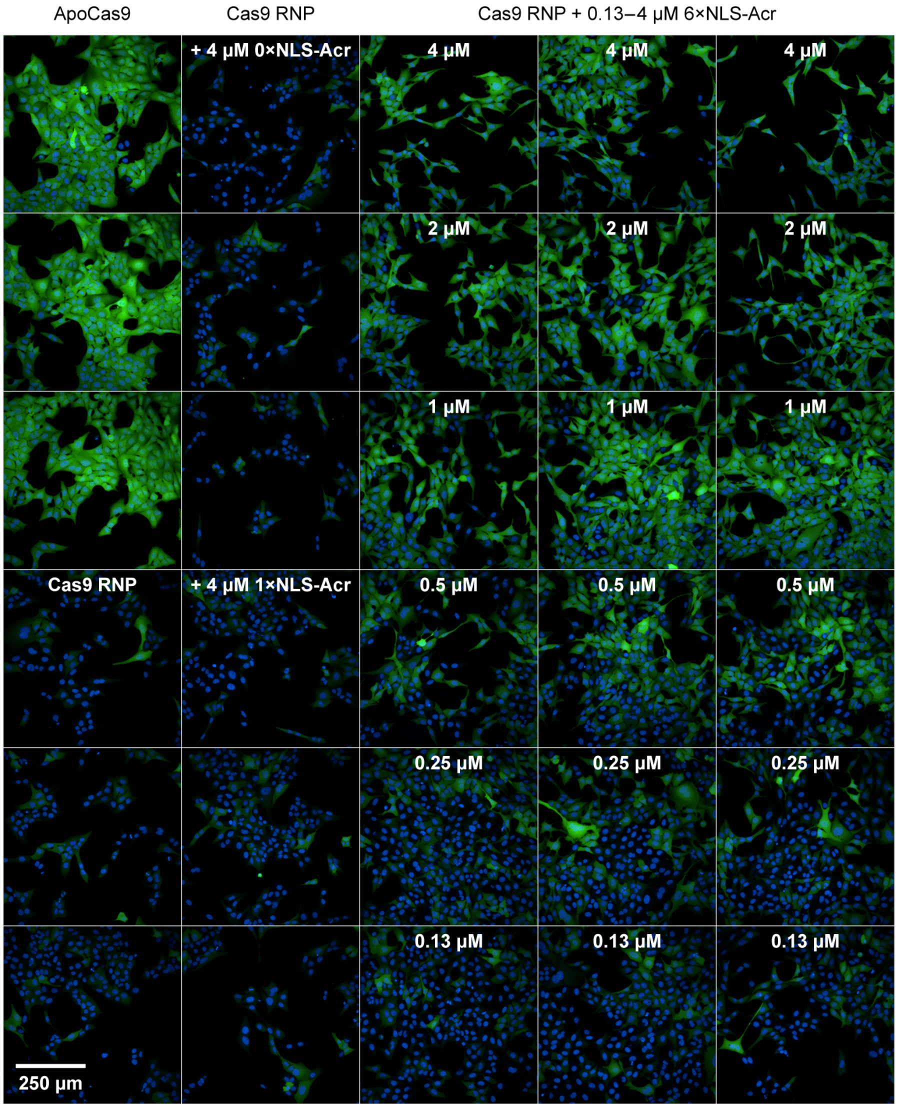


**Figure S5.** Representative fluorescence images from the *GFP*-disruption assay in Figure S4. SpCas9-NLS RNP (20 pmol) targeting *GFP* was delivered via nucleofection into U2OS-EGFP.PEST cells. The cells were seeded, incubated at 37 °C, and treated with 6×NLS-Acr (0.13–4 µM) at 0 h. After 48 h, live cells were stained with Hoechst 33342 (blue, nucleus) and imaged using high-throughput confocal microscopy. Controls include nucleofection of Apo-Cas9, nucleofection of Cas9 RNP, and nucleofection of Cas9 RNP followed by treatment with 0×NLS-Acr (4 µM) or 1×NLS-Acr (4 µM) at 0 h.

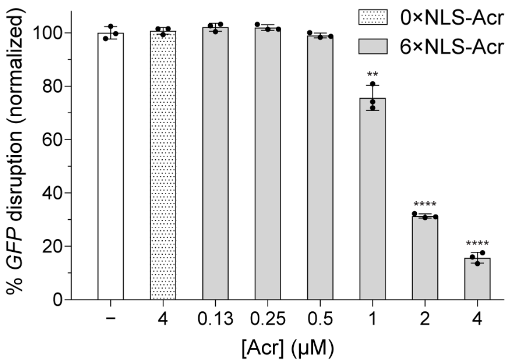


**Figure S6.** Dose-dependent inhibition of Cas9 by 6×NLS-Acr in the *GFP*-disruption assay (2-h dosing time). SpCas9-NLS RNP (20 pmol) targeting *GFP* was delivered via nucleofection into U2OS-EGFP.PEST cells. The cells were seeded, incubated at 37 °C, and treated with 6×NLS-Acr (0.13–4 μM) at 2 h. After 48 h, live cells were stained with Hoechst 33342 and imaged using high-throughput confocal microscopy. Controls include nucleofection of Cas9 RNP (white bar) and nucleofection of Cas9 RNP followed by treatment with 0×NLS-Acr (4 μM) at 2 h. The values were normalized to Cas9 RNP and are the mean ± SD of three independent replicates. The significance of 6×NLS-Acr delivery was determined with an unpaired, two-tailed *t*-test versus Cas9 RNP, where *, **, ***, and **** refer to $P \leq 0.05$, $P \leq 0.01$, $P \leq 0.001$, and $P \leq 0.0001$, respectively.

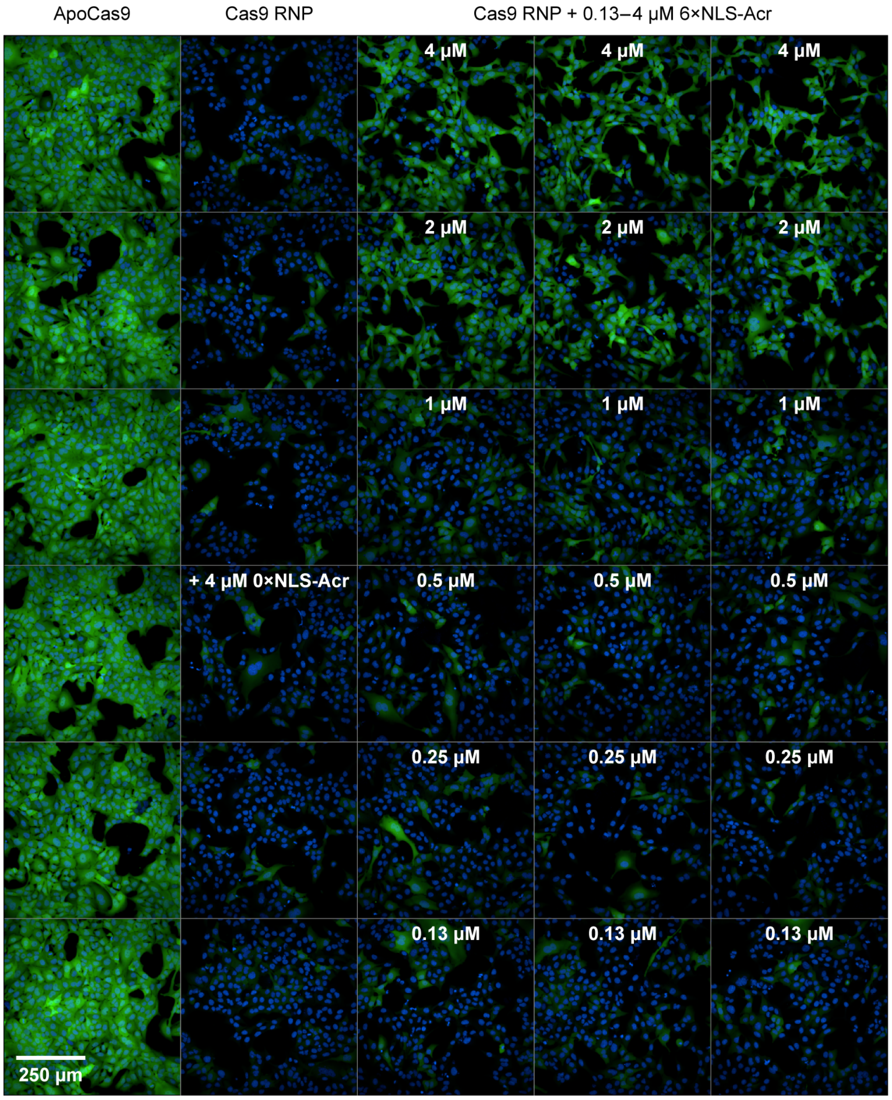


**Figure S7.** Representative fluorescence images from the *GFP*-disruption assay in Figure S6. SpCas9-NLS RNP (20 pmol) targeting *GFP* was delivered via nucleofection into U2OS-EGFP.PEST cells. The cells were seeded, incubated at 37 °C, and treated with 6×NLS-Acr (0.13–4 μM) at 2 h. After 48 h, live cells were stained with Hoechst 33342 (blue, nucleus) and imaged using high-throughput confocal microscopy. Controls include nucleofection of Apo-Cas9, nucleofection of Cas9 RNP, and nucleofection of Cas9 RNP followed by treatment with 0×NLS-Acr (4 μM) at 2 h.

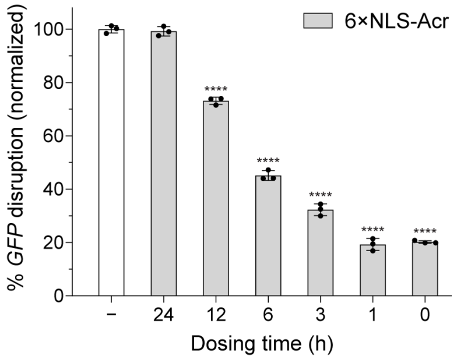


**Figure S8.** Time-dependent inhibition of Cas9 by 6×NLS-Acr in the *GFP*-disruption assay. SpCas9-NLS RNP (20 pmol) targeting *GFP* was delivered via nucleofection into U2OS-EGFP.PEST cells. The cells were seeded, incubated at 37 °C, and treated with 6×NLS-Acr (1 μM), varying the dosing time from 0 to 24 h. After 48 h, live cells were stained with Hoechst 33342 and imaged using high-throughput confocal microscopy. Nucleofection of Cas9 RNP (white bar) serves as the control. The values were normalized to Cas9 RNP and are the mean ± SD of three independent replicates. The significance of 6×NLS-Acr delivery was determined with an unpaired, two-tailed *t*-test versus Cas9 RNP, where *, **, ***, and **** refer to $P \leq 0.05$, $P \leq 0.01$, $P \leq 0.001$, and $P \leq 0.0001$, respectively.

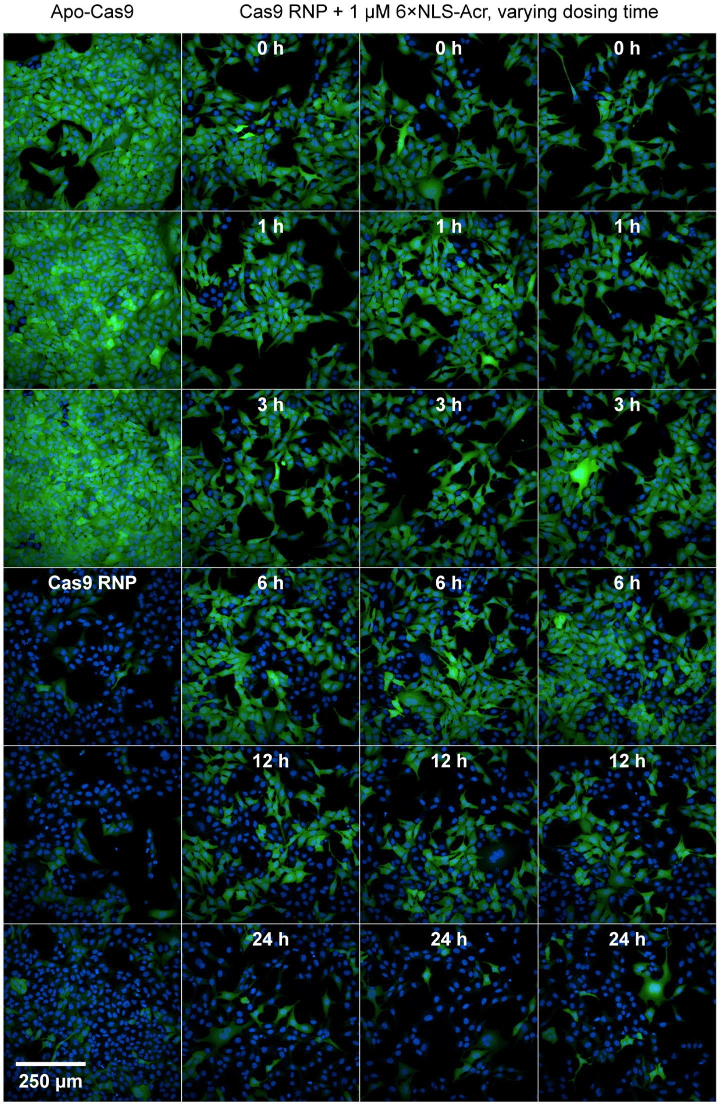


**Figure S9.** Representative fluorescence images from the *GFP*-disruption assay in Figure S8. SpCas9-NLS RNP (20 pmol) targeting *GFP* was delivered via nucleofection into U2OS-EGFP.PEST cells. The cells were seeded, incubated at 37 °C, and treated with 6×NLS-Acr (1 μM), varying the dosing time from 0 to 24 h. After 48 h, live cells were stained with Hoechst 33342 (blue, nucleus) and imaged using high-throughput confocal microscopy. Nucleofection of Cas9 RNP (white bar) serves as the control.

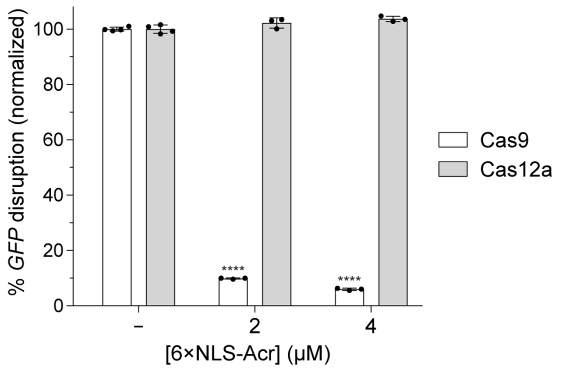


**Figure S10.** Effect of 6×NLS-Acr delivery on Cas protein activity in human cells. SpCas9-NLS (20 pmol) or 2×NLS-LbCas12a (20 pmol) RNP targeting *GFP* was delivered via nucleofection into U2OS-EGFP.PEST cells. The cells were seeded, incubated at 37 °C, and treated with 6×NLS-Acr (2–4 μM) at 0 h. After 48 h, live cells were stained with Hoechst 33342 and imaged using high-throughput confocal microscopy. Controls include nucleofection of Cas9 RNP (−, white bar) and nucleofection of Cas12a RNP (−, gray bar). The values were normalized to Cas9 RNP (white bars) or Cas12a RNP (gray bars) and are the mean ± SD of three independent replicates. The significance of 6×NLS-Acr delivery was determined with an unpaired, two-tailed *t*-test versus Cas9 RNP (white bars) or Cas12a RNP (gray bars), where *, **, ***, and **** refer to $P \leq 0.05$, $P \leq 0.01$, $P \leq 0.001$, and $P \leq 0.0001$, respectively.

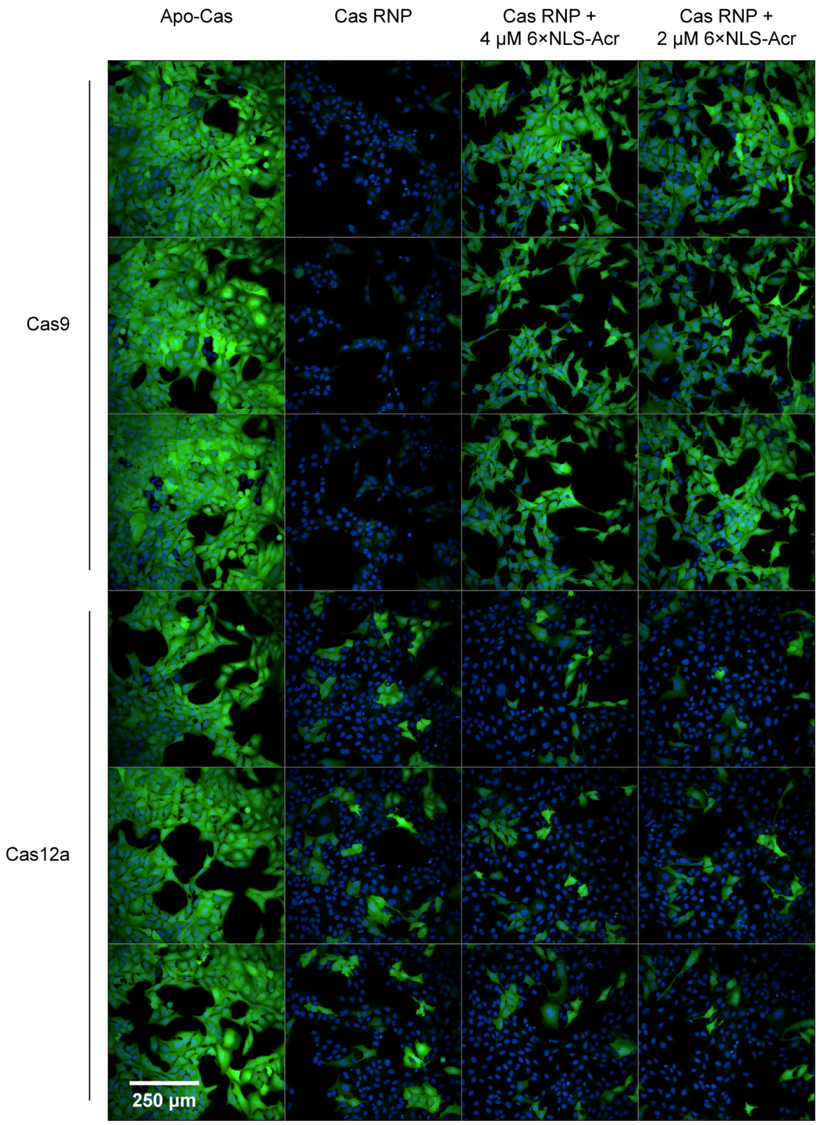


**Figure S11.** Representative fluorescence images from the *GFP*-disruption assay in Figure S10. SpCas9-NLS (20 pmol) or 2×NLS-LbCas12a (20 pmol) RNP targeting *GFP* was delivered via nucleofection into U2OS-EGFP.PEST cells. The cells were seeded, incubated at 37 °C, and treated with 6×NLS-Acr (2–4 μM) at 0 h. After 48 h, live cells were stained with Hoechst 33342 (blue, nucleus) and imaged using high-throughput confocal microscopy. Controls include nucleofection of Apo-Cas proteins and nucleofection of Cas RNPs.

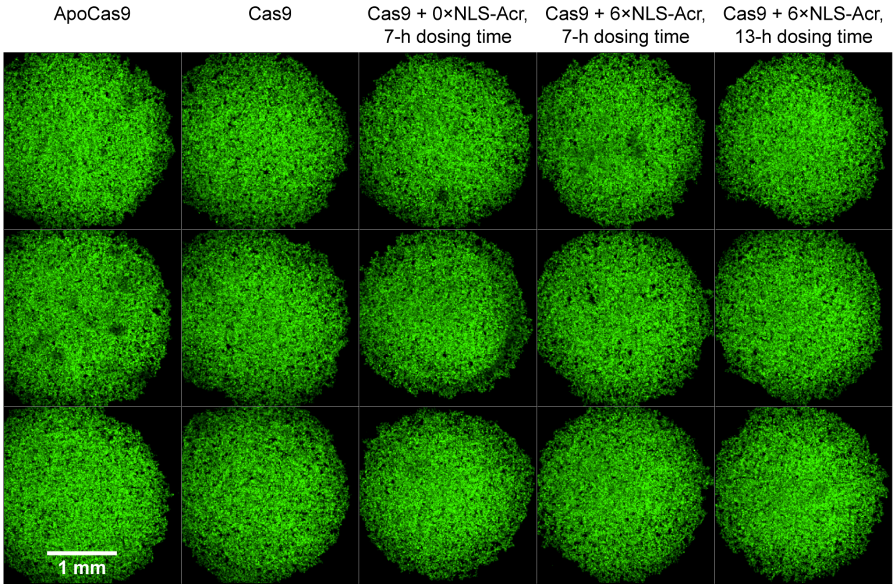


**Figure S12.** Fluorescence imaging shows loosely aggregated spheroids forming 7 h after cell seeding. SpCas9-NLS RNP (20 pmol) targeting *GFP* was delivered via nucleofection into U2OS-EGFP.PEST cells. The cells were seeded in a spheroid plate and incubated at 37 °C. After 7 h, live cells were imaged using high-throughput confocal microscopy. Controls include nucleofection of Apo-Cas9 and nucleofection of Cas9 RNP. Each column is labeled with the Acr that was eventually added to the 3D cell culture. However, these images were captured immediately before treating cells with Acrs at 7 h to show the appearance of the 3D cell cultures at the 7-h dosing time. The images represent maximum intensity projections from a Z-stack. The average diameter of the loosely aggregated spheroids is 2.53 ± 0.08 mm. Although *GFP* disruption has started by 7 h, the loosely aggregated spheroids appear equally fluorescent because GFP protein has not yet degraded.

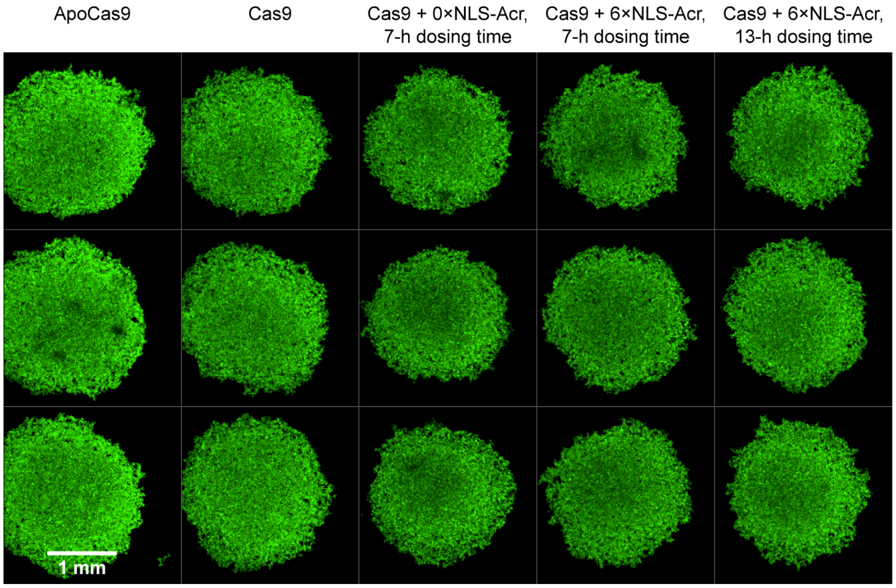


**Figure S13.** Fluorescence imaging shows loosely aggregated spheroids forming 13 h after cell seeding. SpCas9-NLS RNP (20 pmol) targeting *GFP* was delivered via nucleofection into U2OS-EGFP.PEST cells. The cells were seeded in a spheroid plate and incubated at 37 °C. After 13 h, live cells were imaged using high-throughput confocal microscopy. Controls include nucleofection of Apo-Cas9, nucleofection of Cas9 RNP, and nucleofection of Cas9 RNP followed by treatment with 0×NLS-Acr (2.5 μM) at 7 h. Each column is labeled with the Acr that was eventually added to the 3D cell culture. However, these images were captured immediately before treating cells with 6×NLS-Acr at 13 h to show the appearance of the 3D cell cultures at the 13-h dosing time. The images represent maximum intensity projections from a Z-stack. The average diameter of the loosely aggregated spheroids is 2.10 ± 0.08 mm. Although *GFP* disruption has started by 13 h, the loosely aggregated spheroids appear equally fluorescent because GFP protein has not yet degraded. These are the same cells imaged in Figure S12 but at a later time point (13 h).

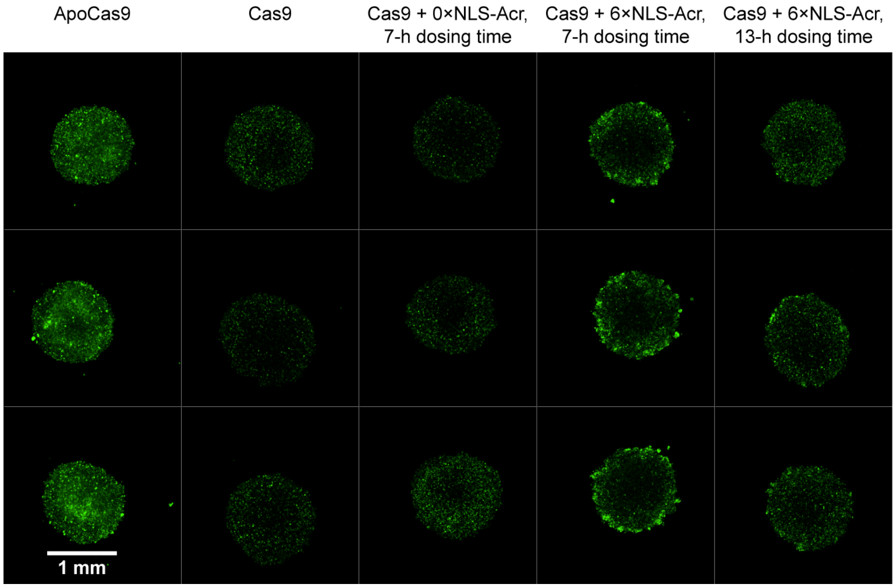


**Figure S14.** Representative fluorescence images from the *GFP*-disruption assay in Figure S16. SpCas9-NLS RNP (20 pmol) targeting *GFP* was delivered via nucleofection into U2OS-EGFP.PEST cells. The cells were seeded in a spheroid plate, incubated at 37 °C, and treated with 6×NLS-Acr (2.5 μM) at 7 and 13 h. After 48 h, live cells were imaged using high-throughput confocal microscopy. Controls include nucleofection of Apo-Cas9, nucleofection of Cas9 RNP, and nucleofection of Cas9 RNP followed by treatment with 0×NLS-Acr (2.5 μM) at 7 h. The images represent maximum intensity projections from a Z-stack. The approximate diameter of these well-defined spheroids is 1.25 ± 0.05 mm. These are the same cells imaged in Figures S12 and S13, but at a later time point (48 h).

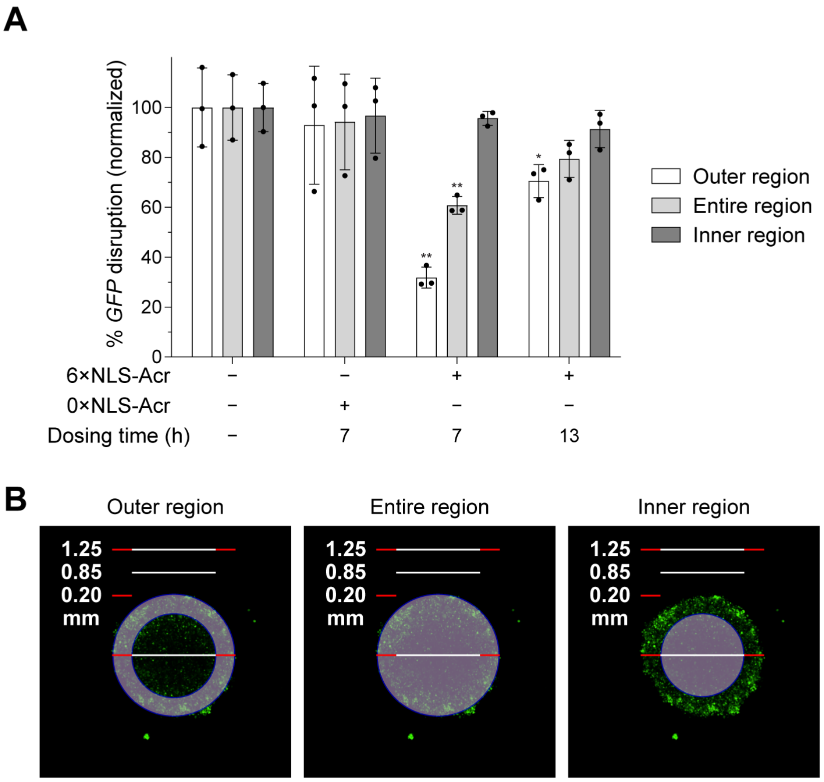


**Figure S15.** Effect of 6×NLS-Acr delivery on Cas9 activity in the different layers of the spheroid. (A) SpCas9-NLS RNP (20 pmol) targeting *GFP* was delivered via nucleofection into U2OS-EGFP.PEST cells. The cells were seeded in a spheroid plate, incubated at 37 °C, and treated with 6×NLS-Acr (2.5 µM) at 7 and 13 h. After 48 h, live cells were imaged using high-throughput confocal microscopy. Controls include nucleofection of Cas9 RNP (−, dosing time) and nucleofection of Cas9 RNP followed by treatment with 0×NLS-Acr (2.5 µM) at 7 h. White bars show *GFP* disruption in the outer spheroid region (the area that is 0.20 mm in depth). Light gray bars show *GFP* disruption in the entire spheroid region, which has an average diameter of 1.25 mm. Dark gray bars show *GFP* disruption in the inner spheroid region (the area that is 0.85 mm in diameter). The values were normalized to Cas9 RNP for each region and are the mean ± SD of three independent replicates. The significance of 6×NLS-Acr delivery was determined with an unpaired, two-tailed *t*-test versus Cas9 RNP for each region, where *, **, ***, and **** refer to $P \leq 0.05$, $P \leq 0.01$, $P \leq 0.001$, and $P \leq 0.0001$, respectively. (B) Visual representation of the spheroid regions defined to quantify *GFP* disruption in panel A. Each region (outer, entire, and inner region) is shaded in light purple. 6×NLS-Acr delivery at 7 and 13 h reduced *GFP* disruption in the outer spheroid region.

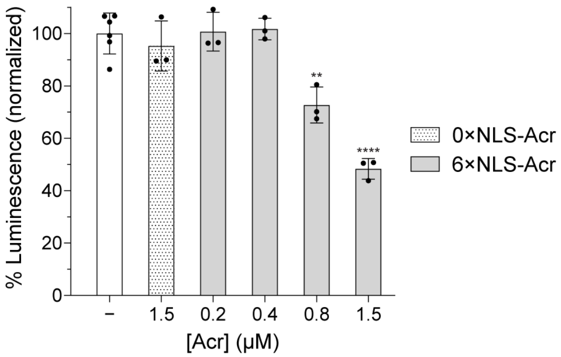


**Figure S16.** Dose-dependent inhibition of Cas9 by 6×NLS-Acr in the HiBiT-knock-in assay (HEK293T cells). SpCas9-NLS RNP (20 pmol) targeting *GAPDH* and the HiBiT ssODN (80 pmol) were co-delivered via nucleofection into HEK293T cells. The cells were seeded, incubated at 37 °C, and treated with 6×NLS-Acr (0.2–1.5 μM) at 0 h. After 72 h, the cells were lysed and combined with LgBiT and furimazine, and their luminescence was quantified. Controls include co-nucleofection of Cas9 RNP and ssODN (white bar) and co-nucleofection of Cas9 RNP and ssODN followed by treatment with 0×NLS-Acr (1.5 μM) at 0 h. The values were normalized to Cas9 RNP + ssODN and are the mean ± SD of three independent replicates. The significance of 6×NLS-Acr delivery was determined with an unpaired, two-tailed *t*-test versus Cas9 RNP + ssODN, where *, **, ***, and **** refer to $P \leq 0.05$, $P \leq 0.01$, $P \leq 0.001$, and $P \leq 0.0001$, respectively.

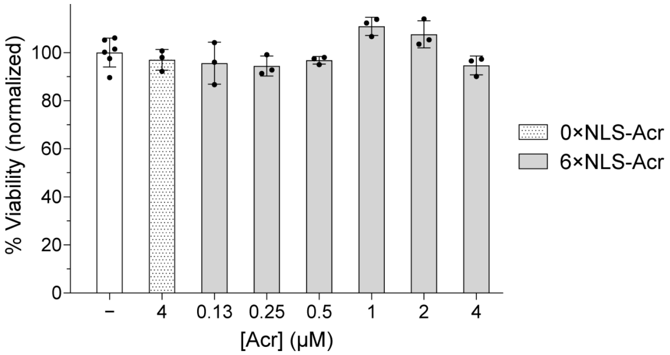


**Figure S17.** Cell viability after 6×NLS-Acr delivery in the HiBiT-knock-in assay (HUES 8 cells). SpCas9-NLS RNP (20 pmol) targeting *GAPDH* and the HiBiT ssODN (80 pmol) were co-delivered via nucleofection into HUES 8 cells. The cells were seeded, incubated at 37 °C, and treated with 6×NLS-Acr (0.13–4 µM) at 0 h. After 24 h, the cells were incubated with the PrestoBlue HS reagent. Cell viability was determined using fluorescence spectroscopy (560 nm excitation/590 nm emission). Controls include co-nucleofection of Cas9 RNP and ssODN (white bar) and co-nucleofection of Cas9 RNP and ssODN followed by treatment with 0×NLS-Acr (4 µM) at 0 h. The values were normalized to Cas9 RNP with ssODN and are the mean ± SD of three independent replicates. The significance of 6×NLS-Acr delivery was determined with an unpaired, two-tailed *t*-test versus Cas9 RNP with ssODN. No statistically significant decreases in viability were detected.

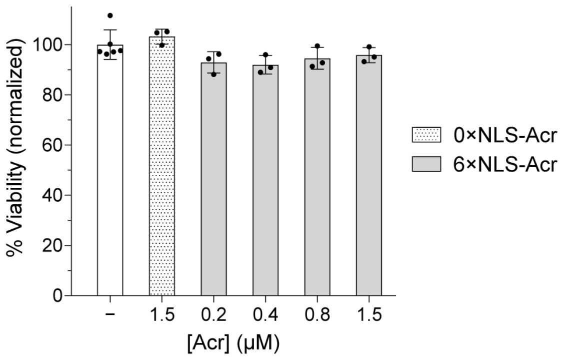


**Figure S18.** Cell viability after 6×NLS-Acr delivery in the HiBiT-knock-in assay (HEK293T cells). SpCas9-NLS RNP (20 pmol) targeting *GAPDH* and the HiBiT ssODN (80 pmol) were co-delivered via nucleofection into HEK293T cells. The cells were seeded, incubated at 37 °C, and treated with 6×NLS-Acr (0.2–1.5 µM) at 0 h. After 72 h, the cells were incubated with the PrestoBlue HS reagent. Cell viability was determined using fluorescence spectroscopy (560 nm excitation/590 nm emission). Controls include co-nucleofection of Cas9 RNP and ssODN (white bar) and co-nucleofection of Cas9 RNP and ssODN followed by treatment with 0×NLS-Acr (1.5 µM) at 0 h. The values were normalized to Cas9 RNP with ssODN and are the mean ± SD of three independent replicates. The significance of 6×NLS-Acr delivery was determined with an unpaired, two-tailed *t*-test versus Cas9 RNP with ssODN. No statistically significant decreases in viability were detected.

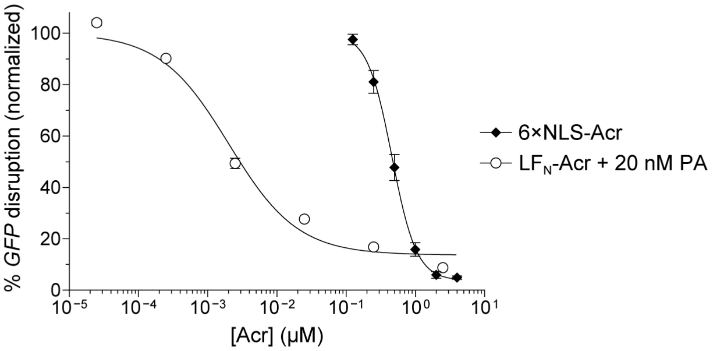


**Figure S19.** Dose-dependent inhibition of Cas9 by 6×NLS-Acr and $LF_N$-Acr in the *GFP*-disruption assay. SpCas9-NLS RNP (20 pmol) targeting *GFP* was delivered via nucleofection into U2OS-EGFP.PEST cells. The cells were seeded, incubated at 37 °C, and treated with 6×NLS-Acr (0.13, 0.25, 0.5, 1, 2, and 4 μM) or $LF_N$-Acr ($2.5 \times 10^{-5}$, $2.5 \times 10^{-4}$, $2.5 \times 10^{-3}$, 0.025, 0.25, and 2.5 μM) plus PA (20 nM) at 0 h. After 48 h, live cells were stained with Hoechst 33342 and imaged using high-throughput confocal microscopy. The values were normalized to Cas9 RNP (100%, not shown) and are the mean ± SD of three independent replicates. Some error bars are too small to visualize.

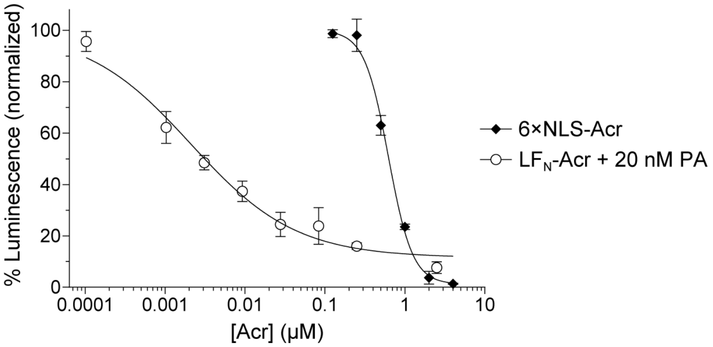


**Figure S20.** Dose-dependent inhibition of Cas9 by 6×NLS-Acr and $LF_N$-Acr in the HiBiT-knock-in assay. SpCas9-NLS RNP (20 pmol) targeting *GAPDH* and the HiBiT ssODN (80 pmol) were co-delivered via nucleofection into HUES 8 cells. The cells were seeded, incubated at 37 °C, and treated with 6×NLS-Acr (0.13, 0.25, 0.5, 1, 2, and 4 µM) or $LF_N$-Acr (0.0001, 0.001, 0.003, 0.009, 0.028, 0.083, 0.25, and 2.5 µM) plus PA (20 nM) at 0 h. After 24 h, the cells were lysed and combined with LgBiT and furimazine, and their luminescence was quantified. The values were normalized to Cas9 RNP with ssODN (100%, not shown) and are the mean ± SD of three independent replicates. Some error bars are too small to visualize.

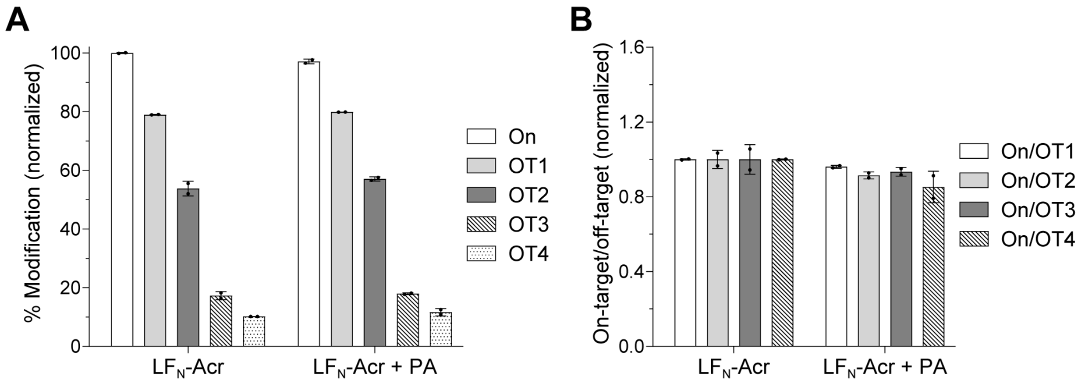


**Figure S21.** Effect of LF$_N$-Acr delivery on genome-editing activity and specificity. (A) LF$_N$-Acr delivery is ineffective after Cas9 lipofection. Plasmids encoding 2×NLS-SpCas9 (750 ng) and *EMX1*-targeting gRNA (250 ng) were delivered via lipofection into HEK293T cells. The cells were incubated at 37 °C and treated with LF$_N$-Acr (250 nM) ± PA (20 nM) at 1 h. After 72 h, genomic DNA was extracted, sequenced with NGS, and analyzed using CRISPResso2 to determine the % modification at the on-target (On) and off-target (OT1, OT2, OT3, and OT4) sites. The values were normalized to Cas9·gRNA with LF$_N$-Acr (On) and are the mean ± SD of two independent replicates. The significance of PA-mediated LF$_N$-Acr delivery was determined using an unpaired, two-tailed *t*-test versus Cas9·gRNA with LF$_N$-Acr (separately for On, OT1, OT2, OT3, and OT4), where *, **, ***, and **** refer to $P \leq 0.05$, $P \leq 0.01$, $P \leq 0.001$, and $P \leq 0.0001$, respectively. The differences are not statistically significant. (B) Because of its ineffective delivery after Cas9 lipofection, LF$_N$-Acr combined with PA does not increase Cas9 specificity. Specificity was calculated as the ratio of on-target to off-target % modification using data from panel A, normalized to the on-target to off-target % modification ratio of Cas9·gRNA with LF$_N$-Acr (separately for On/OT1, On/OT2, On/OT3, and On/OT4). The significance of the specificity was determined using an unpaired, two-tailed *t*-test compared to the specificity of Cas9·gRNA with LF$_N$-Acr (separately for On/OT1, On/OT2, On/OT3, and On/OT4), where *, **, ***, and **** refer to $P \leq 0.05$, $P \leq 0.01$, $P \leq 0.001$, and $P \leq 0.0001$, respectively. The differences are not statistically significant.

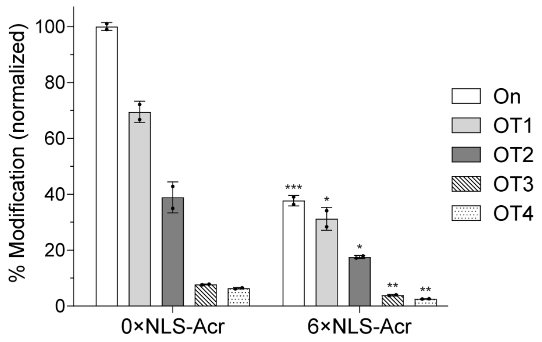


**Figure S22.** 6×NLS-Acr delivery regulates the activity of a less active Cas9 variant. Plasmids encoding SpCas9-NLS (750 ng) and *EMX1*-targeting gRNA (250 ng) were delivered via lipofection into HEK293T cells. The cells were incubated at 37 °C and treated with 0×NLS-Acr (4 μM, control) or 6×NLS-Acr (4 μM) at 1 h. After 72 h, genomic DNA was extracted, sequenced with NGS, and analyzed using CRISPResso2 to determine the % modification at the on-target (On) and off-target (OT1, OT2, OT3, and OT4) sites. The values were normalized to Cas9·gRNA with 0×NLS-Acr (On) and are the mean ± SD of two independent replicates. The significance of 6×NLS-Acr delivery was determined using an unpaired, two-tailed *t*-test versus Cas9·gRNA with 0×NLS-Acr (separately for On, OT1, OT2, OT3, and OT4), where *, **, ***, and **** refer to $P \leq 0.05$, $P \leq 0.01$, $P \leq 0.001$, and $P \leq 0.0001$, respectively.

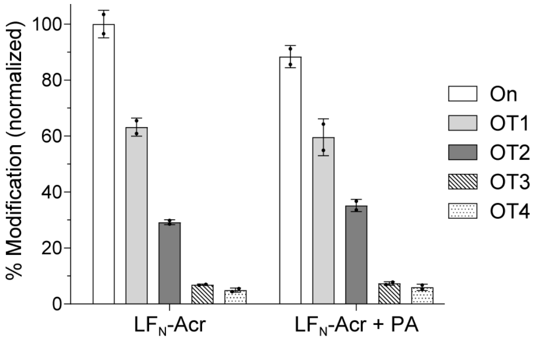


**Figure S23.** LF$_N$-Acr delivery is ineffective after lipofection of a less active Cas9 variant. Plasmids encoding SpCas9-NLS (750 ng) and *EMX1*-targeting gRNA (250 ng) were delivered via lipofection into HEK293T cells. The cells were incubated at 37 °C and treated with LF$_N$-Acr (250 nM) ± PA (20 nM) at 1 h. After 72 h, genomic DNA was extracted, sequenced with NGS, and analyzed using CRISPResso2 to determine the % modification at the on-target (On) and off-target (OT1, OT2, OT3, and OT4) sites. The values were normalized to Cas9·gRNA with LF$_N$-Acr (On) and are the mean ± SD of two independent replicates. The significance of PA-mediated LF$_N$-Acr delivery was determined using an unpaired, two-tailed *t*-test versus Cas9·gRNA with LF$_N$-Acr (separately for On, OT1, OT2, OT3, and OT4), where *, **, ***, and **** refer to $P \leq 0.05$, $P \leq 0.01$, $P \leq 0.001$, and $P \leq 0.0001$, respectively. The differences are not statistically significant.

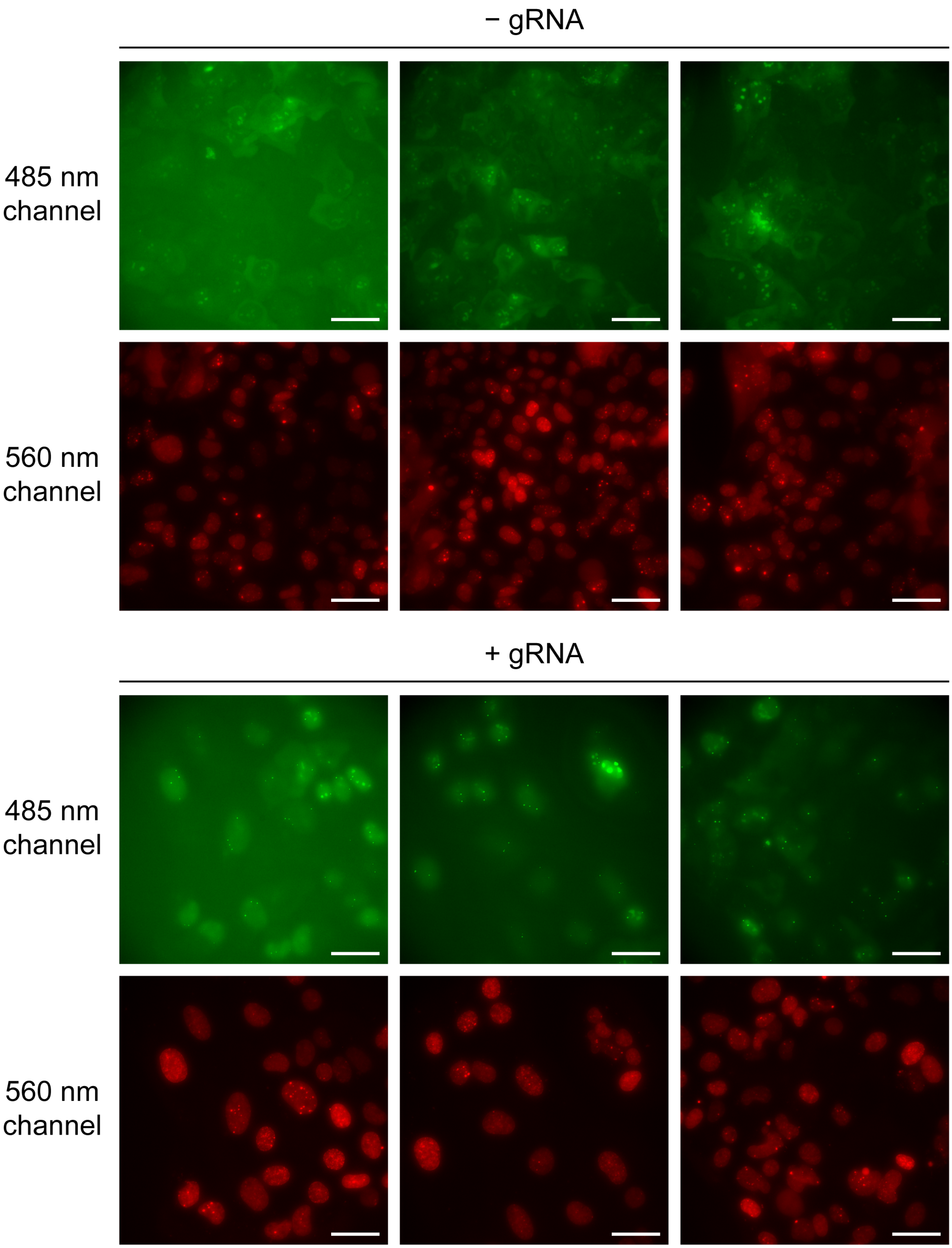


**Figure S24A.** (The figure is continued on the next page.)

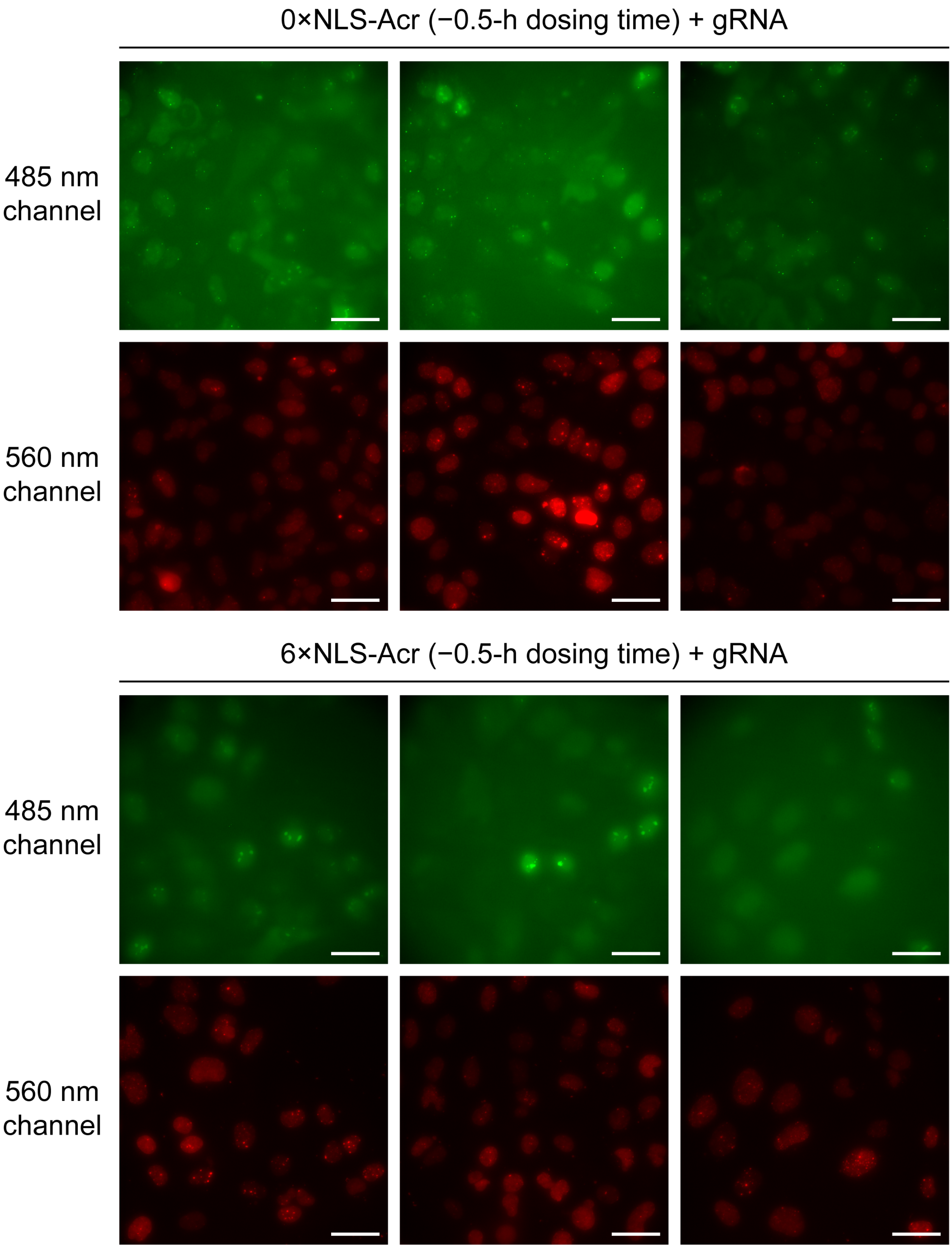


**Figure S24B.** (The figure is continued on the next page.)

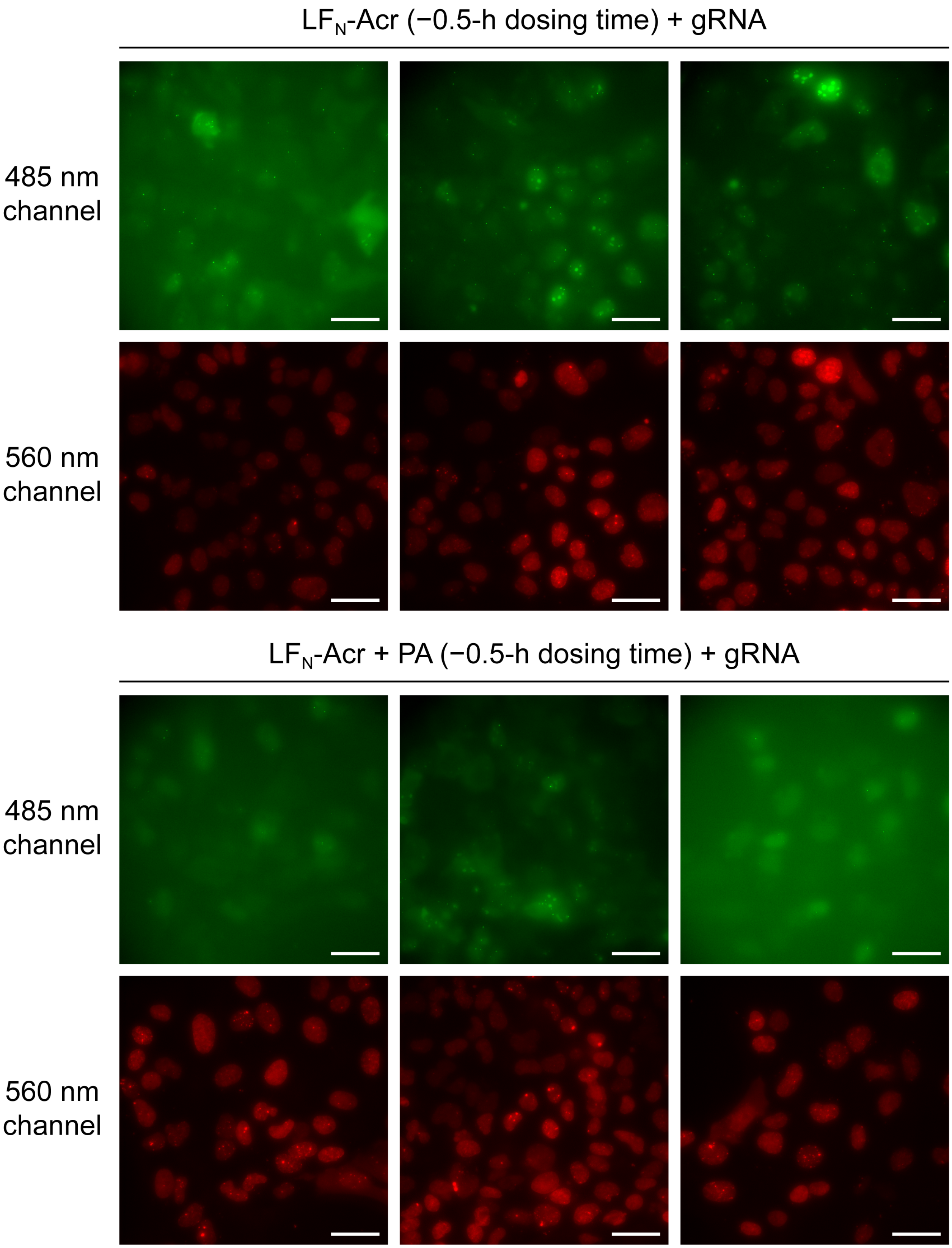


**Figure S24C.** (The figure is continued on the next page.)

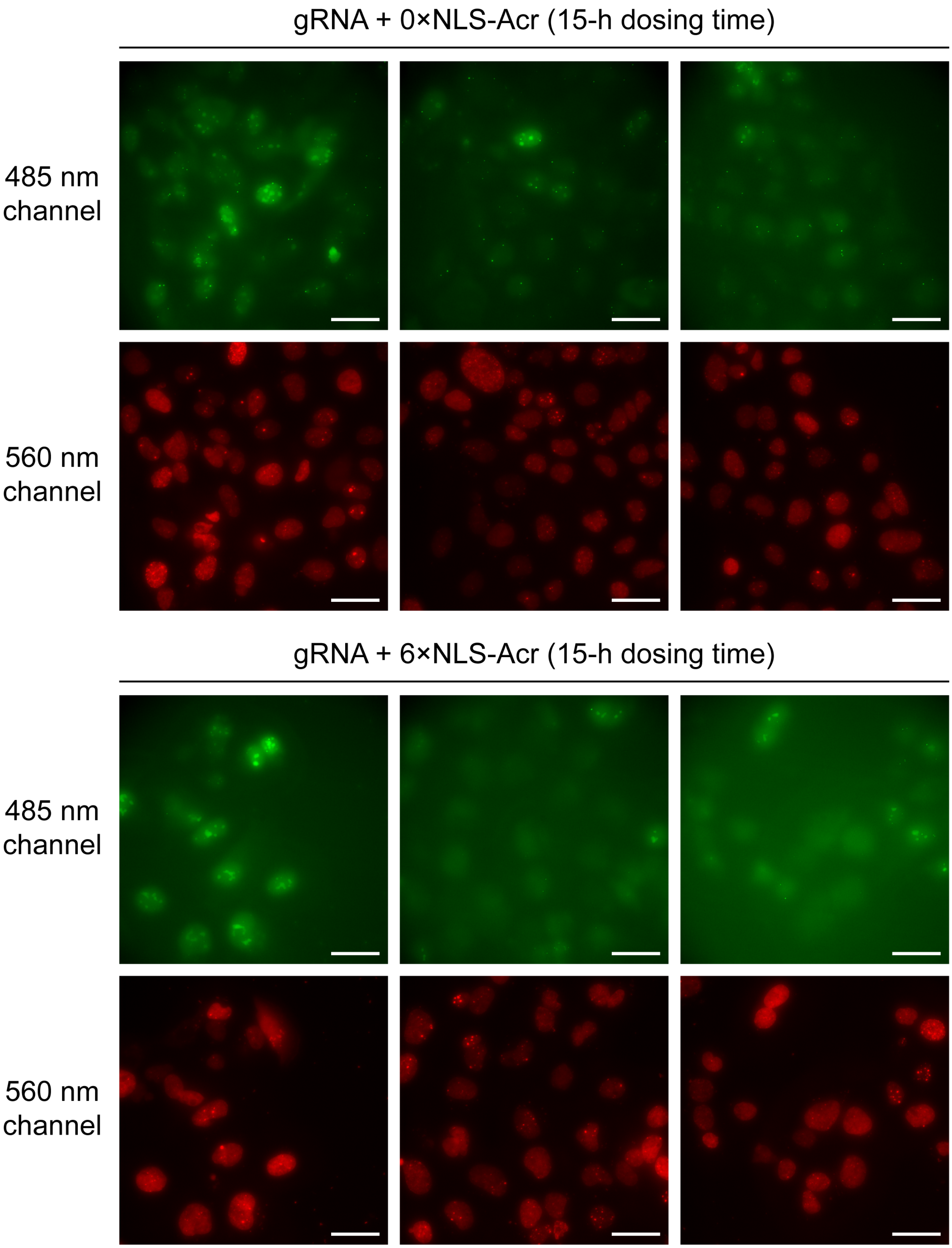


**Figure S24D.** (The figure is continued on the next page.)

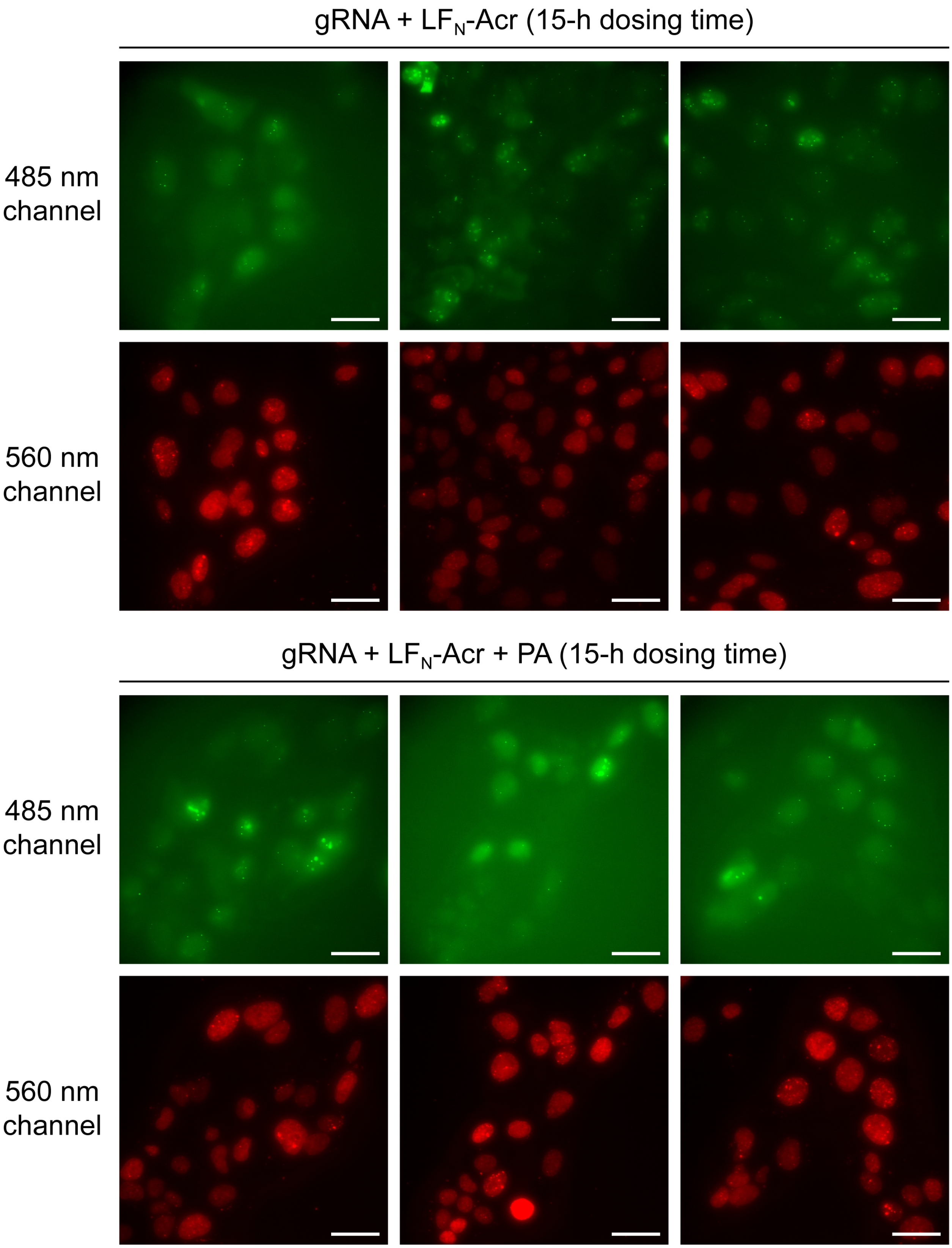


**Figure S24E.** (The figure is continued on the next page.)

**Figure S24.** (A–E) Direct visualization of the inhibition of Cas9–GFP by 6×NLS-Acr and $LF_N$-Acr. Ch3Rep gRNA was delivered via lipofection into a U2OS cell line stably expressing SpCas9–GFP and 53BP1–mCherry. Cells were treated with 6×NLS-Acr (2.5 μM) or $LF_N$-Acr (2.5 μM) plus PA (20 nM) either 0.5 h before (−0.5 h, panels B and C) or 15 h after (15 h, panels D and E) gRNA transfection. Controls include cells transfected with or without gRNA (panel A) and cells treated with 0×NLS-Acr (2.5 μM) or $LF_N$-Acr (2.5 μM) at −0.5 h (panels B and C) or 15 h (panels D and E). Live cells were imaged using widefield fluorescence microscopy (60× objective) at 15 h (panels A, B, and C) and 20 h (panels D and E) to detect fluorescence from Cas9–GFP (green; 485 nm excitation) and 53BP1–mCherry (red nuclear marker; 560 nm excitation). Three representative images are included for each condition. Scale bars, 40 μm.

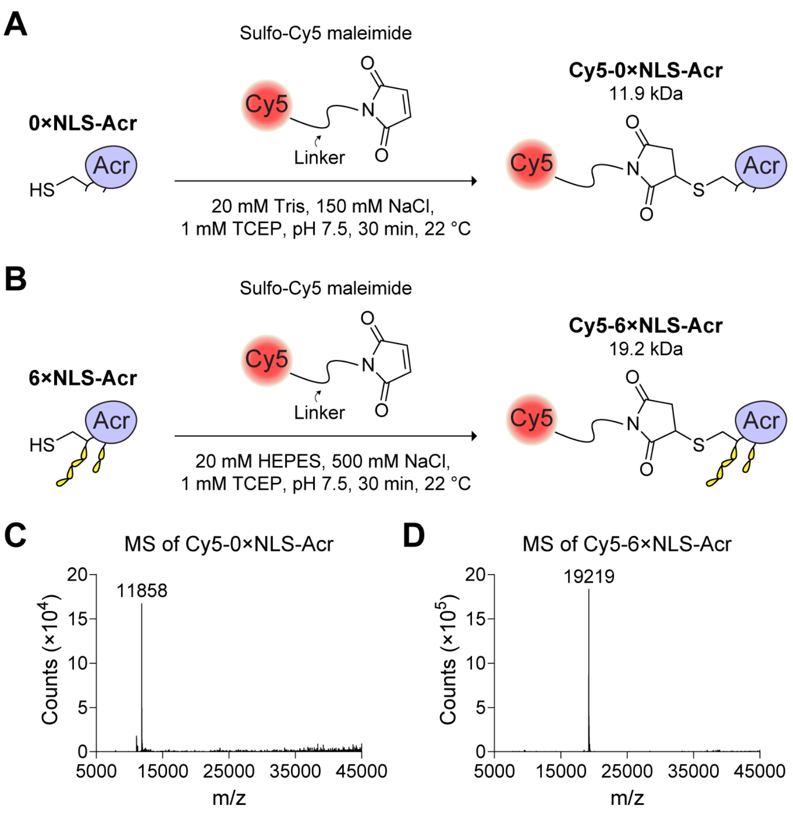


**Figure S25.** (A) Synthesis of Cy5-0×NLS-Acr. 0×NLS-Acr was reacted with sulfo-Cy5 maleimide to generate Cy5-0×NLS-Acr. (B) Synthesis of Cy5-6×NLS-Acr. 6×NLS-Acr was reacted with sulfo-Cy5 maleimide to generate Cy5-6×NLS-Acr. (C) Q-TOF LC–MS of Cy5-0×NLS-Acr. LC–MS confirmed that 0×NLS-Acr (11,093 Da) was almost entirely converted to Cy5-0×NLS-Acr (11,858 Da). A low-abundance peak with an m/z of 11,095 was also detected in the mass spectrum. This m/z is close to the mass of 0×NLS-Acr (11,093 Da), suggesting a small amount of unreacted 0×NLS-Acr is present. (D) Q-TOF LC–MS of Cy5-6×NLS-Acr. LC–MS confirmed that 6×NLS-Acr (18,454 Da) was fully converted to Cy5-6×NLS-Acr (19,219 Da).

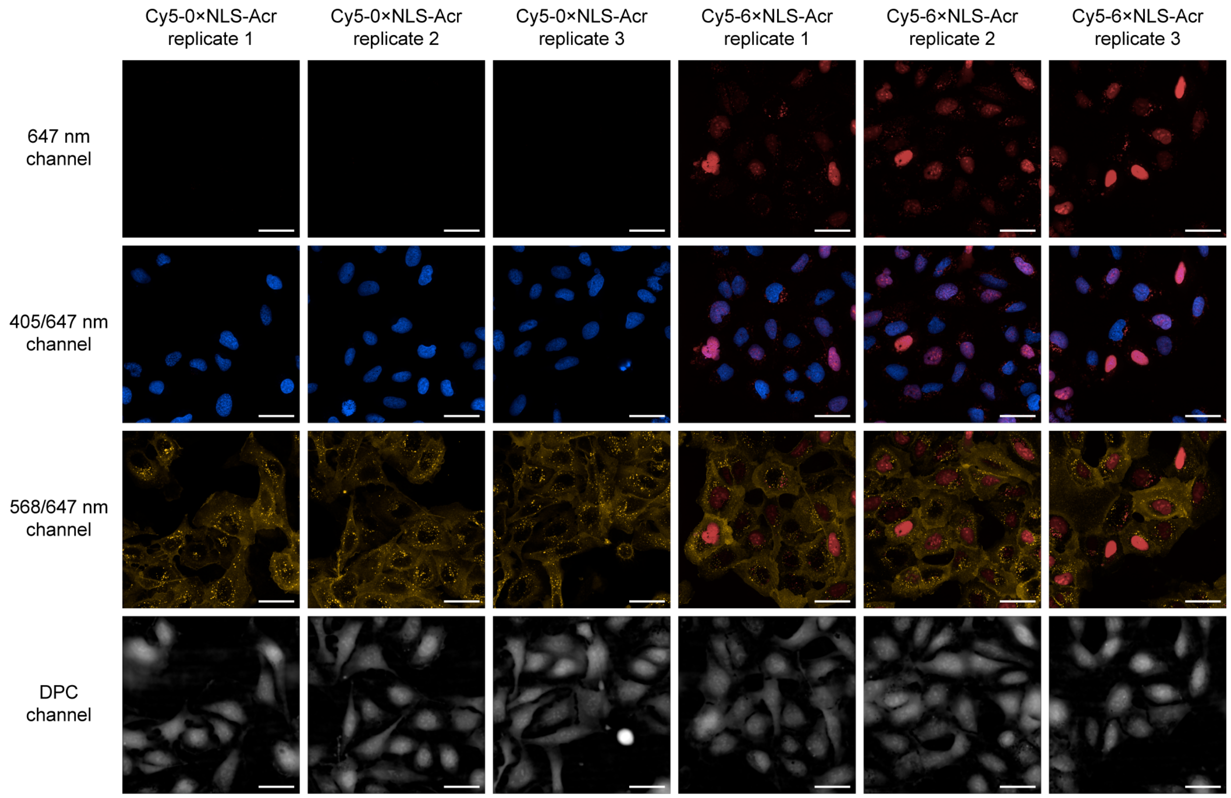


**Figure S26.** Visualizing the delivery of fluorescently labeled Acrs at 37 °C. U2OS cells were seeded and incubated overnight at 37 °C. Cells were then treated with Cy5-0×NLS-Acr (1.5 µM, control) or Cy5-6×NLS-Acr (1.5 µM) and incubated for 20 min at 37 °C. Cells were washed and stained with Hoechst 33342 and WGA555 in FluoroBrite DMEM, then imaged using confocal microscopy (60× objective) to detect fluorescence from Hoechst 33342 (blue nuclear marker; 405 nm excitation), WGA555 (yellow cytosolic marker; 568 nm excitation), and Cy5 (red; 647 nm excitation). Digital phase contrast (DPC) microscopy was also performed. Scale bars, 40 µm.

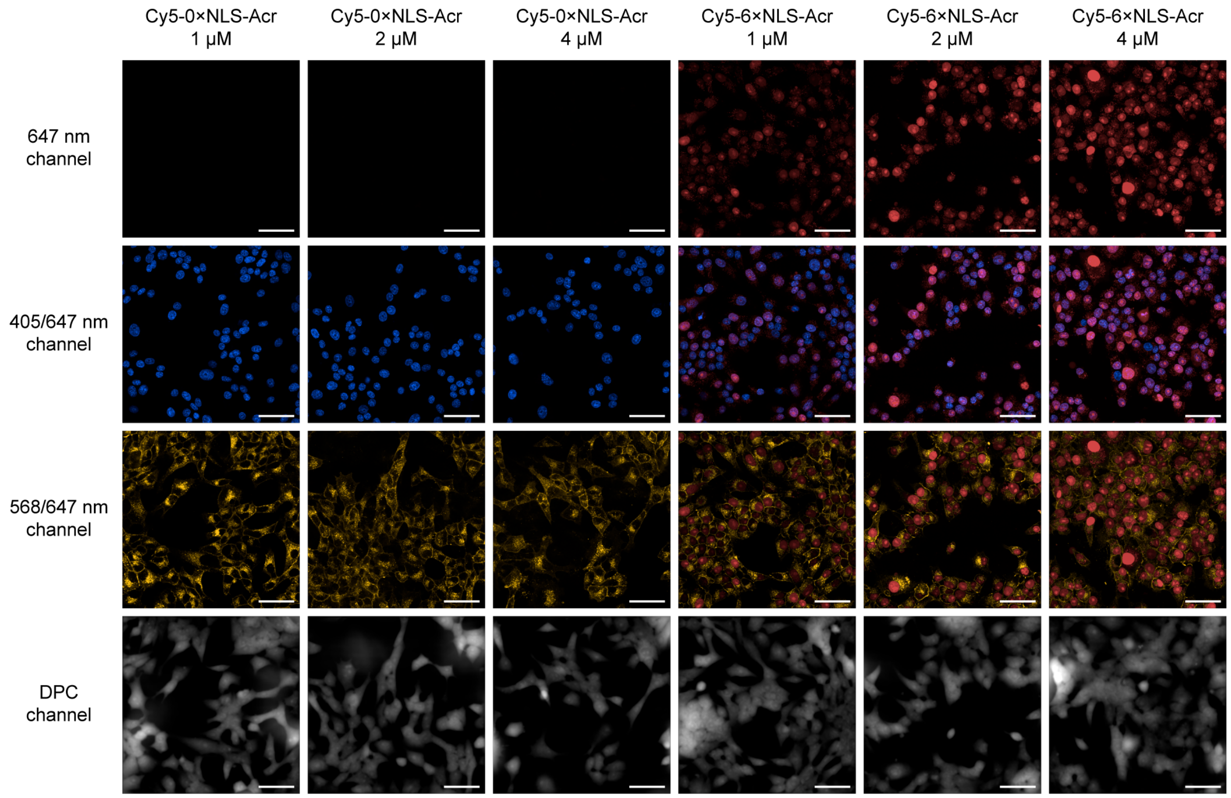


**Figure S27.** Dose-dependent delivery of fluorescently labeled Acrs. INS-1E rat β-cells were seeded and incubated overnight at 37 °C. Cells were then treated with Cy5-0×NLS-Acr (1–4 μM, control) or Cy5-6×NLS-Acr (1–4 μM) and incubated for 2 h at 37 °C. Cells were washed and stained with Hoechst 33342 and WGA555 in FluoroBrite DMEM, then imaged using confocal microscopy (60× objective) to detect fluorescence from Hoechst 33342 (blue nuclear marker; 405 nm excitation), WGA555 (yellow cytosolic marker; 568 nm excitation), and Cy5 (red; 647 nm excitation). Digital phase contrast (DPC) microscopy was also performed. Scale bars, 40 μm.

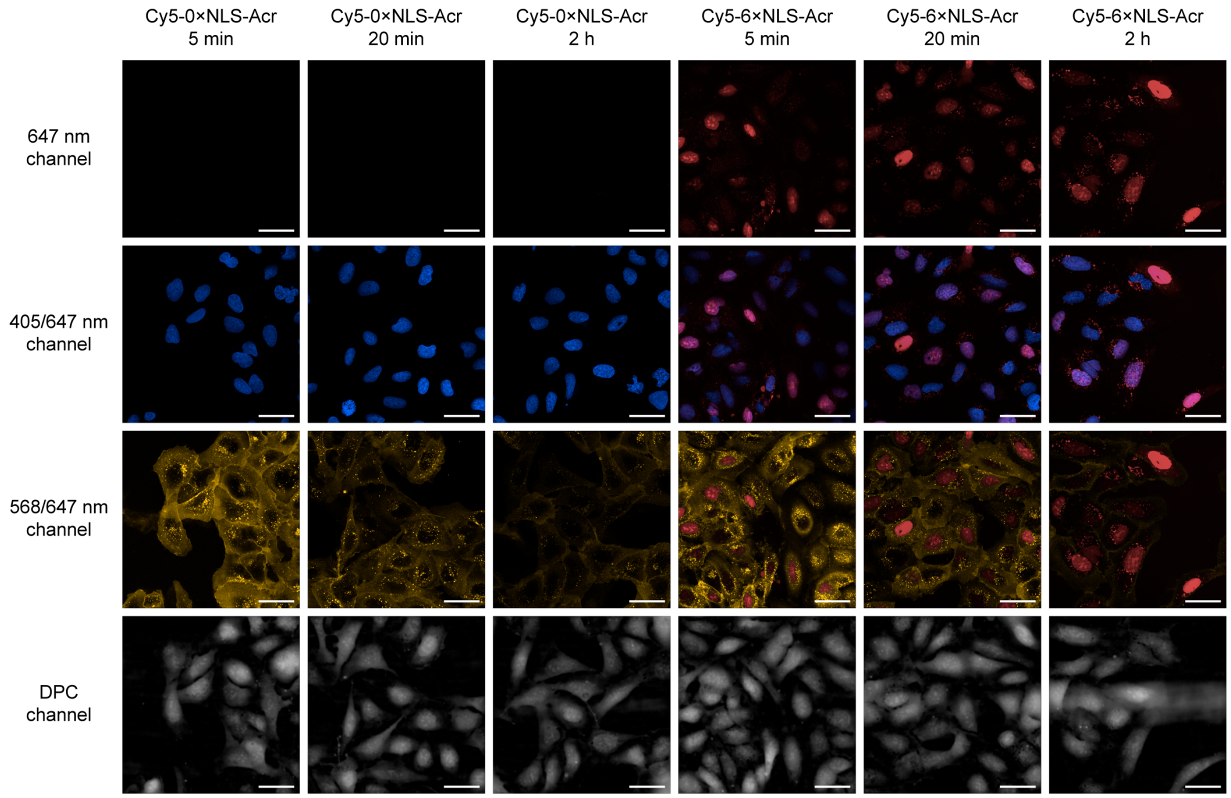


**Figure S28.** Time-dependent delivery of fluorescently labeled Acrs. U2OS cells were seeded and incubated overnight at 37 °C. Cells were then treated with Cy5-0×NLS-Acr (1.5 µM, control) or Cy5-6×NLS-Acr (1.5 µM) and incubated for 5 min to 2 h at 37 °C. Cells were washed and stained with Hoechst 33342 and WGA555 in FluoroBrite DMEM, then imaged using confocal microscopy (60× objective) to detect fluorescence from Hoechst 33342 (blue nuclear marker; 405 nm excitation), WGA555 (yellow cytosolic marker; 568 nm excitation), and Cy5 (red; 647 nm excitation). Digital phase contrast (DPC) microscopy was also performed. Scale bars, 40 µm.

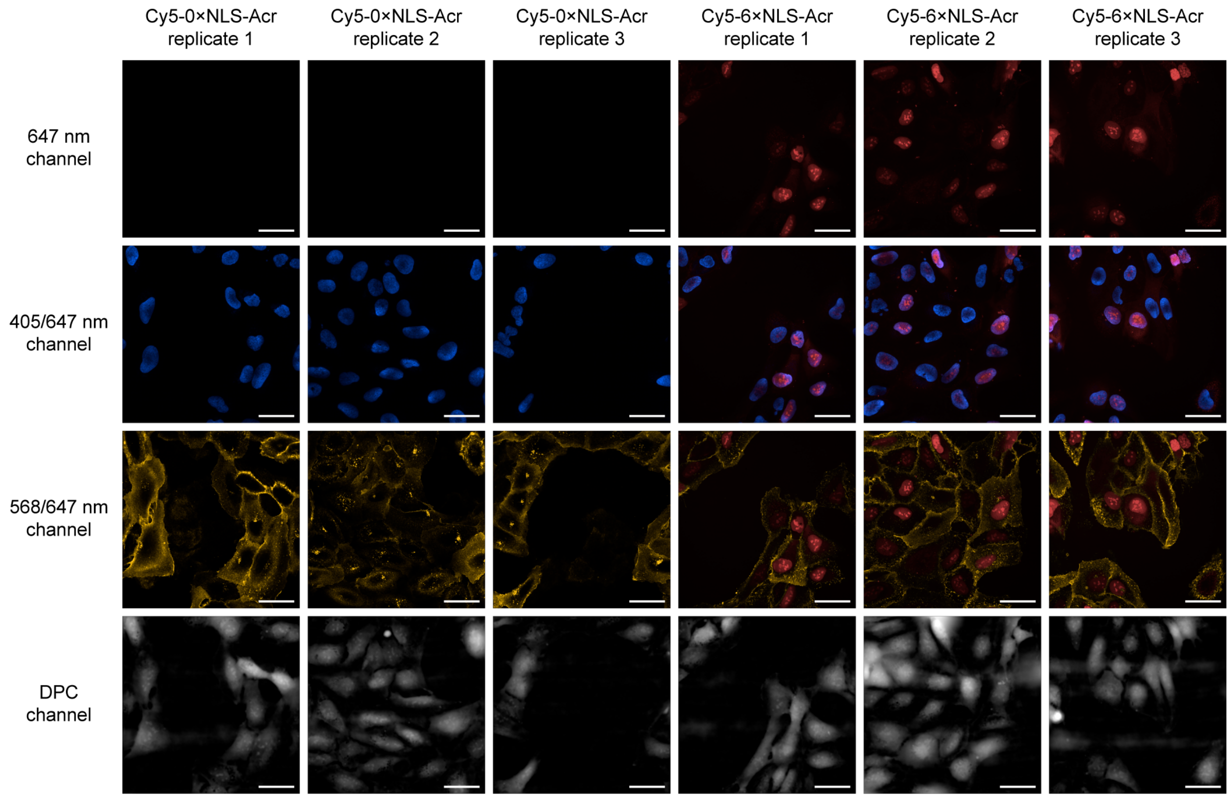


**Figure S29.** Visualizing the delivery of fluorescently labeled Acrs at 4 °C. U2OS cells were seeded and incubated overnight at 37 °C. Cells were then treated with Cy5-0×NLS-Acr (1.5 µM, control) or Cy5-6×NLS-Acr (1.5 µM) and incubated for 20 min at 4 °C. Cells were washed and stained with Hoechst 33342 and WGA555 in FluoroBrite DMEM, then imaged using confocal microscopy (60× objective) to detect fluorescence from Hoechst 33342 (blue nuclear marker; 405 nm excitation), WGA555 (yellow cytosolic marker; 568 nm excitation), and Cy5 (red; 647 nm excitation). Digital phase contrast (DPC) microscopy was also performed. Scale bars, 40 µm.

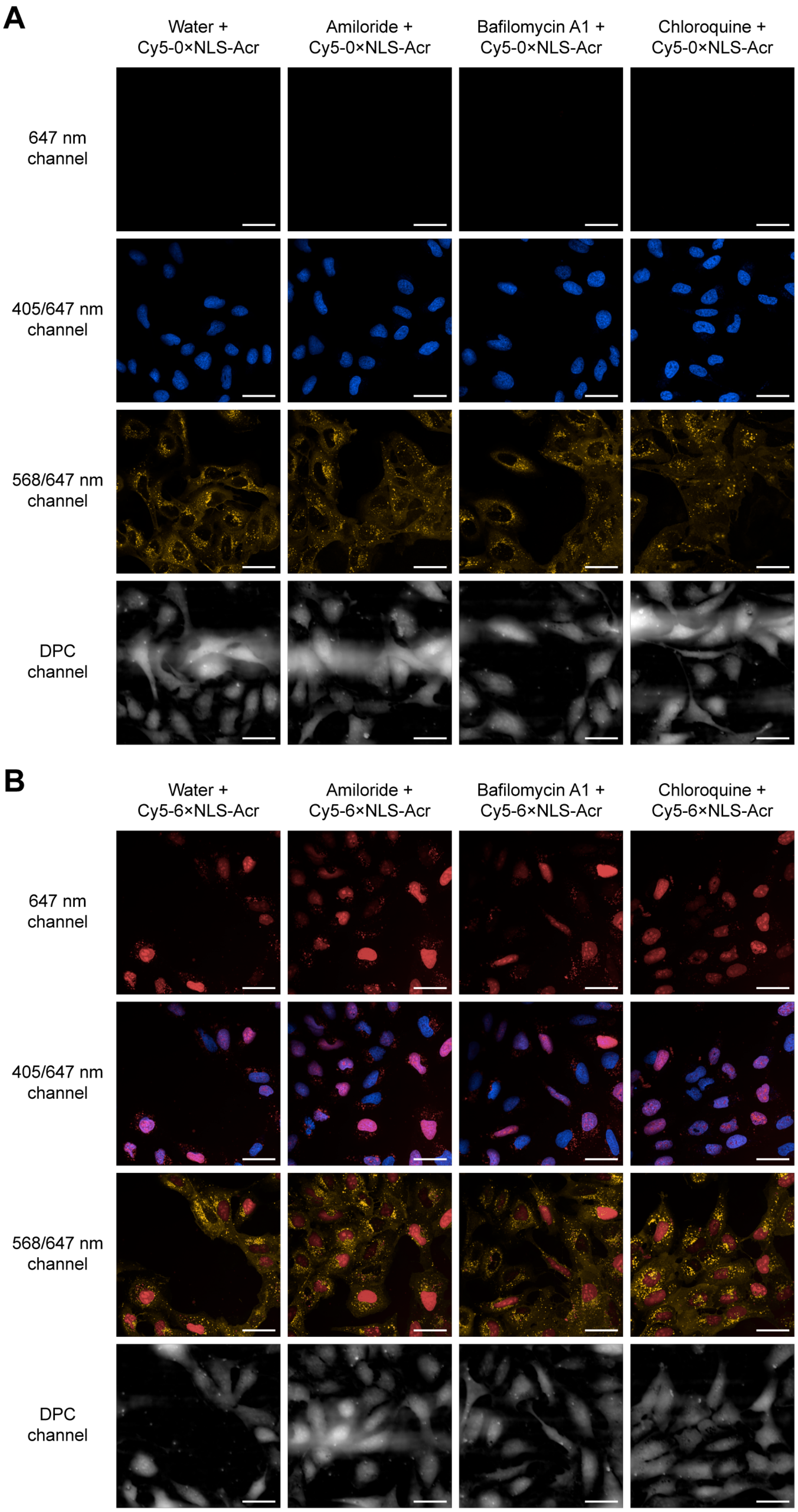


**Figure S30.** (The figure is continued on the next page.)

**Figure S30.** Effect of small-molecule inhibitors of endocytosis on the delivery of fluorescently labeled Acrs. U2OS cells were seeded and incubated overnight at 37 °C. Cells were first treated with water (2% v/v, control) or aqueous solutions (2% v/v) of amiloride hydrochloride (50 μM), bafilomycin A1 (25 nM), or chloroquine diphosphate (100 μM), and incubated for 3 h at 37 °C. Cells were then treated with Cy5-0×NLS-Acr (1.5 μM, panel A) or Cy5-6×NLS-Acr (1.5 μM, panel B) and incubated for 20 min at 37 °C. Cells were washed and stained with Hoechst 33342 and WGA555 in FluoroBrite DMEM, then imaged using confocal microscopy (60× objective) to detect fluorescence from Hoechst 33342 (blue nuclear marker; 405 nm excitation), WGA555 (yellow cytosolic marker; 568 nm excitation), and Cy5 (red; 647 nm excitation). Digital phase contrast (DPC) microscopy was also performed. Scale bars, 40 μm.

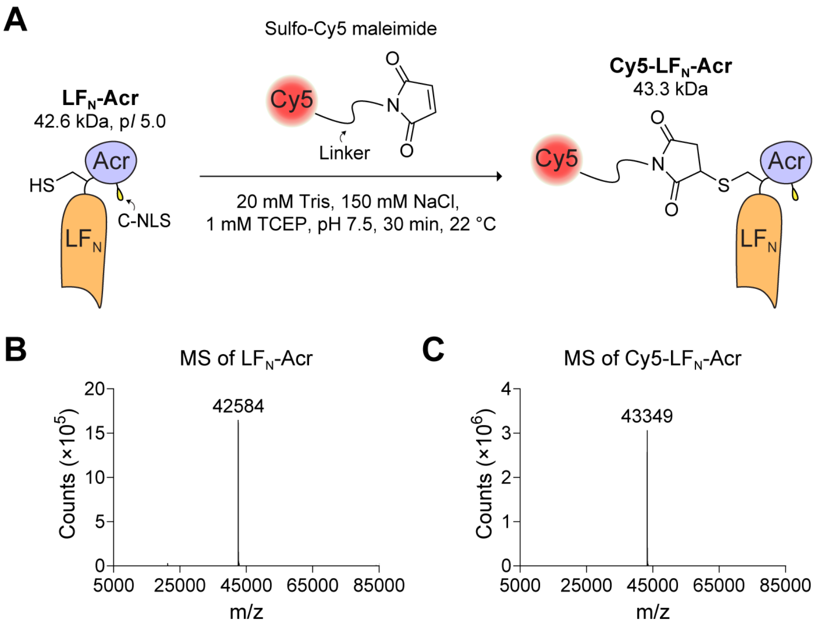


**Figure S31.** (A) Synthesis of Cy5-LF$_N$-Acr. LF$_N$-Acr was reacted with sulfo-Cy5 maleimide to generate Cy5-LF$_N$-Acr. (B) Q-TOF LC–MS of LF$_N$-Acr. (C) Q-TOF LC–MS of Cy5-LF$_N$-Acr. LC–MS confirmed that LF$_N$-Acr (42,584 Da) was fully converted to Cy5-LF$_N$-Acr (43,349 Da).

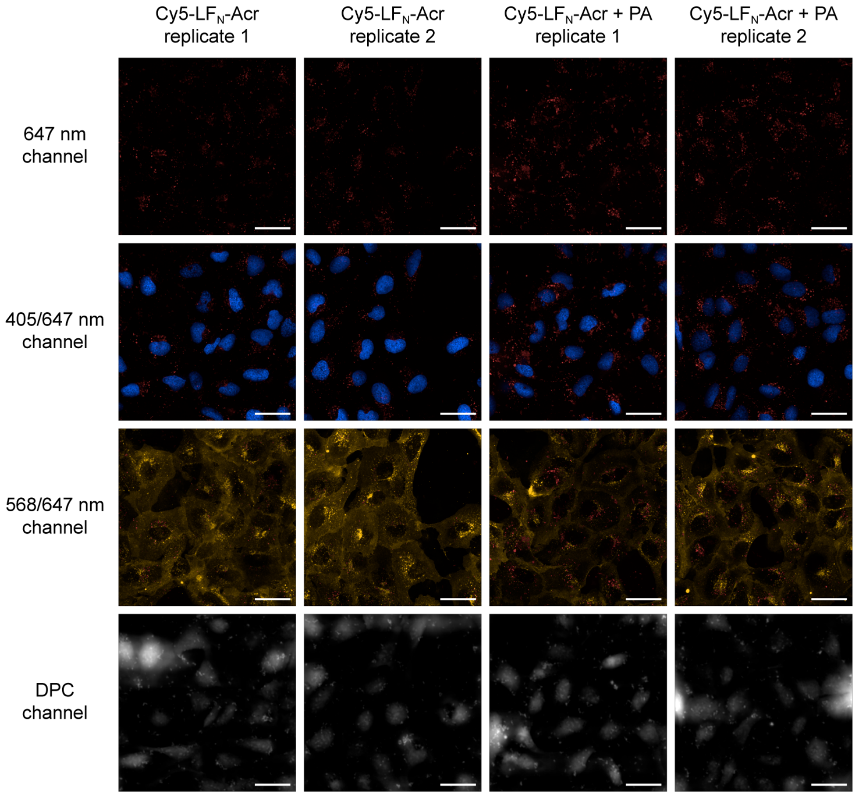


**Figure S32.** Visualizing the delivery of fluorescently labeled $LF_N$-Acr. U2OS cells were seeded and incubated overnight at 37 °C. Cells were then treated with $LF_N$-Acr (2.5 µM) ± PA (20 nM) and incubated for 2 h at 37 °C. Cells were washed and stained with Hoechst 33342 and WGA555 in FluoroBrite DMEM, then imaged using confocal microscopy (60× objective) to detect fluorescence from Hoechst 33342 (blue nuclear marker; 405 nm excitation), WGA555 (yellow cytosolic marker; 568 nm excitation), and Cy5 (red; 647 nm excitation). Digital phase contrast (DPC) microscopy was also performed. Scale bars, 40 µm.

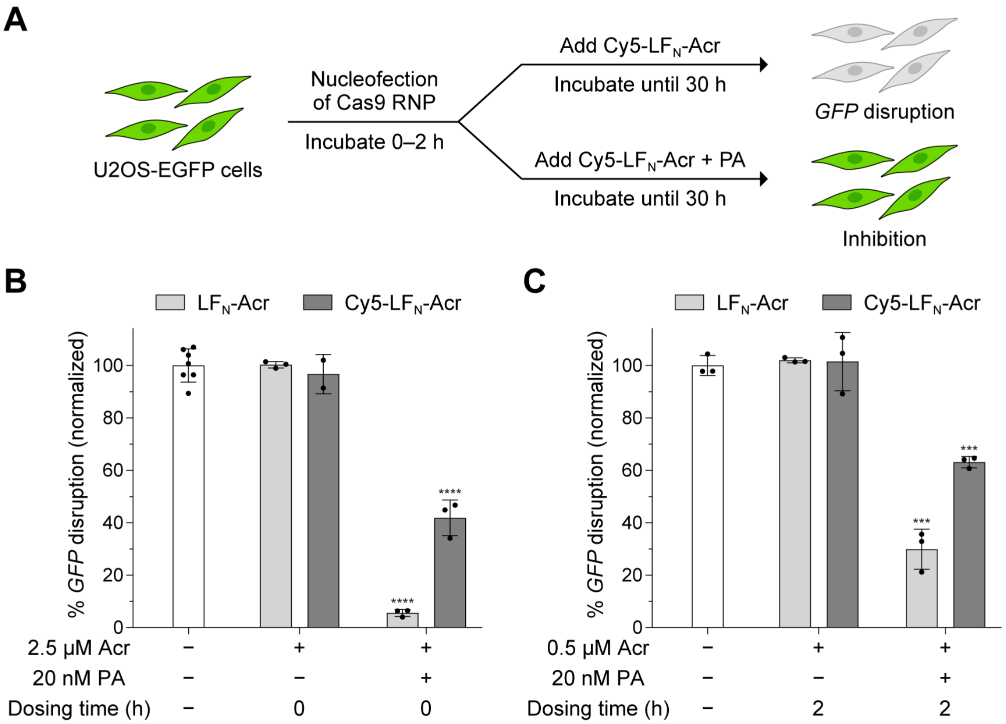


**Figure S33.** Effect of Cy5-$LF_N$-Acr delivery on *GFP* disruption. (A) A cartoon illustrating the experimental setup for monitoring Cy5-Acr delivery and activity in the *GFP*-disruption assay. Cy5-Acr delivery prevents *GFP* disruption. (B, C) Inhibition of Cas9 by Cy5-$LF_N$-Acr in the *GFP*-disruption assay. SpCas9-NLS RNP (20 pmol) targeting *GFP* was delivered via nucleofection into U2OS-EGFP.PEST cells. The cells were seeded at a density of 20,000 cells per well in 100 µL of culture medium, incubated at 37 °C, and treated with Cy5-$LF_N$-Acr (0.5–2.5 µM) ± PA (20 nM) at 0 h (panel B) or 2 h (panel C). After 30 h, live cells were stained with Hoechst 33342 and imaged using high-throughput confocal microscopy. Controls include nucleofection of Cas9 RNP (white bar) and nucleofection of Cas9 RNP followed by treatment with $LF_N$-Acr (0.5–2.5 µM) ± PA (20 nM) at 0 h (panel B) or 2 h (panel C). The values were normalized to Cas9 RNP and are the mean ± SD of three independent replicates. The significance of PA-mediated $LF_N$-Acr delivery was determined with an unpaired, two-tailed *t*-test versus Cas9 RNP, where *, **, ***, and **** refer to $P \leq 0.05$, $P \leq 0.01$, $P \leq 0.001$, and $P \leq 0.0001$, respectively.

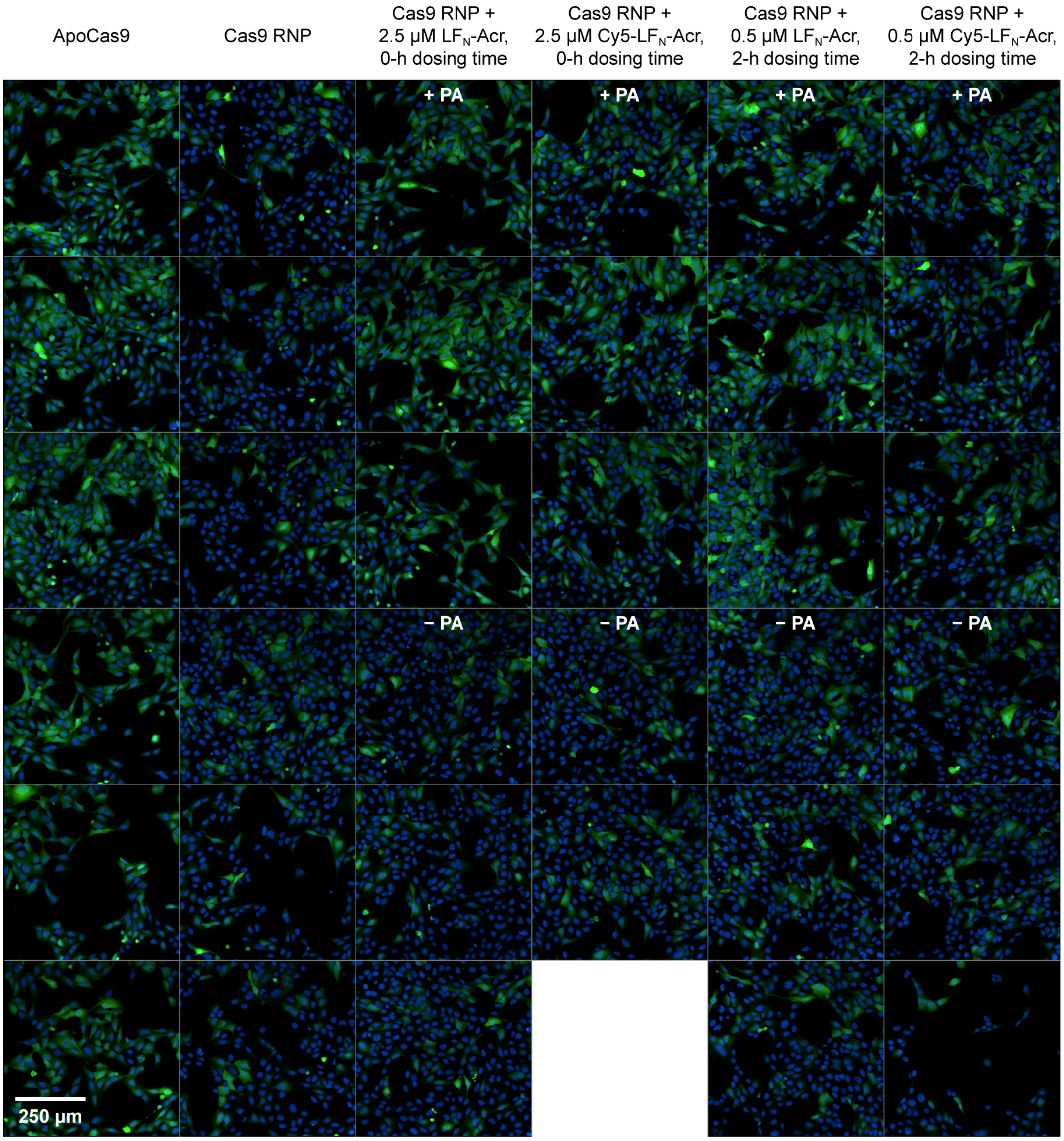


**Figure S34.** Representative fluorescence images from the *GFP*-disruption assay in Figure S33. SpCas9-NLS RNP (20 pmol) targeting *GFP* was delivered via nucleofection into U2OS-EGFP.PEST cells. The cells were seeded at a density of 20,000 cells per well in 100 μL of culture medium, incubated at 37 °C, and treated with Cy5-$LF_N$-Acr (0.5–2.5 μM) ± PA (20 nM) at 0 or 2 h. After 30 h, live cells were stained with Hoechst 33342 (blue, nucleus) and imaged using high-throughput confocal microscopy (20× objective). Controls include nucleofection of Cas9 RNP and nucleofection of Cas9 RNP followed by treatment with $LF_N$-Acr (0.5–2.5 μM) ± PA (20 nM) at 0 or 2 h.

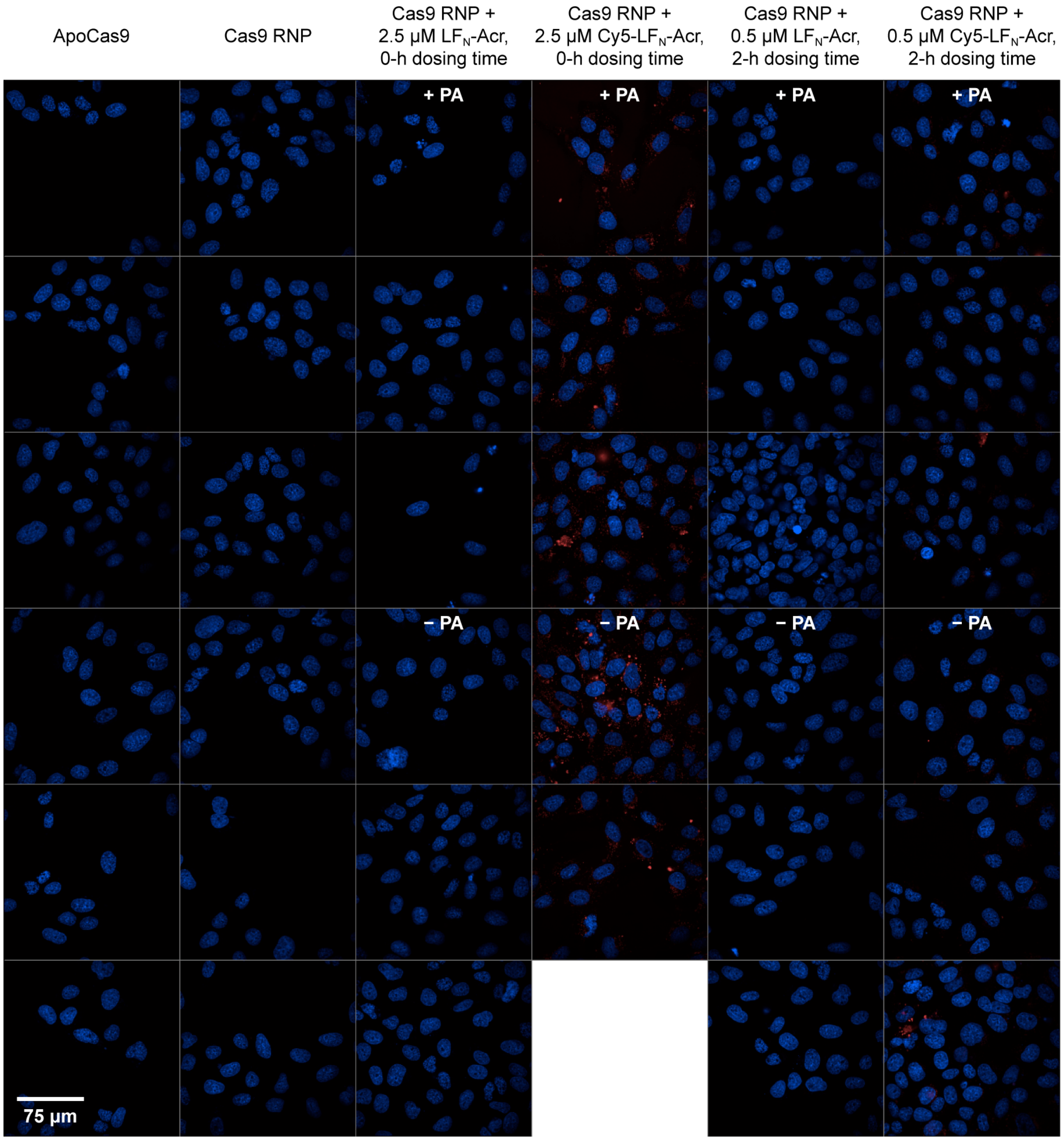


**Figure S35.** Direct visualization of Cy5-LF$_N$-Acr delivery in the *GFP*-disruption assay. SpCas9-NLS RNP (20 pmol) targeting *GFP* was delivered via nucleofection into U2OS-EGFP.PEST cells. The cells were seeded at a density of 20,000 cells per well in 100 µL of culture medium, incubated at 37 °C, and treated with Cy5-LF$_N$-Acr (0.5–2.5 µM) ± PA (20 nM) at 0 or 2 h. After 30 h, live cells were stained with Hoechst 33342 (blue, nucleus) and imaged using high-throughput confocal microscopy (60× objective). Controls include nucleofection of Cas9 RNP and nucleofection of Cas9 RNP followed by treatment with LF$_N$-Acr (0.5–2.5 µM) ± PA (20 nM) at 0 or 2 h. These are the same wells shown in Figure S34, imaged at higher magnification, including the Cy5 (red) channel and omitting the GFP channel for clarity.

### Plasmid Sequences

#### pAV13-MBP-TEV-4×NLS-Cys-AcrIIA4-2×NLS

The sequence that encodes 6×His-MBP-TEV-4×NLS-$G_4CG_4S$-AcrIIA4-GS-2×NLS is in uppercase, and its domains are underlined differently: 6×His-MBP tag, TEV site, 4×SV40 NLS, $G_4CG_4S$ linker, **AcrIIA4**, GS linker, 2×SV40 NLS.

cgtctccgggagctgcatgtgtcagaggttttcaccgtcatcaccgaaacgcgcgaggcagctgcggtaa
agctcatcagcgtggtcgtgaagcgattcacagatgtctgcctgttcatccgcgtccagctcgttgagtt
tctccagaagcgttaatgtctggcttctgataaagcgggccatgttaagggcggtttttcctgtttggt
cactgatgcctccgtgtaagggggatttctgttcatgggggtaatgataccgatgaaacgagagaggatg
ctcacgatacgggttactgatgatgaacatgcccggttactggaacgttgtgagggtaaacaactggcgg
tatggatgcggcgggaccagagaaaaatcactcagggtcaatgccagcgcttcgttaatacagatgtagg
tgttccacagggtagccagcagcatcctgcgatgcagatccggaacataatggtgcagggcgctgacttc
cgcgtttccagactttacgaaacacggaaaccgaagaccattcatgttgttgctcaggtcgcagacgttt
tgcagcagcagtcgcttcacgttcgctcgcgtatcggtgattcattctgctaaccagtaaggcaaccccg
ccagcctagccgggtcctcaacgacaggagcacgatcatgcgcacccgtggccaggacccaacgctgccc
gagatgcgccgcgtgcggctgctggagatggcggacgcgatggatatgttctgccaagggttggtttgcg
cattcacagttctccgcaagaattgattggctccaattcttggagtggtgaatccgttagcgaggtgccg
ccggcttccattcaggtcgaggtggcccggctccatgcaccgcgacgcaacgcggggaggcagacaaggt
atagggcggcgcctacaatccatgccaacccgttccatgtgctcgccgaggcggcataaatcgccgtgac
gatcagcggtccaatgatcgaagttaggctggtaagagccgcgagcgatccttgaagctgtccctgatgg
tcgtcatctacctgcctggacagcatggcctgcaacgcgggcatcccgatgccgccggaagcgagaagaa
tcataatggggaaggccatccagcctcgcgtcgcgaacgccagcaagacgtagcccagcgcgtcggccgc
catgccggcgataatggcctgcttctcgccgaaacgtttggtggcgggaccagtgacgaaggcttgagcg
agggcgtgcaagattccgaataccgcaagcgacaggccgatcatcgtcgcgctccagcgaaagcggtcct
cgccgaaaatgacccagagcgctgccggcacctgtcctacgagttgcatgataaagaagacagtcataag
tgcggcgacgatagtcatgccccgcgcccaccggaaggagctgactgggttgaaggctctcaagggcatc
ggtcgacgctctcccttatgcgactcctgcattaggaagcagcccagtagtaggttgaggccgttgagca
ccgccgccgcaaggaatggtgcatgcaaggagatggcgcccaacagtcccccggccacggggcctgccac
catacccacgccgaaacaagcgctcatgagcccgaagtggcgagcccgatcttccccatcggtgatgtcg
gcgatataggcgccagcaaccgcacctgtggcgccggtgatgccggccacgatgcgtccggcgtagagga
tcgagatctcgatcccgcgaaattaatacgactcactatagggagaccacaacggtttccctctagtgcc
ggctccggagagctctttaattaagcggccgccctgcaggactcgagttctagaaataattttgtttaac
tttaagaaggagatatacatATGAAATCTTCTCACCATCACCATCACCATGGTTCTTCTATGAAAATCGA
AGAAGGTAAACTGGTAATCTGGATTAACGGCGATAAAGGCTATAACGGTCTCGCTGAAGTCGGTAAGAAA
TTCGAGAAAGATACCGGAATTAAAGTCACCGTTGAGCATCCGGATAAACTGGAAGAGAAATTCCCACAGG
TTGCGGCAACTGGCGATGGCCCTGACATTATCTTCTGGGCACACGACCGCTTTGGTGGCTACGCTCAATC
TGGCCTGTTGGCTGAAATCACCCCGGACAAAGCGTTCCAGGACAAGCTGTATCCGTTTACCTGGGATGCC
GTACGTTACAACGGCAAGCTGATTGCTTACCCGATCGCTGTTGAAGCGTTATCGCTGATTTATAACAAAG
ATCTGCTGCCGAACCCGCCAAAAACCTGGGAAGAGATCCCGGCGCTGGATAAAGAACTGAAAGCGAAAGG
TAAGAGCGCGCTGATGTTCAACCTGCAAGAACCGTACTTCACCTGGCCGCTGATTGCTGCTGACGGGGGT

TATGCGTTCAAGTATGAAAACGGCAAGTACGACATTAAAGACGTGGGCGTGGATAACGCTGGCGCGAAAG
CGGGTCTGACCTTCCTGGTTGACCTGATTAAAAACAAACACATGAATGCAGACACCGATTACTCCATCGC
AGAAGCTGCCTTTAATAAAGGCGAAACAGCGATGACCATCAACGGCCCGTGGGCATGGTCCAACATCGAC
ACCAGCAAAGTGAATTATGGTGTAACGGTACTGCCGACCTTCAAGGGTCAACCATCCAAACCGTTCGTTG
GCGTGCTGAGCGCAGGTATTAACGCCGCCAGTCCGAACAAAGAGCTGGCAAAAGAGTTCCTCGAAAACTA
TCTGCTGACTGATGAAGGTCTGGAAGCGGTTAATAAAGACAAACCGCTGGGTGCCGTAGCGCTGAAGTCT
TACGAGGAAGAGTTGGCGAAAGATCCACGTATTGCCGCCACTATGGAAAACGCCCAGAAAGGTGAAATCA
TGCCGAACATCCCGCAGATGTCCGCTTTCTGGTATGCCGTGCGTACTGCGGTGATCAACGCCGCCAGCGG
TCGTCAGACTGTCGATGAAGCCCTGAAAGACGCGCAGACTAATTCGAGCTCGAACAACAACAACAATAAC
AATAACAACAACCTCGGGATCGAGGAAAACCTGTACTTCCAATCCAATGCCACTCCCAAGAAGAAGCGAA
AGGTGGGTGGGTCCCCAAAGAAGAAACGGAAAGTAGGCGGCTCCCCCAAAAAGAAGCGAAAAGTAGGGGG
TAGCCCCAAGAAGAAGCGGAAGGTAGGTATCCACGGAGTCCCAGCAGCTACCGGCGGTGGTGGTTGTGGT
GGGGGCGGCAGC**ATGAATATTAATGACTTAATTAGAGAAATCAAAAACAAAGATTACACAGTGAAATTGA**
**GTGGTACGGATAGCAATAGTATCACACAGCTAATTATTCGCGTTAATAATGATGGCAACGAGTATGTAAT**
**TTCTGAAAGTGAAAATGAATCAATCGTTGAAAAATTCATCTCTGCATTCAAAAACGGTTGGAATCAAGAA**
**TACGAGGATGAAGAAGAATTTTATAATGACATGCAAACAATCACCTTAAAAAGTGAGTTGAAC**GGGTCAC
CTAAGAAAAAACGAAAAGTTGAGGATCCTAAAAAGAAACGAAAAGTTGATACCGGTTAAtaacattggaa
gtggataacggatccgcgatcgcggcgcgccacctggtggccggccggtaccacgcgtgcgcgctgatcc
ggctgctaacaaagcccgaaaggaagctgagttggctgctgccaccgctgagcaataactagcataaccc
cttggggcctctaaacgggtcttgaggggttttttgctgaaaggaggaactatatccggatatccacagg
acgggtgtggtcgccatgatcgcgtagtcgatagtggctccaagtagcgaagcgagcaggactgggcggc
ggccaaagcggtcggacagtgctccgagaacgggtgcgcatagaaattgcatcaacgcatatagcgctag
cagcacgccatagtgactggcgatgctgtcggaatggacgatatcccgcaagaggcccggcagtaccggc
ataaccaagcctatgcctacagcatccagggtgacggtgccgaggatgacgatgagcgcattgttagatt
tcatacacggtgcctgactgcgttagcaatttaactgtgataaactaccgcattaaagcttatcgatgat
aagctgtcaaacatgagaattcttgaagacgaaagggcctcgtgatacgcctatttttataggttaatgt
catgataataatggtttcttagacgtcaggtggcacttttcggggaaatgtgcgcggaacccctatttgt
ttatttttctaaatacattcaaatatgtatccgctcatgagacaataaccctgataaatgcttcaataac
attgaaaaaggaagagtatgagtattcaacatttccgtgtcgcccttattcccttttttgcggcattttg
ccttcctgtttttgctcacccagaaacgctggtgaaagtaaaagatgctgaagatcagttgggtgcacga
gtgggttacatcgaactggatctcaacagcggtaagatccttgagagttttcgccccgaagaacgttttc
caatgatgagcacttttaaagttctgctatgtggcgcggtattatcccgtgttgacgccgggcaagagca
actcggtcgccgcatacactattctcagaatgacttggttgagtactcaccagtcacagaaaagcatctt
acggatggcatgacagtaagagaattatgcagtgctgccataaccatgagtgataacactgcggccaact
tacttctgacaacgatcggaggaccgaaggagctaaccgcttttttgcacaacatgggggatcatgtaac
tcgccttgatcgttgggaaccggagctgaatgaagccataccaaacgacgagcgtgacaccacgatgcct
gcagcaatggcaacaacgttgcgcaaactattaactggcgaactacttactctagcttcccggcaacaat
taatagactggatggaggcggataaagttgcaggaccacttctgcgctcggcccttccggctggctggtt
tattgctgataaatctggagccggtgagcgtgggtctcgcggtatcattgcagcactggggccagatggt
aagccctcccgtatcgtagttatctacacgacggggagtcaggcaactatggatgaacgaaatagacaga
tcgctgagataggtgcctcactgattaagcattggtaactgtcagaccaagtttactcatatatacttta

gattgatttaaaacttcatttttaatttaaaaggatctaggtgaagatcctttttgataatctcatgacc
aaaatcccttaacgtgagttttcgttccactgagcgtcagaccccgtagaaaagatcaaaggatcttctt
gagatcctttttttctgcgcgtaatctgctgcttgcaaacaaaaaaaccaccgctaccagcggtggtttg
tttgccggatcaagagctaccaactctttttccgaaggtaactggcttcagcagagcgcagataccaaat
actgtccttctagtgtagccgtagttaggccaccacttcaagaactctgtagcaccgcctacatacctcg
ctctgctaatcctgttaccagtggctgctgccagtggcgataagtcgtgtcttaccgggttggactcaag
acgatagttaccggataaggcgcagcggtcgggctgaacggggggttcgtgcacacagcccagcttggag
cgaacgacctacaccgaactgagatacctacagcgtgagctatgagaaagcgccacgcttcccgaaggga
gaaaggcggacaggtatccggtaagcggcagggtcggaacaggagagcgcacgagggagcttccaggggg
aaacgcctggtatctttatagtcctgtcgggtttcgccacctctgacttgagcgtcgatttttgtgatgc
tcgtcaggggggcggagcctatggaaaaacgccagcaacgcggcctttttacggttcctggccttttgct
ggccttttgctcacatgttctttcctgcgttatcccctgattctgtggataaccgtattaccgcctttga
gtgagctgataccgctcgccgcagccgaacgaccgagcgcagcgagtcagtgagcgaggaagcggaagag
cgcctgatgcggtattttctccttacgcatctgtgcggtatttcacaccgcaatggtgcactctcagtac
aatctgctctgatgccgcatagttaagccagtatacactccgctatcgctacgtgactgggtcatggctg
cgccccgacacccgccaacacccgctgacgcgccctgacgggcttgtctgctcccggcatccgcttacag
acaagctgtgac

#### Equations

$$\begin{matrix}\text{Normalized}\\ \text{Cas9 specificity}\\ \text{without Acr}\end{matrix} = \frac{\text{On/OT Cas9}}{\text{On/OT Cas9}} = 1 \quad \text{(S1)}$$

$$\begin{matrix}\text{Normalized}\\ \text{Cas9 specificity}\\ \text{with Acr}\end{matrix} = \frac{\text{On/OT Acr–inhibited Cas9}}{\text{On/OT Cas9}} = 1.10 - 1.41 \quad \text{(S2)}$$

$$\begin{matrix}\text{Increase in}\\ \text{Cas9 specificity}\\ \text{with Acr}\end{matrix} = \frac{\begin{matrix}\text{Normalized}\\ \text{Cas9 specificity}\\ \text{with Acr}\end{matrix} - \begin{matrix}\text{Normalized}\\ \text{Cas9 specificity}\\ \text{without Acr}\end{matrix}}{\begin{matrix}\text{Normalized}\\ \text{Cas9 specificity}\\ \text{without Acr}\end{matrix}} \times 100\% = 10 - 41\% \quad \text{(S3)}$$

$$\begin{matrix}\text{Total increase in}\\ \text{Cas9 specificity}\\ \text{with Acr}\end{matrix} = \begin{matrix}\text{Increase in}\\ \text{Cas9 specificity}\\ \text{with Acr (On/OT1)}\end{matrix} + \begin{matrix}\text{Increase in}\\ \text{Cas9 specificity}\\ \text{with Acr (On/OT2)}\end{matrix} + \begin{matrix}\text{Increase in}\\ \text{Cas9 specificity}\\ \text{with Acr (On/OT3)}\end{matrix} + \begin{matrix}\text{Increase in}\\ \text{Cas9 specificity}\\ \text{with Acr (On/OT4)}\end{matrix} = 90\% \quad \text{(S4)}$$

#### References

(1) Staahl, B. T.; Benekareddy, M.; Coulon-Bainier, C.; Banfal, A. A.; Floor, S. N.; Sabo, J. K.; Urnes, C.; Munares, G. A.; Ghosh, A.; Doudna, J. A. Efficient genome editing in the mouse brain by local delivery of engineered Cas9 ribonucleoprotein complexes. *Nat. Biotechnol.* **2017**, *35*, 431–434.

(2) Vera, A. O.; Truex, N. L.; Sreekanth, V.; Pentelute, B. L.; Choudhary, A.; Raines, R. T. Protective antigen–mediated delivery of an anti-CRISPR protein for precision genome editing. *Proc. Natl. Acad. Sci. U. S. A.* **2025**, *122*, e2426960122.

(3) Kapust, R. B.; Tözsér, J.; Fox, J. D.; Anderson, D. E.; Cherry, S.; Copeland, T. D.; Waugh, D. S. Tobacco etch virus protease: Mechanism of autolysis and rational design of stable mutants with wild-type catalytic proficiency. *Protein Eng.* **2001**, *14*, 993–1000.

(4) Cong, L.; Ran, F. A.; Cox, D.; Lin, S.; Barretto, R.; Habib, N.; Hsu, P. D.; Wu, X.; Jiang, W.; Marraffini, L. A. Multiplex genome engineering using CRISPR/Cas systems. *Science* **2013**, *339*, 819–823.

(5) Sreekanth, V.; Zhou, Q.; Kokkonda, P.; Bermudez-Cabrera, H. C.; Lim, D.; Law, B. K.; Holmes, B. R.; Chaudhary, S. K.; Pergu, R.; Leger, B. S.; Walker, J. A.; Gifford, D. K.; Sherwood, R. I.; Choudhary, A. Chemogenetic system demonstrates that Cas9 longevity impacts genome editing outcomes. *ACS Cent. Sci.* **2020**, *6*, 2228–2237.

(6) Reyon, D.; Tsai, S. Q.; Khayter, C.; Foden, J. A.; Sander, J. D.; Joung, J. K. FLASH assembly of TALENs for high-throughput genome editing. *Nat. Biotechnol.* **2012**, *30*, 460–465.

(7) Cowan, C. A.; Klimanskaya, I.; McMahon, J.; Atienza, J.; Witmyer, J.; Zucker, J. P.; Wang, S.; Morton, C. C.; McMahon, A. P.; Powers, D.; Melton, D. A. Derivation of embryonic stem-cell lines from human blastocysts. *N. Engl. J. Med.* **2004**, *350*, 1353–1356.

(8) Liu, Y.; Zou, R. S.; He, S.; Nihongaki, Y.; Li, X.; Razavi, S.; Wu, B.; Ha, T. Very fast CRISPR on demand. *Science* **2020**, *368*, 1265–1269.

(9) Pomerantsev, A. P.; McCall, R. M.; Chahoud, M.; Hepler, N. K.; Fattah, R.; Leppla, S. H. Genome engineering in *Bacillus anthracis* using tyrosine site-specific recombinases. *PLoS Biol.* **2017**, *12*, e0183346.

(10) Singh, Y.; Chaudhary, V. K.; Leppla, S. H. A deleted variant of *Bacillus anthracis* protective antigen is non-toxic and blocks anthrax toxin action in vivo. *J. Biol. Chem.* **1989**, *264*, 19103–19107.
